# An early Cambrian ecdysozoan with a terminal mouth but no anus

**DOI:** 10.1101/2020.09.04.283960

**Authors:** Yunhuan Liu, Huaqiao Zhang, Shuhai Xiao, Tiequan Shao, Baichuan Duan

## Abstract

The ecdysozoans are the most diverse animal group on Earth^1, 2^. Molecular clock studies indicate that the ecdysozoans may have diverged and diversified in the Ediacaran Period^3, 4^, but unambiguous ecdysozoan fossils first appear in the earliest Cambrian and are limited to cycloneuralians^5–7^. Here we report new material of the early Cambrian microscopic animal *Saccorhytus coronarius*, which was previously interpreted as a deuterostome^8^. *Saccorhytus coronarius* is reconstructed as a millimetric and ellipsoidal meiobenthic animal with a spinose armor and an anterior mouth but no anus. Purported pharyngeal gills in support of the deuterostome hypothesis^8^ are shown to be taphonomic artifacts. Phylogenetic analyses indicate that *Saccorhytus coronarius* belongs to the total-group Ecdysozoa, highlighting the morphological and ecological diversity of early Cambrian ecdysozoans.

---

The microscopic animal *Saccorhytus coronarius*^8^ was first reported from the early Cambrian Zhangjiagou Lagerstätte^9^ (*c.* 535 Ma^10^; Extended Data Fig. 1) of South China. It was interpreted as an anusless, meiobenthic deuterostome animal^8^. This interpretation was largely based on the presence of pharyngeal gills. If this interpretation is confirmed, *Saccorhytus coronarius* would fill a critical gap in the fossil record of the Deuterostomia and present an intriguing case of secondary loss of the anus, which is known to have occurred independently in several animal groups^11^. To further test the deuterostome interpretation and to shed fresh light on the morphology and ecology of this enigmatic animal, we analyzed new material of *Saccorhytus coronarius* from the Zhangjiagou Lagerstätte^9^.

As other fossils in the Zhangjiagou Lagerstätte^9^, the new fossils of *Saccorhytus coronarius* are secondarily phosphatized. Although most examined specimens are flattened or deformed, some are three dimensionally preserved with minimal deformation (Extended Data Fig. 2a–c; Supplementary Movies 1–3). Only the integument is preserved, and no traces of internal soft tissues were discovered in all specimens examined by SRXTM^12^ (Extended Data Figs. 2, 3; Supplementary Movies 1–9). The integument is preserved as a two-layered structure (Fig. 1d; Extended Data Figs. 2d, 3c, 4a, 5e), which may represent two sub-layers of the cuticle^8^ that were thickened by excessive phosphate coating (Extended Data Fig. 3c). Alternatively, the two layers may represent phosphatic coatings on two separate organic sheets, as evidenced by the two detached or separate layers in some specimens (Fig. 1d). No cilium insertion sites were observed on the integument, even in high-magnification SEM images. A chevron pattern, which may be a diagenetic artifact, is present in some integument (Fig. 2h; Extended Data Figs. 3h, 4e, 5e, 6f, 7a, c).

**Figure 1.**
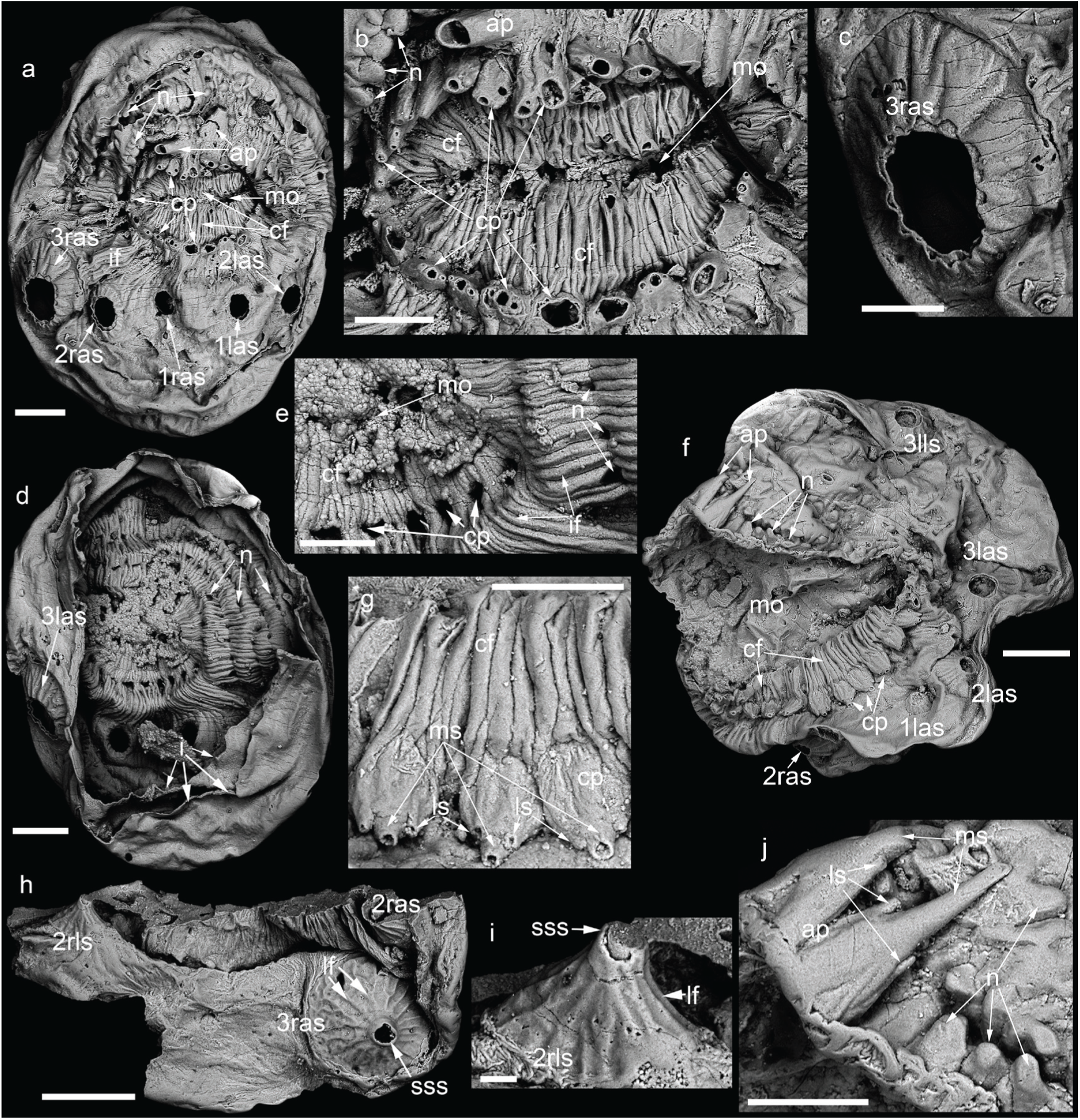
*Saccorhytus coronarius*. **a**–**e**, UMCU2016009; **a**, anterior view; **b**, detail of mouth; **c**, detail of broken spinose sclerite with jagged edge exposing two integument layers; **d**, opposite side of **a**, with posterior integument missing, showing inner view of mouth and detached integument layers; **e**, detail of inner side of mouth, showing hollow nature of circumoral protrusions; **f**, **g**, **j**, UMCU2019016; **f**, anterior left view; **g**, detail of circumoral protrusions; **j**, detail of anterior protrusions; **h**, **i**, UMCU2018011; **h**, a fragment; **i**, detail of spinose sclerite, showing remnant of apical spine. Abbreviations: ap, anterior protrusion; bss, base of spinose sclerite; cf, circumoral fold; ch, chevron; cp, circumoral protrusion; i, integument; if, integument fold; lf, longitudinal fold; ls, lateral spine; mo, mouth opening; ms, main spine; n, node; (1-3)l/ras,1st to 3rd left/right antero-ventral spinose sclerite; (1-5)l/rls, 1st to 5th left/right dorso-lateral spinose sclerite; sp, (posterior) spine; sss, apical spine of spinose sclerite. Scale bar: 200 μm (**a**, **d**, **f**, **h**), 100 μm (**b**, **c**, **e**, **g**, **j**), 20 μm (**i**).

**Figure 2.**
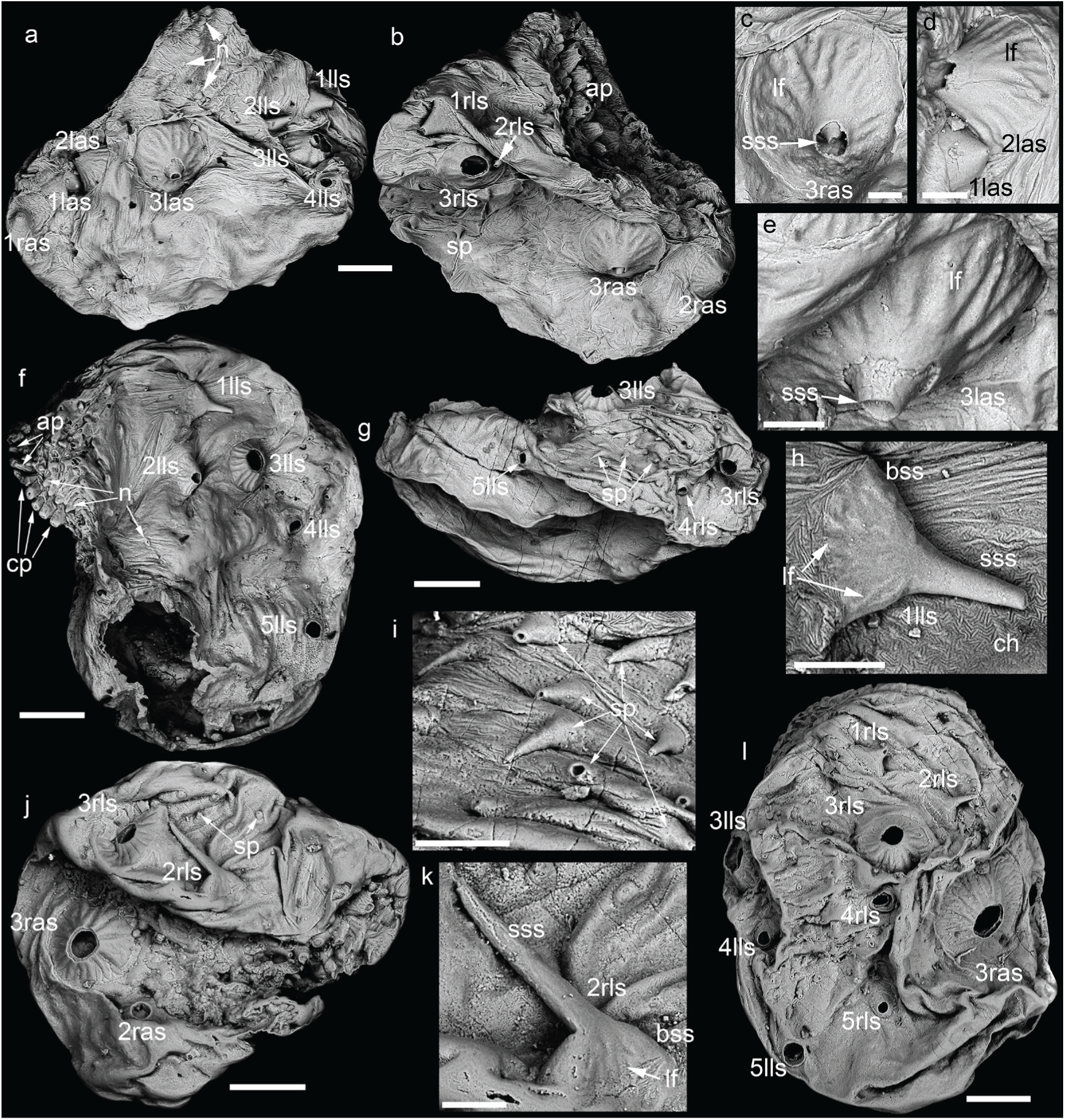
*Saccorhytus coronarius*. **a**–**e**, UMCU2020021; **a**, left view; **b**, right view; **c**–**e**, detail of spinose sclerites tilted ∼40° from **a** and **b**, showing remnant of apical spine; **f**, **h**, UMCU2016008; **f**, left view; **h**, detail of spinose sclerite adpressed against body wall; **g**, **i**, UMCU2016010; **g**, posterior view; **i**, detail of posterior spines adpressed against body wall; **j**, **k**, UMCU2018012; **j**, right view; **k**, detail of spinose sclerite adpressed against body wall; **l**, UMCU2019018, posterior right view. Scale bar: 200 μm (**a**, **b**, **f**, **g**, **j**, **l**), 60 μm (**c**–**e**, **h**, **i**, **k**). See Fig. 1 for abbreviations.

*Saccorhytus coronarius* is ellipsoidal in overall shape, with a terminal mouth (Fig. 1a, f; Extended Data Figs. 2a, d, 3d, e, 4a, h), which is regarded as the anterior end because in most living bilaterians the mouth is located at or near the anterior end. When the mouth is closed, it forms a flattened slit (Fig. 1b; Extended Data Figs. 7f, 8a). From the anterior view, the flattened mouth divides the body into two halves, with one half slightly wider than the other (Extended Data Fig. 2a, b; Supplementary Movie 3); the wider half is regarded as the ventral side because it helps to balance the animal for hydrodynamic stability. When the mouth is partially open, it has V-shaped left and right corners (Fig. 1f; Extended Data Figs. 2d, 3d, 4h, 5c, 6a). When fully open, the mouth is nearly circular (Extended Data Figs. 6d, 8b). Unlike the prominent mouth at the anterior end, no traces of an anus are found on any specimen, confirming the absence of an anus in *Saccorhytus coronarius*^8^.

The mouth is surrounded by radially arranged circumoral folds of the integument (Fig. 1b, e, f; Extended Data Figs. 2d, f, g, 3d, 4a, c, h, 6a, d, 8a). The folds appear flexible, and thus were able to control the opening and closure of the mouth. The folds are surrounded by a circlet of about 30 circumoral protrusions that are in close contact with each other (Fig. 1f; Extended Data Figs. 2d, 4h). The circumoral protrusions are internally hollow (Fig. 1f), largest on the dorsal and ventral sides, but decrease in size toward the left and right sides (Fig. 1f; Extended Data Figs. 2d, f, 4h, 5c), thus exhibiting biradial symmetry. Each circumoral protrusion is flattened and tridented, with a larger main spine flanked by two smaller lateral spines (Fig. 1g; Extended Data Figs. 2f, g, 4g, 6c, 8e). At the antero-dorsal margin of the circumoral protrusions, there is an array of anterior protrusions. The number of anterior protrusions varies from one (Extended Data Figs. 3d, 5a), two (Fig. 1a, f; Extended Data Figs. 4a, h, 6a, d, 7f, 8b, c), three (Extended Data Fig. 2d), four (Extended Data Figs. 4c, 5c), to five (Extended Data Figs. 2a, 7d), possibly representing intraspecific or ontogenetic variations. They are arranged in one row if fewer than five, or in two rows if five. The anterior protrusions are also flattened and tridented, with a prominently longer and larger main spine flanked by two lateral spines or denticles (Fig. 1j; Extended Data Figs. 4c, 6a, 8a, e). The circumoral and anterior protrusions have similar morphology, and the only difference is that the latter are larger and longer. Because the anterior protrusions occur only on the dorsal side, they impose a dorsal-ventral polarity and render a true bilateral symmetry to the animal.

Further posterior to the circumoral and anterior protrusions, there are integument folds that are arranged anteriorly-posteriorly but appear radial in anterior view (Fig. 1a, e; Extended Data Figs. 2d, 4a, c, 5c, 6a, d, 7b, 8a), as well as nodes or tubercles on the dorsal and lateral sides (Fig. 1a, f; Extended Data Figs. 2d, e, 4a, c, h, 5c, 6a, d, 7d). These nodes are arranged in crescent half-circlets or sets parallel to the circumoral protrusions. The first set of nodes are aligned with the anterior protrusions and extend laterally (Fig. 1j; Extended Data Figs. 4a, c, 6a, d, 7d); the next two or three sets extend continuously to both lateral sides, whereas the remaining two to three sets are more sparsely and less regularly arranged (Extended Data Figs. 4a, c, 7d), and there are sometimes single nodes positioned dorsally (Extended Data Fig. 7d).

Further posteriorward, there are two sets of spinose sclerites, one set antero-ventrally and the other dorso-laterally. The antero-ventral spinose sclerites consist of three pairs that are bilaterally symmetrically arranged (Fig. 1a, f; Extended Data Figs. 2d, 4a, i, 5c, d, 6d, 8b). The medial (1st) pair are the smallest, whereas the lateral-most (3rd) pair are the largest (Supplementary Table 1). Each spinose sclerite has an expanded conical base, and the completely preserved sclerites have a long distal spine with a closed tip. Most sclerites are broken at the juncture between the conical base and the apical spine, leaving behind jagged edges of the conical base and exposing the two integument layers (Fig. 2j, l; Extended Data Figs. 4i, j, 8b), but some are broken from the base or middle of the conical base (Fig. 1c; Extended Data Figs. 2d, 5e, 7f, 8b), and others are broken from the apical spine, leaving the proximal part of the distal spine still attached to the conical base (Figs. 1h, 2c, e). The conical base is ornamented with longitudinal folds or ridges (Figs. 1c, h, 2c–e; Extended Data Figs. 5a, e, 8b), and it is radially symmetrical or slightly lopsided (with the center offset slightly medially; Fig. 1c; Extended Data Figs. 5a, 8b). The dorso-lateral set consists of up to five pairs of spinose sclerites arranged dorso-laterally in a bilaterally symmetric fashion (Fig. 2f, l; Extended Data Figs. 4b, j, 5c, d, 6e, 7b, e, g). The third pair are always present (Figs. 1f, 2a, b, f, j, l; Extended Data Figs. 2b, 3d, 4b, h, j, 5b, c, 6e, h, 7b, g, 8f) and are the largest in size (Supplementary Table 1), whereas the other spinose sclerites may be absent (Fig. 1f; Extended Data Fig. 2e), probably reflecting intraspecific or ontogenetic variations. These spinose sclerites also have a conical base with longitudinal folds, which are most prominent on the third pair (Figs. 1i, 2f, h, j; Extended Data Figs. 4e, 6f, g, i, 7a, c), and a long distal spine with a closed tip (Fig. 2h, k; Extended Data Fig. 4k). Undeformed conical bases always have a circular footprint (e.g., Extended Data Figs. 4e, 7h), although they can be flattened when adpressed against the body wall (Fig. 2h, k; Extended Data Figs. 4j, k, 6f, g, i, 7a), suggesting a degree of flexibility. The apical spines are typically broken from the base (Fig. 2f, l; Extended Data Figs. 5g, 6f, g, i, 7c), but they can be partially broken (Fig. 1i) or nearly completely preserved (Fig. 2h, k; Extended Data Figs. 3i, 4k, 7a, 8d). The completely preserved spines are always adpressed on the body wall (thus avoiding being abraded) and they are about three to four times the height of the base. Our observation suggests that the 1st-3rd pairs of antero-ventral spinose sclerites are equivalent to 1st-3rd pairs of body cones, and the 3rd pair of dorso-lateral spinose sclerites are equivalent to the 4th pair of body cones of Han et al.^8^, who interpreted these structures as gill slits.

On the posterior side of the body, there are many short and slender spines with a sharp tip (Fig. 2g, i; Extended Data Figs. 2e, 3h, 4b, 6e). They may extend to the fifth dorso-lateral spinose sclerites (Fig. 2f; Extended Data Figs. 4b, 6e). They are distributed randomly, separate from each other, and directed postero-ventrally. They vary in size, with the ventral ones slightly smaller, but they are generally much smaller than the spinose sclerites described above (Supplementary Table 1) and they are typically broken from the base (Extended Data Figs. 4b, 5d, 6e). Again, the more completely preserved posterior spines tend to be those that are adpressed against the integument (Fig. 2g, i) and thus are not abraded during fossil preservation or preparation.

*Saccorhytus* is reconstructed as a sac-like animal with a spinose armor and an anterior mouth but no anus (Fig. 3a–c; Supplementary Movie 10). Its small body size indicates a meiobenthic life style in a physical niche with low Reynolds number^13^. The non-ciliated integument implies that it did not have cilia for locomotion or feeding. It also lacked setae or paired appendages, thus was unable to creep or crawl. Locomotion by hydrostatic skeletons seems unlikely. *Saccorhytus* may be a free epibenthic or interstitial animal. The circumoral protrusions, anterior protrusions, spinose sclerites, and posterior spines may have functioned as sensory and/or defense organs, whereas the mouth appears to be the only body opening for feeding and excretion.

**Figure 3.**
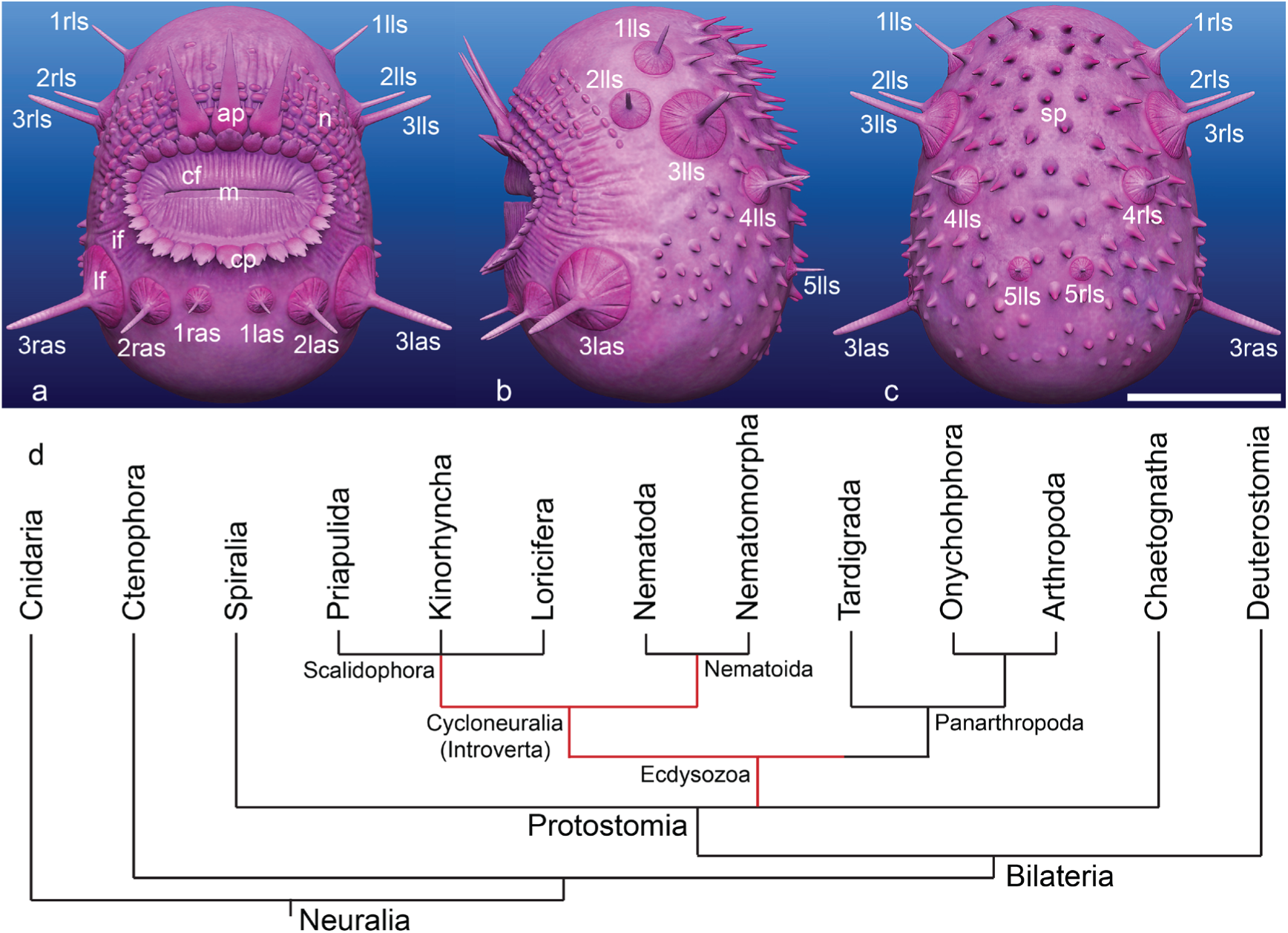
Reconstruction and phylogenetic interpretation of *Saccorhytus coronarius*. **a**–**c**, reconstructions showing anterior (**a**), left (**b**), and posterior (**c**) views, apical spines in 1l/ras, 2l/ras, and 3-5l/rls are inferred from their similarity to 3l/ras and 1-2l/rls where apical spines are directly observed; scale bar 500 μm; see Fig. 1 for abbreviations; **d**, simplified phylogeny of Neuralia (based on ref^14^ and ref^25^) and possible phylogenetic positions of *Saccorhytus* (red line).

The bilateral symmetry with both anterior-posterior and dorsal-ventral axes suggests that *Saccorhytus* should fall within the Bilateria^14^. *Saccorhytus* lacks an anus, but without a phylogenetic context, this condition may represent either primitive absence or secondary loss, because a through gut with a terminal anus may have evolved independently in different bilaterian lineages^15^, and secondary loss of the anus has also occurred several times in at least nine bilaterian groups^11^. The previously proposed deuterostome hypothesis^8^ was based on similarities between the body cones of *Saccorhytus* and the body openings (interpreted as pharyngeal gills) of vetulicolians^16^ and vetulocystids^17^. Better preserved specimens illustrated here (Figs. 1h, 2c, e), however, show that the so-called body cones in *Saccorhytus* are in fact remnants of broken spinose sclerites. *Saccorhytus* thus lacks homologs of pharyngeal gills^14, 18^, and as such should not be considered as a deuterostome. Indeed, the revised anatomical observations, when incorporated into the morphological character matrix of Han et al.^8^ and subjected to phylogenetic analyses, resolved *Saccorhytus* as the sister-group of the Acoelomorpha, outside the clade that includes protostomes and deuterostomes (Extended Data Fig. 9a).

As the data matrix of Han et al.^8^ has limited character and taxonomic coverage, we conducted new cladistic analyses based on a comprehensive morphological matrix published by Peterson and Eernisse^19^ to more thoroughly investigate the phylogenetic affinity of *Saccorhytus*. Various coding schemes and multiple cladistic methods were explored to test the sensitivity of the phylogenetic interpretation of *Saccorhytus*. In the most conservative coding scheme, *Saccorhytus* was resolved to be a total-group ecdysozoan (Extended Data Fig. 9b, a PAUP tree) or a stem-group nematoid (Extended Data Fig. 9c, a MrBayes tree), both with relatively low statistical support. The ecdysozoan affinity of *Saccorhytus* is supported by the radially arranged circumoral folds that are regarded as an autapomorphy of the total-group Ecdysozoa^20^, as well as a terminal mouth and the absence of cilia.

In a separate coding scheme, we consider the possibility that the two-layered integument of *Saccorhytus* (Fig. 1d; Extended Data Fig. 3c) represents evidence for an outer epicuticle and an inner procuticle^2, 14, 21^. This possibility is consistent with the consideration that recalcitrant tissues such as cuticle are often preferentially phosphatized in the Zhangjiagou Lagerstätte and other taphonomically similar Cambrian assemblages^22^. A cladistic analysis, assuming the presence of cuticle and ecdysis in *Saccorhytus*, gave the same results as far as *Saccorhytus* is concerned, but with higher statistical support (Extended Data Fig. 10a, b).

We note that the circumoral and anterior protrusions of *Saccorhytus* are internally hollow and may have contained soft tissues when alive. These protrusions are thus similar to the scalids or spinoscalids of the scalidophorans that contain soft tissue internally^23^; although arrow worms (Chaetognatha) also have internally hollow perioral spines or grasping spines, these spines do not form radially arranged circumoral circlets^2, 14^. In addition, the spinose sclerites and posterior spines are morphologically similar to the spinose armors of the coeval scalidophorans^6, 7^ in their general shape and internally hollow nature. Considering these similarities, additional cladistic analyses, assuming homology between the circumoral/anterior protrusions and the scalids/spinoscalids, resolved *Saccorhytus* as a stem-group scalidophoran (Extended Data Fig. 10c, d). We regard the stem-group scalidophoran interpretation as tentative, because *Saccorhytus* does not have a differentiated introvert, a feature characteristic of the Cycloneuralia and especially the more restrictive group of Scalidophora^14, 24^.

To summarize, *Saccorhytus* does not have pharyngeal gills and is not a deuterostome animal. Sequential cladistic analyses, starting with more conservative character coding and progressively testing more ambiguous characters, resolved *Saccorhytus* as a total-group ecdysozoan and possibly a stem-group scalidophoran (Fig. 3d). A crown-group panarthropod interpretation is unlikely due to lack of appendages in *Saccorhytus*. Together with co-occurring cycloneuralians from the Zhangjiagou Lagerstätte^5, 7^ and equivalent strata in South China^6^, *Saccorhytus* testifies a remarkable morphological and ecological diversity of early Cambrian ecdysozoans. With a better phylogenetic constraint, *Saccorhytus* presents another ecdysozoan case of secondary loss of the anus in addition to the Onychophora^11^.

## Supporting information

Liuetal_Movie 1

Liuetal_Movie 2

Liuetal_Movie 3

Liuetal_Movie 4

Liuetal_Movie 5

Liuetal_Movie 6

Liuetal_Movie 7

Liuetal_Movie 8

Liuetal_Movie 9

Liuetal_Movie 10

## Online content

Any methods, additional references, source data, extended data, supplementary information, acknowledgements; details of author contributions and competing interests; and statements of data and code availability are available at https://doi.org/XXXX.

## Methods

Rock samples were macerated in diluted acetic acid (10%). Microfossils were handpicked from residues under a binocular microscope. Specimens were glued on pin-type aluminum stubs for observation under a LEO 1530VP field-emission environmental scanning electron microscope.

Three specimens were scanned using SRXTM^12^ at the X02DA TOMCAT beam line, Swiss Light Source, Paul Scherrer Institute, Villigen, Switzerland. The SRXTM data were obtained using a 20 µm LuAg:Ce scintillator, a 20× objective lens, at an energy level of 14 KeV and an exposure time of 200 ms, as a series of 1501 equiangular projections while the samples were rotated through 180° within the beam. The projections were post-processed and rearranged into flat- and dark-field-corrected sinograms, and reconstruction was performed on a 60-core Linux PC farm, using a highly optimized routine based on the Fourier transform method and a regridding procedure^26^. The SRXTM data were analysed using the AVIZO8.0 (ThermoFisher Scientific) software package.

Phylogenetic analyses were carried out using the software PAUP4^27^, MrBayes3.2.7a^28^, and TNT1.1^29^. The most parsimonious trees and 50% majority rule trees were generated using PAUP4 (heuristic search; addition sequence = random, number of replicates = 100, hold one tree at each step; swapping algorithm = TBR). Bootstrap support values were calculated also using PAUP4 (full heuristic search, number of replicates = 10; addition sequence = simple, hold one tree at each step; swapping algorithm = TBR), whereas Bremer support values were calculated using TNT1.1 (saving trees of up to extra 10 steps longer). The Bayesian 50% majority trees were generated in MrBayes3.2.7a using the MK + informative model (which accounts for only parsimony-informative characters having been scored, and ascertainment bias) with equal rate variation (MKI + equal).

## Data availability

The data that support the findings of this study are available in the paper and its Supplementary Information, or from the corresponding authors upon reasonable request. All specimens illustrated in this paper are deposited in University Museum of Chang’an University (accession numbers UMCU2014001–2014005, 2016006–2016010, 2018011–2018015, 2019016–2019020, and 2020021–2020025). SRXTM data are archived in Morphosource.

## Acknowledgements

Supported by National Natural Science Foundation of China (nos. 41872014 and 41572007), Strategic Priority Research Program of CAS (no. XDB26000000), State Key Laboratory of Palaeobiology and Stratigraphy, Nanjing Institute of Geology and Paleontology, CAS (nos. 193123 and 20191104), Youth Innovation Promotion Association, CAS (no. 2016283). S.X. was supported by the U.S. National Science Foundation (EAR-2021207). We thank Dinghua Yang for assistance in artistic reconstructions, and Philip C. J. Donoghue for help in SRXTM scan in Swiss Light Source.

## Author contributions

H.Z. designed the research; Y.L. and T.S. obtained the fossils; H.Z. did the SEM work and performed the SRXTM scans; B.D. analyzed the SRXTM data; H.Z. and S.X. developed the interpretation and prepared the manuscript.

## Competing interests

The authors declare no competing interests.

## Extended Data

**Extended Data Figure 1.**
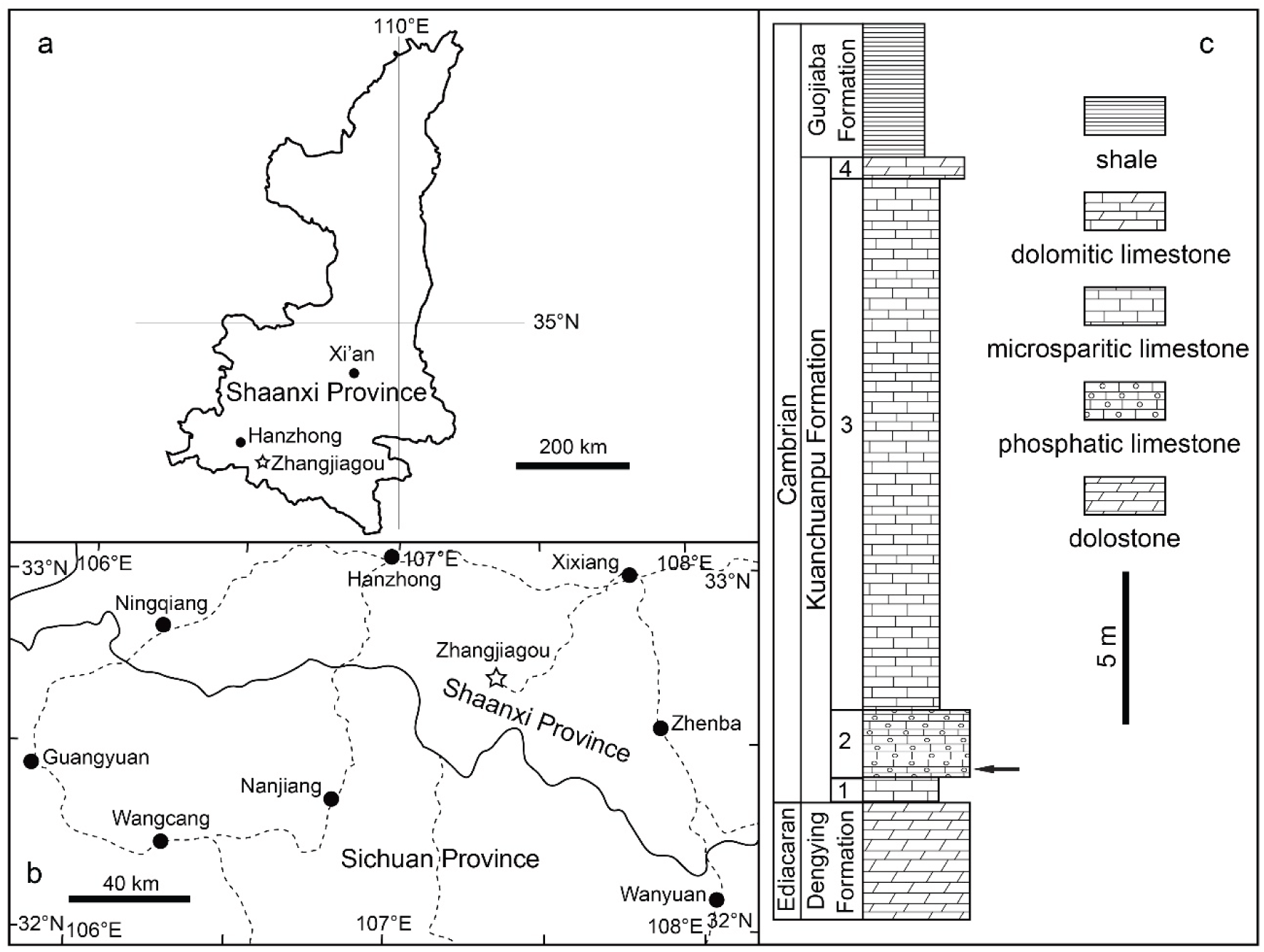
Location map and stratigraphic column of the Zhangjiagou section in southern Shaanxi Province, South China. **a**, map of Shaanxi Province, with star marking Zhangjiagou section where fossils were collected; **b**, detailed map of southern Shaanxi Province showing fossil locality (star); **c**, stratigraphic column of Zhangjiagou section showing key horizon (arrow) where fossils were collected.

**Extended Data Figure 2.**
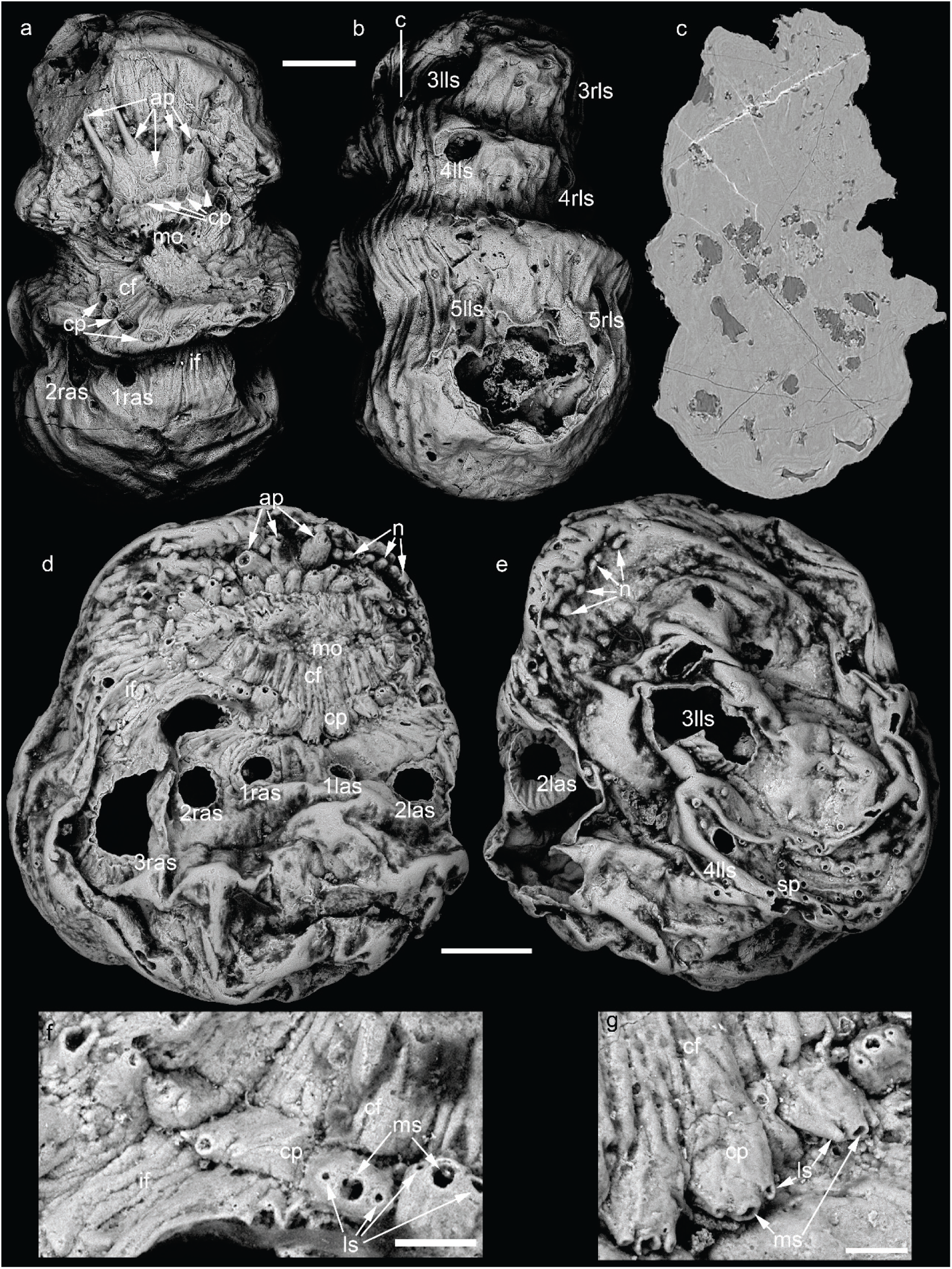
*Saccorhytus coronarius*. **a**–**c**, UMCU2014005, with five anterior protrusions; **a**, anterior view; **b**, posterior view; **c**, SRXTM image, transverse section marked in **b**; **d**–**g**, UMCU2014001, with three anterior protrusions; **d**, anterior view; **e**, posterior view; **f**, **g**, detail of circumoral protrusions. Scale bar: 200 μm (**a**–**e**), 50 μm (**f**), 40 μm (**g**). See Fig. 1 for abbreviations.

**Extended Data Figure 3.**
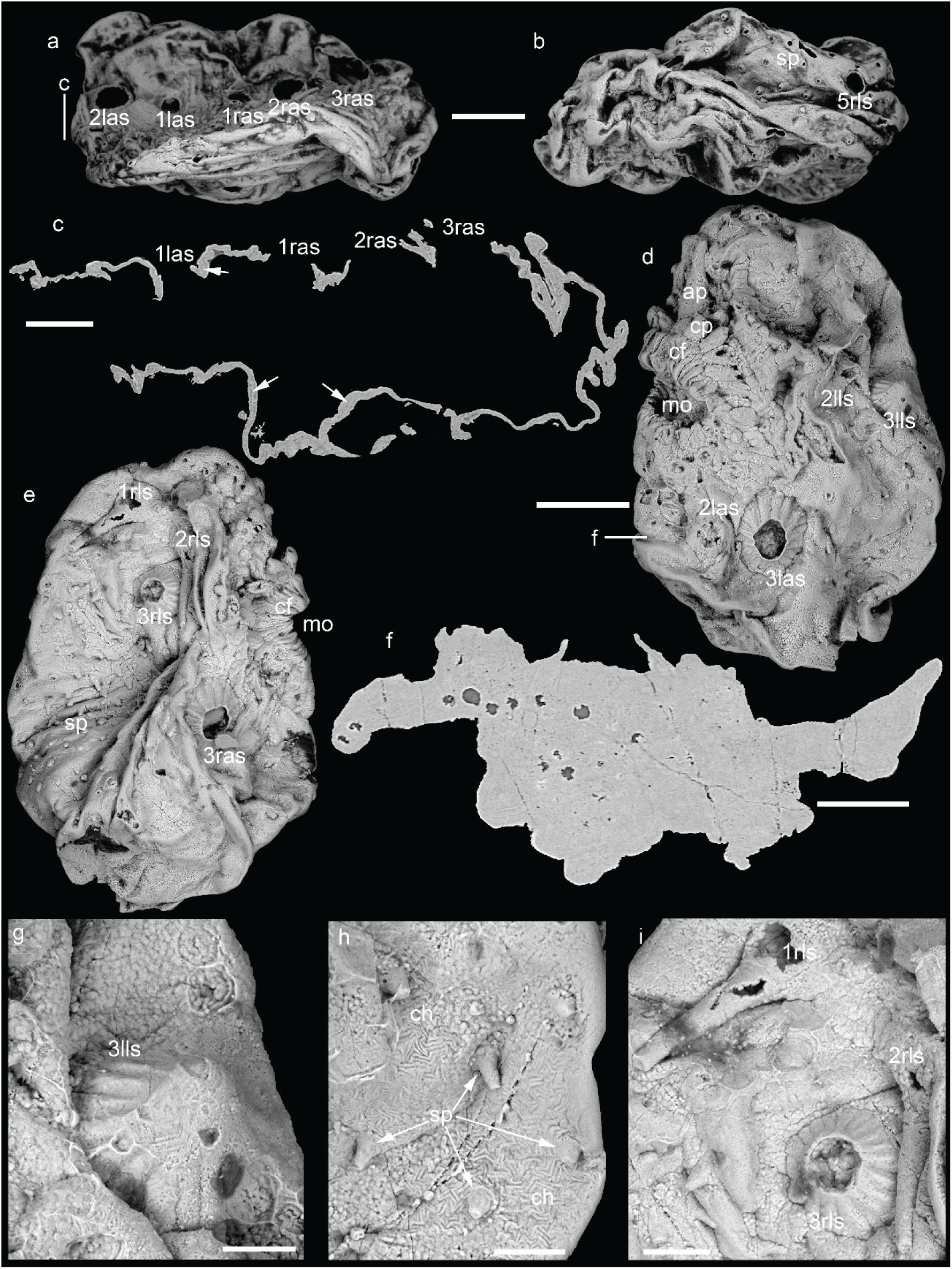
*Saccorhytus coronarius*. **a**–**c**, UMCU2014001, same specimen as in Extended Data Fig. 2d; **a**, dorso-anterior view; **b**, ventral view; **c**, SRXTM image, longitudinal section marked in **a**, with arrows marking boundary between two layers of integument; **d**–**i**, UMCU2014002; **d**, left view; **e**, right view; **f**, SRXTM image, tangential coronal section marked in **d**; **g**, close-up of spinose sclerite in central right of **d**; **h**, detail of posterior spines and chevron patterns in lower right of **d**; **i**, detail of spinose sclerites in upper central of **e**. Scale bar: 200 μm (**a**, **b**, **d**, **e**); 100 μm (**c**, **f**); 40 μm (**g**, **h**), 60 μm (**i**). See Fig. 1 for abbreviations.

**Extended Data Figure 4.**
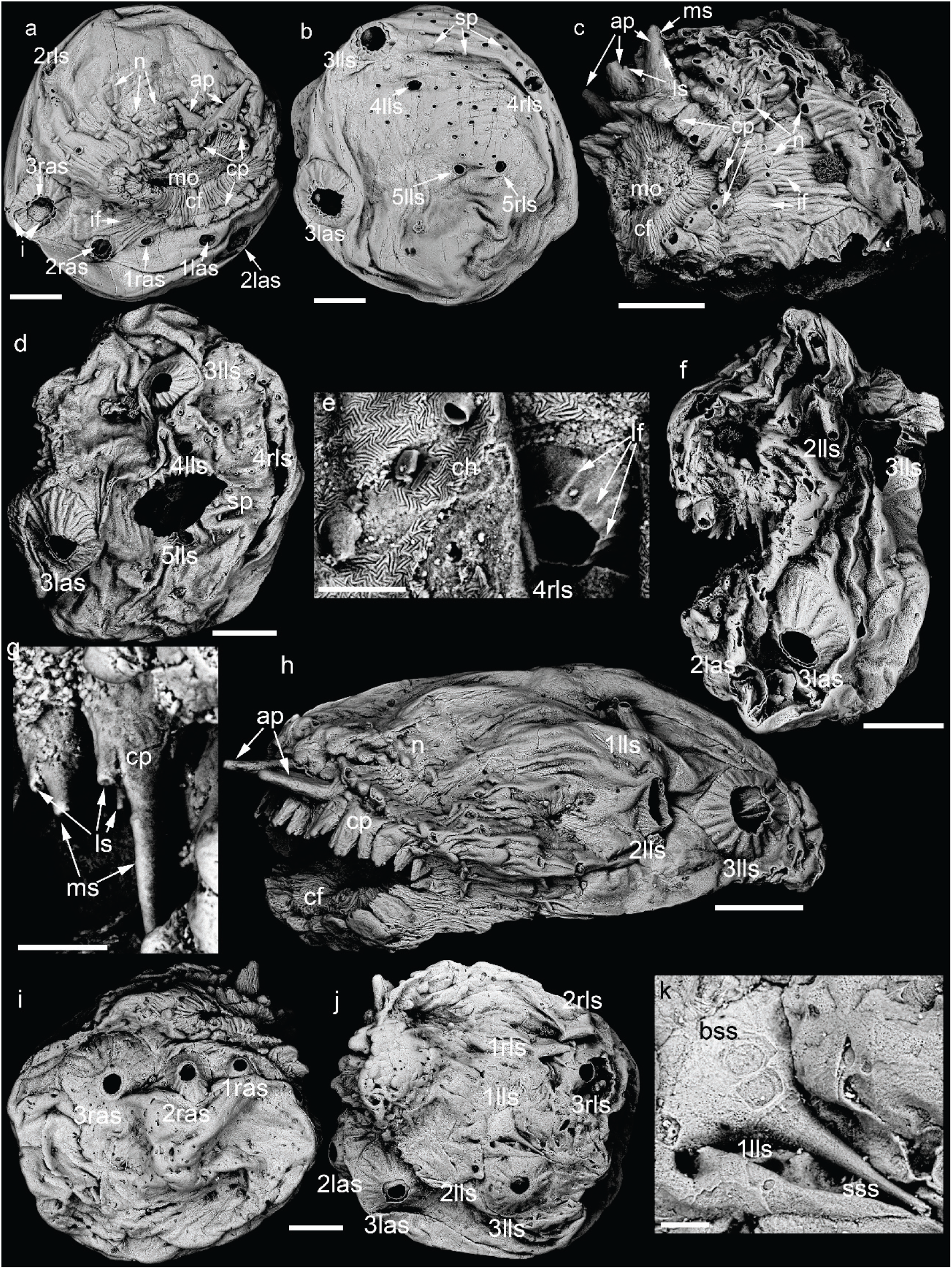
*Saccorhytus coronarius*. **a**, **b**, UMCU2019017, with two anterior protrusions; **a**, anterior view; **b**, posterior view; **c**, UMCU2016006, with four anterior protrusions, antero-left view; **d**, **e**, UMCU2020022; **d**, left view; **e**, detail of spinose sclerite and chevron pattern in central right of **d**; **f**, **g**, UMCU2020023; **f**, left view; **g**, detail of circumoral protrusions in central left of **f**; **h**, UMCU2018013, with two anterior protrusions, antero-left view; **i**–**k**, UMCU2020024; **i**, right ventral view; **j**, left dorsal view; **k**, detail of spinose sclerite in central of **j**. Scale bar: 200 μm (**a**–**d**, **f**, **h**–**j**), 40 μm (**e**, **g**, **k**). See Fig. 1 for abbreviations.

**Extended Data Figure 5.**
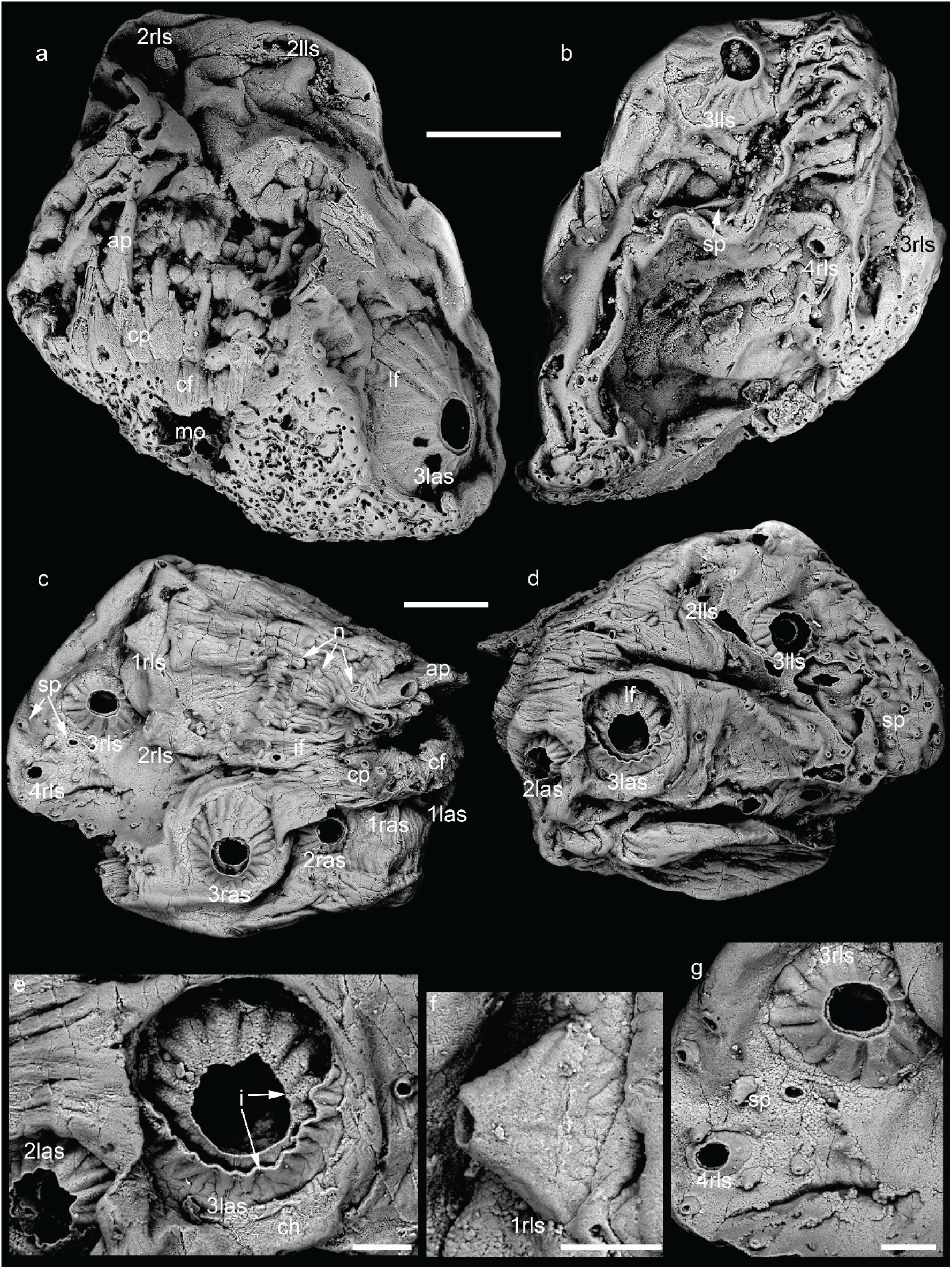
*Saccorhytus coronarius*. **a**, **b**, UMCU2014004, with only one anterior protrusion; **a**, anterior dorsal view; **b**, posterior view; **c**–**g**, UMCU2018014, with four anterior protrusions; **c**, right view; **d**, left view; **e**–**g**, detail of spinose sclerites and posterior spines in central left of **d**, upper left of **c**, and central left of **c**, respectively. Scale bar: 200 μm (**a**–**d**), 60 μm (**e**–**g**). See Fig. 1 for abbreviations.

**Extended Data Figure 6.**
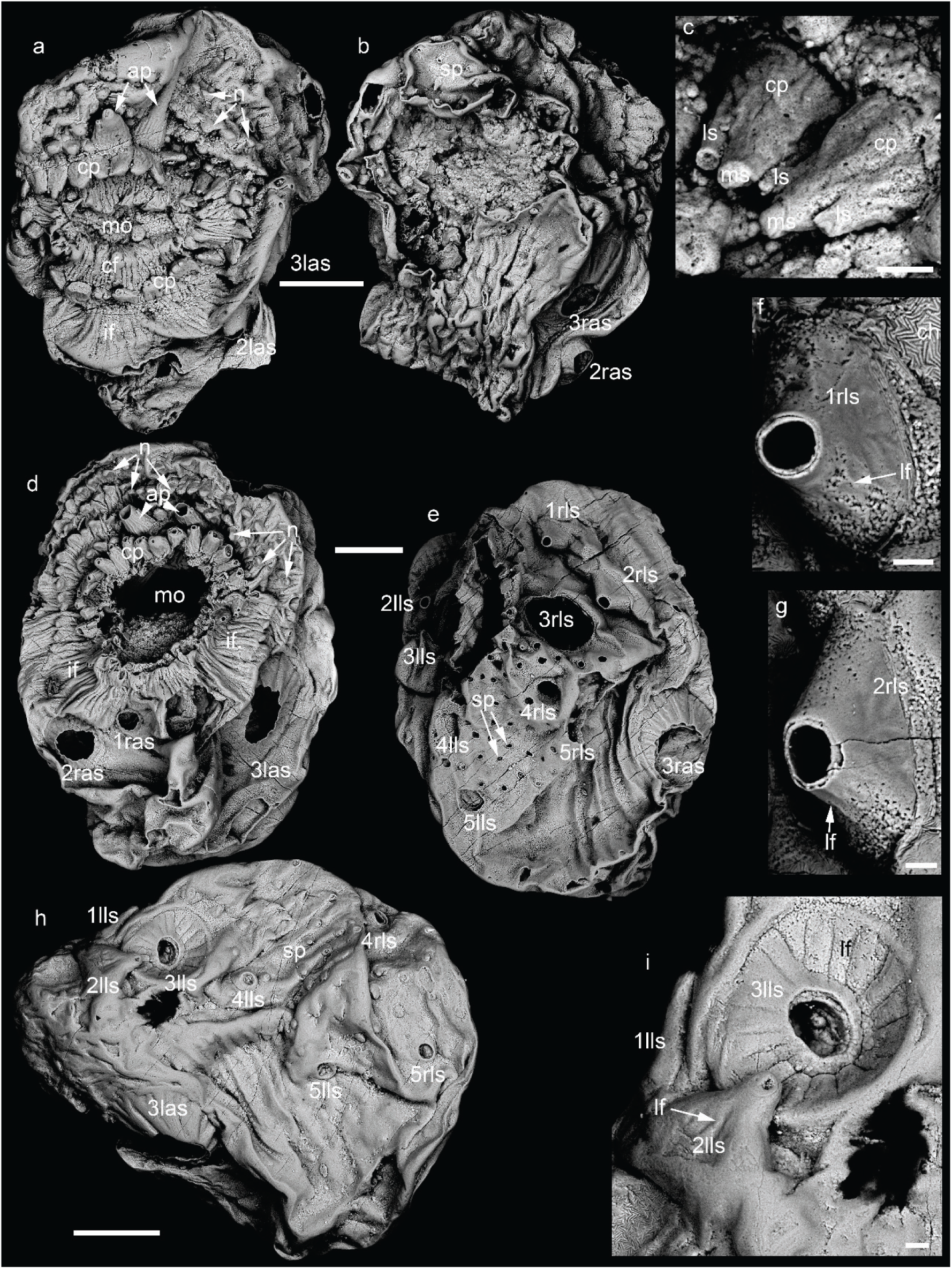
*Saccorhytus coronarius*. **a**–**c**, UMCU2016007, with two anterior protrusions; **a**, anterior view; **b**, posterior view; **c**, detail of circumoral protrusions in central right of **a**; **d**–**g**, UMCU2019019, with two anterior protrusions; **d**, anterior view; **e**, posterior view; **f**, **g**, detail of spinose sclerites in central upper and upper right of **e**; **h**, **i**, UMCU2018012, same specimen as in Fig. 2j; **h**, left view; **i**, detail of spinose sclerites in upper left of **h**. Scale bar: 200 μm (**a**, **b**, **d**, **e**, **h**), 20 μm (**c**, **f**, **g**, **i**). See Fig. 1 for abbreviations.

**Extended Data Figure 7.**
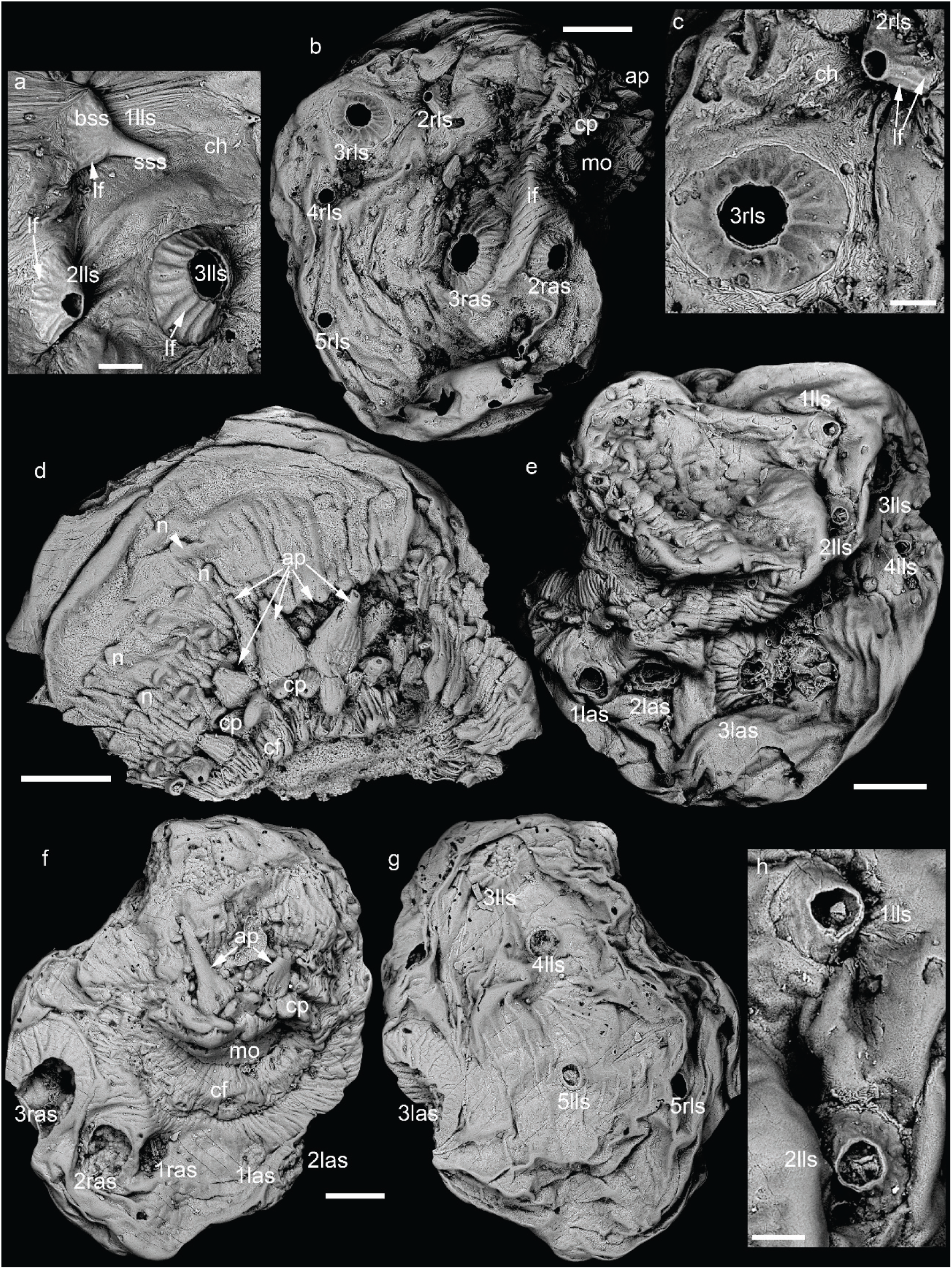
*Saccorhytus coronarius*. **a**–**c**, UMCU2016008, same specimen as in Fig. 2f, with three anterior protrusions; **a**, detail of spinose sclerites in central upper of Fig. 2f; **b**, right view; **c**, detail of spinose sclerites in upper left of **b**; **d**, UMCU2019020, a fragment with five anterior protrusions, dorsal anterior view; **e**, **h**, UMCU2020025; **e**, left view; **h**, detail of 1lls and 2lls, exhibiting round conical bases; **f**, **g**, UMCU2018015, with two anterior protrusions; **f**, anterior view; **g**, posterior view. Scale bar: 60 μm (**a**), 200 μm (**b**, **d**–**g**), 50 μm (**c**, **h**). See Fig. 1 for abbreviations.

**Extended Data Figure 8.**
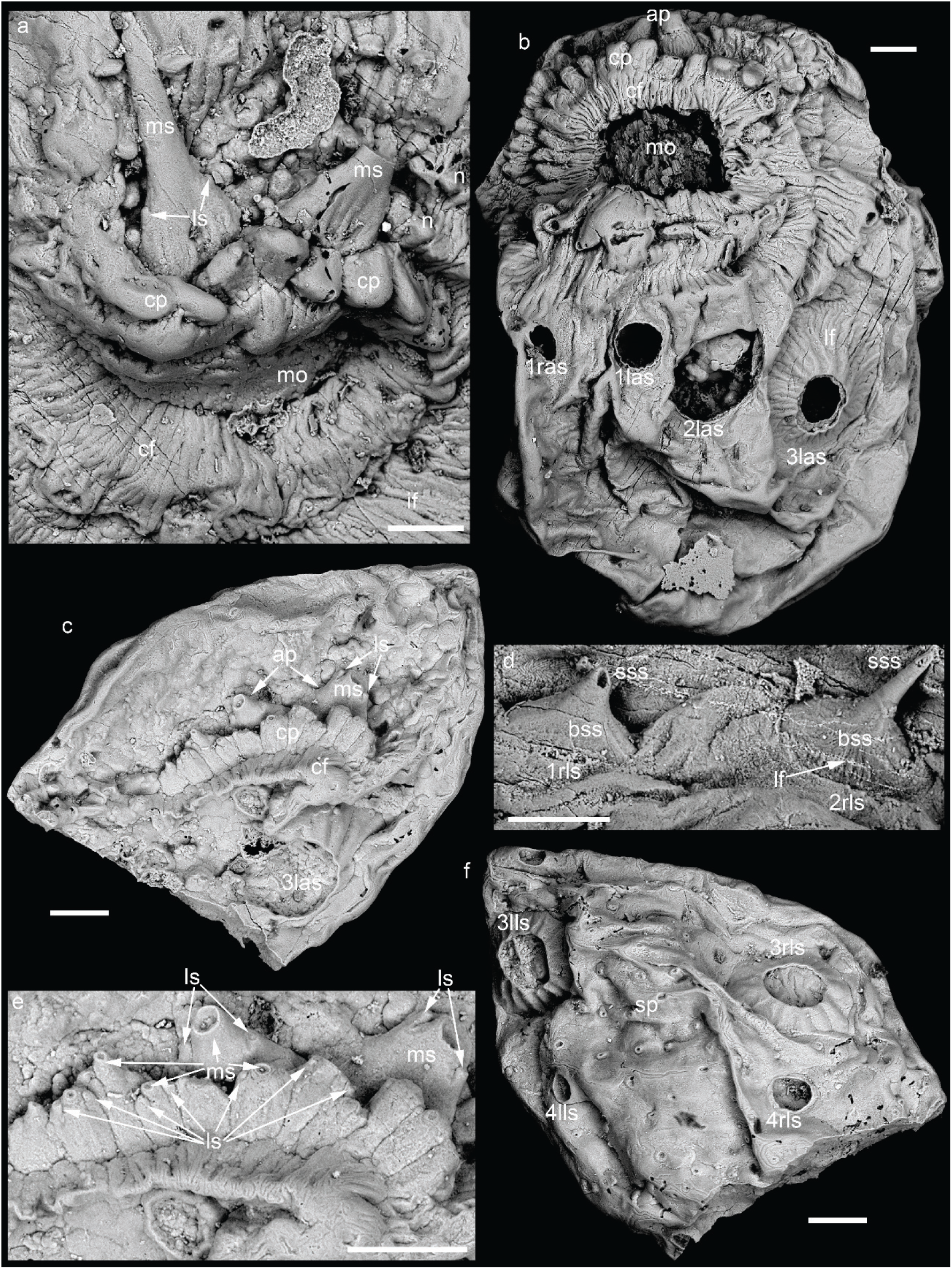
*Saccorhytus coronarius*. **a**, UMCU2018015, same specimen as in Extended Data Fig. 7f, exhibiting circumoral folds and anterior protrusions; **b**, **d**, UMCU2019018, same specimen as in Fig. 2l, with two anterior protrusions; **b**, ventral anterior view; **d**, detail of spinose sclerites in central upper of Fig. 2l; **c**, **e**, **f**, UMCU2014003, a fragment with two anterior protrusions; **c**, anterior view; **e**, detail of circumoral protrusions and anterior protrusions; **f**, posterior view. Scale bar represents 100 μm in all images. See Fig. 1 for abbreviations.

**Extended Data Figure 9.**
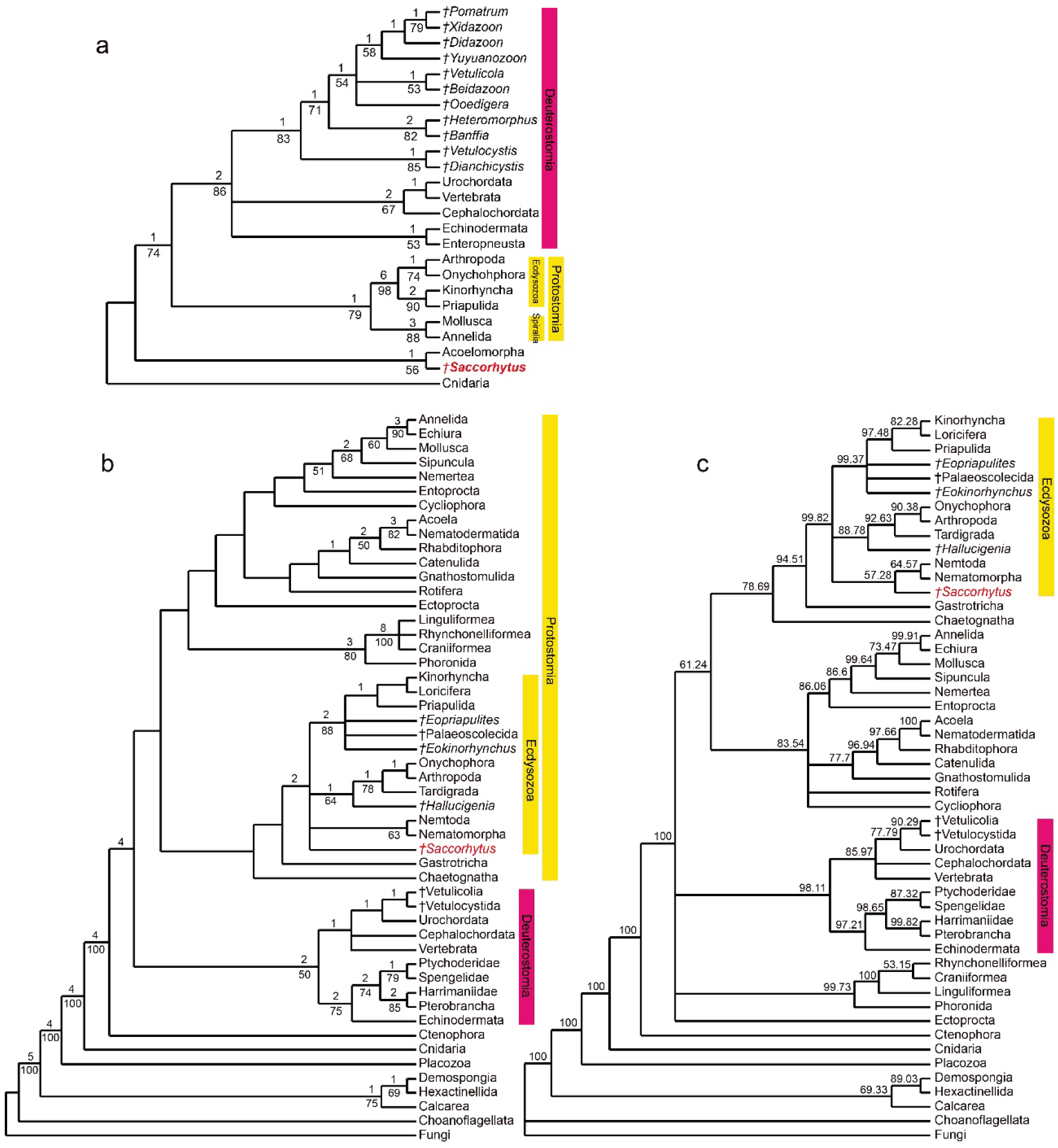
Results of phylogenetic analyses based on data matrix 1 (**a**) and data matrix 2 (**b, c**). **a**, strict consensus tree based on nine most parsimonious trees (tree length = 88, consistency index = 0.716, retention index = 0.866); **b**, 50% majority rule tree based on 22,848 most parsimonious trees (tree length = 285, consistency index = 0.568, retention index = 0.827); in **a** and **b**, numbers above nodes are Bremer support values (up to extra 10 steps), and numbers below nodes are Bootstrap support values (only values greater than 50% are shown); **c**, Bayesian 50% majority rule tree, numbers above nodes are clade credibility values. Symbol † denotes extinct taxon.

**Extended Data Figure 10.**
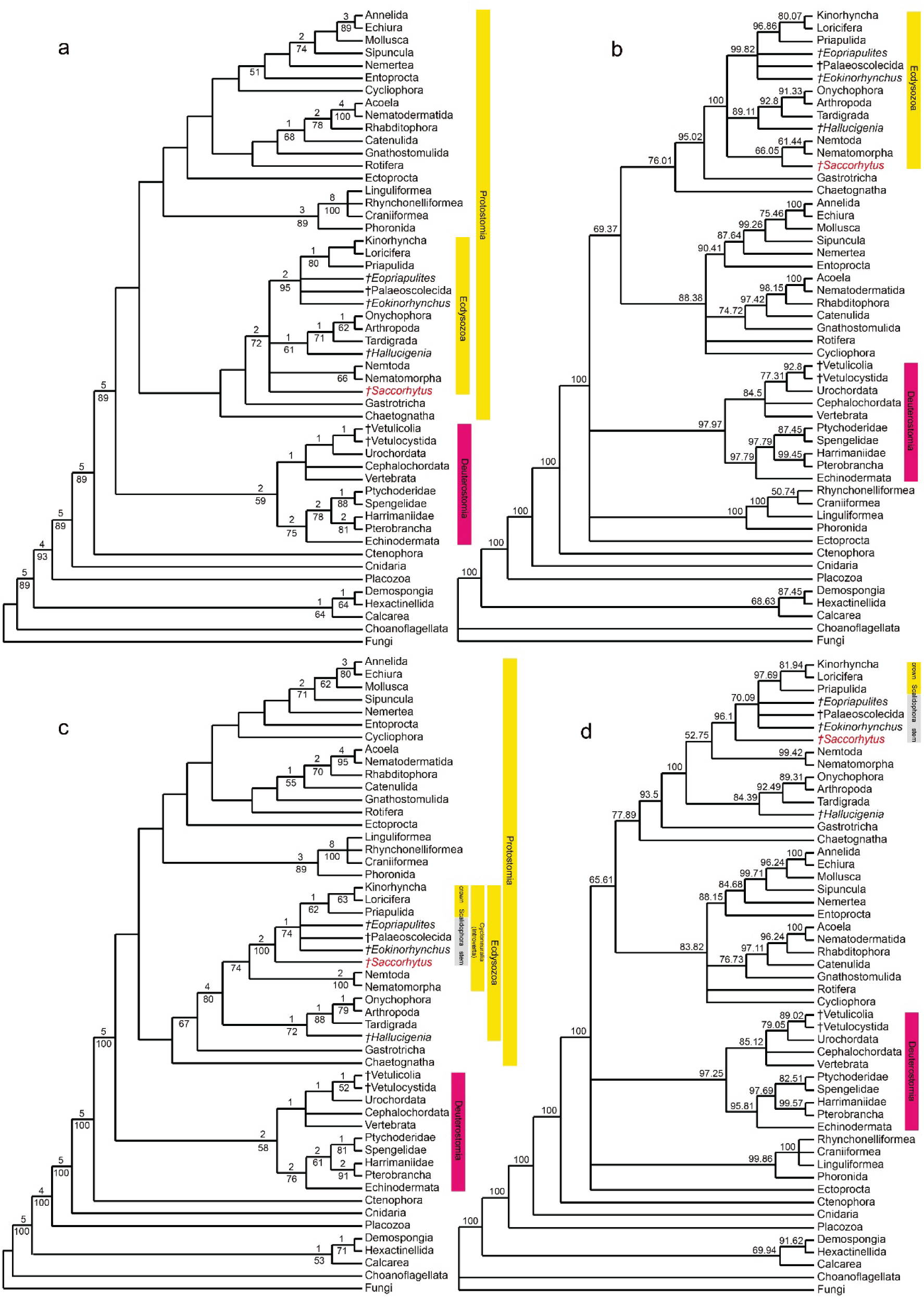
Results of phylogenetic analyses based on data matrix 3 (**a**, **b**) and data matrix 4 (**c**, **d**). **a**, 50% majority rule tree based on 22,848 most parsimonious trees (tree length = 285, consistency index = 0.568, retention index = 0.827); **b**, Bayesian 50% majority rule tree; **c**, 50% majority rule tree based on 3,264 most parsimonious trees (tree length = 285, consistency index = 0.568, retention index = 0.828); **d**, Bayesian 50% majority rule tree. In **a** and **c**, numbers above nodes are Bremer support values (up to extra 10 steps), and numbers below nodes are Bootstrap support values (only values greater than 50% are shown). In **b** and **d**, numbers above nodes are clade credibility values. Symbol † denotes extinct taxon.

## Supplementary Information

### Systematic palaeontology

Bilateria Hatschek^30^

Protostomia Grobben^31^

Ecdysozoa Aguinaldo et al.^1^ total group

Phylum and Class uncertain

Order Saccorhytida Han, Shu, Ou, and Conway Morris in Han et al.^8^ Family Saccorhytidae Han, Shu, Ou, and Conway Morris in Han et al.^8^, **emended.**

**Emended diagnosis:** Millimetric, ellipsoidal body with an anterior mouth but no anus, a spinose armor including circumoral protrusions, anterior protrusions, spinose sclerites, and posterior spines.

Genus *Saccorhytus* Han, Shu, Ou, and Conway Morris in Han et al.^8^, **emended.**

1988 *Clypecella* Li^32^ [nom. nud.]

2017 *Saccorhytus* Han, Shu, Ou, and Conway Morris in Han et al.^8^

**Type species:** *Saccorhytus coronarius* Han, Shu, Ou, and Conway Morris in Han et al.^8^

**Emended diagnosis:** As for the emended diagnosis of the type species.

**Remarks:** Similar fossils were first described as *Clypecella hexinsis* Li, 1988 (ref.^32^), but no holotype was designated. According to ICZN, *Clypecella* and *Clypecella hexinsis* are invalid.

*Saccorhytus coronarius* Han, Shu, Ou, and Conway Morris in Han et al.^8^, **emended.**

Figures 1, 2, 3a–c; Extended Data Figures 2–8; Supplementary Movies 1–10

1988 *Clypecella hexinsis* Li^32^, p. 63, 64, pl. 1, figs. 1, 2 [nom. nud.]

2017 *Saccorhytus coronarius* Han, Shu, Ou, and Conway Morris in Han et al.^8^, p. 229, fig. 1; p. 230, figs. 2, 3; extended data figs. 2–6; supplementary table 1.

**Holotype:** Specimen XX45-20, as was originally designated by Han et al.^8^, and is deposited at the Early Life Institute, Northwest University, Xi’an, China.

**Material and repository information:** A total of 270 specimens were extracted and examined, and 25 well-preserved specimens are illustrated in this paper. Specimens of *Saccorhytus coronarius* illustrated in this paper are deposited at the University Museum of Chang’an University (UMCU), and the accession numbers are UMCU2014001–2014005, 2016006–2016010, 2018011–2018015, 2019016–2019020, and 2020021–2020025.

**Occurrence:** Kuanchuanpu Formation, Zhangjiagou section, Dahe Village, Xixiang County, southern Shaanxi Province, South China; small shelly fossil *Anabarites trisulcatus*-*Protohertzina anabarica* Assemblage Zone, about 535 Ma, Fortunian Stage, Cambrian System.

**Emended diagnosis:** Millimetric, ellipsoidal animal with ventral half slightly wider than the dorsal half; integument two-layered and non-ciliated; anterior mouth surrounded by circumoral folds, one circlet of circumoral protrusions, one to five anterior protrusions, all protrusions bearing a central main spine flanked by two lateral spines with a closed tip; three pairs of spinose sclerites antero-ventrally, one to five pairs of spinose sclerites dorso-laterally, each with an expanded conical base that is ornamented with longitudinal folds and supports a long apical spine with a closed tip; nodes antero-dorsally; posterior spines with a sharp tip; anus absent.

**Preservation:** All the specimens are three-dimensionally phosphatized although most are folded or deformed. The integument is flexible and deformed in most specimens, implying that they were originally non-biomineralized. SRXTM analyses revealed no trace of internal anatomies (Extended Data Figs. 2c, 3c, f), and only the two-layered integument is preserved. In some cases, the two layers are closely adhered to each other but are discernable in SEM and SRXTM (Extended Data Figs. 2d, 3c); it is possible that these two layers represent the sub-layers of cuticle^8^. In other cases, the two layers are detached from each other, with a significant gap in between (Fig. 1d), indicating that the two layers may be phosphatic coating on two detached organic sheets (possibly representing taphonomically detached sub-layers of cuticles) that served as substrates for phosphatic coating.

No cilium insertion sites were observed on the integument, even in high-magnification SEM images. Cilia are rarely preserved in the fossil record, but putative cilia have been reported from Cambrian pterobranch fossils^33^ and compound cilia from Cambrian ctenophore fossils^34^. We infer that the integument of *Saccorhytus* is non-ciliated because no cilium insertion sites are found on the integument, but acknowledge this inference is based on the absence of evidence. Thus, in ensuing cladistic analyses (see below) the cilium-related characters are variously coded as “absent”, “uncertain”, or “present” in order to test the sensitivity of our phylogenetic interpretation to these characters.

An important observation made in this study is the preservation of spines, including the apical spines on spinose sclerites as well as the posterior spines in the back of the body. As mentioned in the main text and discussed further below, most spinose sclerites and posterior spines are broken, leaving a hole on the conical base of the spinose sclerites or on the posterior part of the body integument. In rare cases, however, spines are partially or completely preserved. We note that complete preservation of spines happens only when the spines are adpressed or tucked on the body integument so that they escaped from being abraded during fossil preservation and preparation. Given that microfossils in the Zhangjiagou Lagerstätte constitute grains in phosphatic limestone (see extended data fig. 1 in Han et al.^8^), it is likely that they have been locally reworked. Thus, it is possible that most spines were broken during reworking, particularly if they were relatively rigid structure sticking out on the surface of the globose fossil. The relative rigidity is consistent with the observation that the spines typically are not folded or deformed (e.g., Extended Data Fig. 4k), and implies that the spines may have been more robustly sclerotized than the rest of the integument or even biomineralized.

**Orientation:** *Saccorhytus* has a roughly globose body with a prominent mouth, and as such the oral-aboral axis defines a major polarity of the animal. Because in most living bilaterian animals the mouth is located at or near the anterior end, the mouth is taken as the anterior end of *Saccorhytus*, and the opposite side as the posterior end. When the mouth is flattened, it divides the body into two halves, with one half slightly wider than the other (Extended Data Fig. 2a, b; Supplementary Movie 3). The wider half is regarded as the ventral side because it helps to balance the animal for hydrodynamic stability. The anterior protrusions only occur on the dorsal side. The orientation adopted here is different from that of Han et al.^8^, who placed the mouth at a centro-ventral position, which is rather unusual for a bilaterian animal. Thus, these two orientations are rotated 90° from each other about the left-right axis. We note that Han et al.’s orientation was based on the holotype^8^, which is a folded specimen (their fig. 1a). Because of its mechanical weakness, the mouth is expected to cave in when folding occurs, so that the anterior and posterior ends are brought closer to each other, with the mouth located in the central concavity in the folded specimen. In unfolded specimens (Fig. 1a; Extended Data Figs. 2a, d, 3d, 4a, 6d; Han et al.’s fig. 1g, Extended Data Fig. 6f), the mouth is terminally located and defines one polar end of the body. Because Han et al.’s orientation was based on the taphonomically folded holotype^8^ and is inconsistent with location of the mouth in most living bilaterian animals, the orientation of *Saccorhytus* is here revised.

The ventral side *Saccorhytus* is mostly free of spinose sclerites and spines. In addition, the ventral side is wider than the dorsal side, perhaps helping to balance the animal and to achieve hydrodynamic stability. Thus, we infer that the ventral side of *Saccorhytus* was in contact with the benthic substrate (if *Saccorhytus* was an epibenthic animal) or with sand grains (if it was an interstitial animal).

**Comparison with Han et al.’s description and reconstruction:** The general morphology of the current specimens accords well with those described in Han et al.^8^, including a bag-shaped body, a prominent mouth surrounded by radial folds, and at least four pairs of spinose sclerites with an expanded base ornamented with longitudinal folds.

The key difference between our reconstruction and Han et al.’s reconstruction^8^ has to do with the structures described as spinose sclerites in this paper but as body cones in Han et al.^8^. Our observation indicates that the sclerites 1–3l/ras correspond with the body cones L/Rbc1–3 of Han et al.^8^, and the sclerites 3l/rls with the body cones L/Rbc4 of Han et al.^8^, whereas the sclerites 1–2l/rls and 4–5l/rls are structures newly identified in this study. Of these spinose sclerites, 3l/ras and 3l/rls are the largest and ornamented with more prominent longitudinal folds on the conical base, whereas the other spinose sclerites are smaller and sometimes the longitudinal folds are weakly developed. But these spinose sclerites are at least two to four times larger than the posterior spines (Supplementary Table 1), which do not have a well-defined conical base and lack longitudinal folds. These observations, together with the fact that the spinose sclerites are consistently arranged in a dorso-lateral set and an antero-ventral set, indicate that they are related or serially homologous structures. In our reconstruction, these spinose sclerites are composed of an expanded conical base ornamented with longitudinal folds and supporting a distally closed apical spine. In Han et al.’s reconstruction^8^, the body cones were understood as conical structures with a biologically programmed aperture and were interpreted as gill slits.

We contend that Han et al.’s reconstruction^8^ was misguided by broken sclerites and that the opening on the conical base is not a biologically programmed structure, but rather a taphonomic artifact related to physical damage. In both our specimens and Han et al.’s specimens^8^, the spinose sclerites are mostly damaged; breakage typically occurs at the junction between the conical base and the apical spine (Fig. 2f, j, l, Extended Data Fig. 6f, g; Han et al.’s fig. 2d, Extended Data Fig. 5e), but can also occur at the base of the sclerite (Fig. 1a; Extended Data Figs. 2d, 4b, 8b; Han et al.’s figs. 1d, g, extended data figs. 2k and 4h), the middle of the conical base (Fig. 1c; Extended Data Figs. 2d, 5e; Han et al.’s fig. 2g, extended data figs. 3g), the lower part of the apical spine (Figs. 1h, i, 2c, e), and the distal end of the apical spine (Fig. 2h, k; Extended Data Figs. 3i, 4k, 8d). Thus, the height of the surviving conical base varies greatly (compare Figs. 1c, h, 2c, d, e, and Extended Data Figs. 2d, 8b in this paper; compare Han et al.’s extended data figs. 2k, 3c, g, 4e). Evidence for physical damage also comes from the following observations. First, the rim of most conical bases are jagged (Fig. 1c, Extended Figs. 2d, e, 4a, b, 5d, e, 8b), exposing the broken edges of the two integument layers. This implies that these conical bases (or the “body cones”) were physically damaged; if the body cones had a biologically programmed and taphonomically intact aperture, the two integument layers would be continuous and would wrap around the rim of the aperture. Second, the relative size of the hole on broken bases varies greatly, ranging from 15% to 100% of the maximum diameter of the conical base (Supplementary Table 1). This is consistent with physical damage but inconsistent with the gill slit interpretation.

We argue that the spinose sclerites, including those equivalent to the body cones of Han et al.^8^, had an apical spine with a closed and pointed distal tip. A number of spinose sclerites bear nearly completely preserved apical spines (e.g., 1lls in Fig. 2h and Extended Data Fig. 7a; 2rls in Fig. 2k; 1rls and 2rls in Extended Data Fig. 3i; 1lls in Extended Data Fig. 4k; 1rls and 2rls in Extended Data Fig. 8d), which are always adpressed against the body wall and thus escaped from being abraded during fossil preservation and preparation. More importantly, spinose sclerites that are directly equivalent to “body cones” of Han et al.^8^ also bear remnants of the apical spine (3ras in Fig. 1h and Fig. 2c; 3las in Fig. 2e). This is strong evidence that the “body cones” are broken spinose sclerites, and they cannot be interpreted as gill slits.

There are other minor differences between our and Han et al.’s reconstructions. Although Han et al.^8^ did not explicitly identify the spinose sclerites 1–2 and 4–5l/rls, their fig. 2h did illustrate a spinose sclerite (1rls in our terminology) with an expanded conical base, and their fig. 2h also exhibited (but not labeled) the remnant of an expanded conical base of a spinose sclerite (2rls in our terminology). However, their reconstruction (fig. 3a of Han et al.^8^) depicted such sclerites as conical spines without an expanded base, thus glossing over the similarity between the closed spinose sclerites and open “body cones”. As another example, Han et al.^8^ reconstructed the circumoral protrusions as tubercles (their fig. 3, even though their fig. 1c illustrated a spinose protrusion), and our observation based on more completely preserved specimens shows that both the circumoral and anterior protrusions are tridented structures each with three spines. Here again, the tridented spines are preserved only when they are adpressed against the body wall, and in most cases they are either partially or entirely broken. Like the broken spinose sclerites, the broken circumoral and anterior protrusions leave a series of holes around the mouth (see Han et al.’s fig. 1g). It is interesting to note that these holes are treated differently in Han et al.’s^8^ reconstruction: the circumoral protrusions were depicted as closed tubercles, the body cones were reconstructed as open gill slits, and the anterior protrusions were not included in their reconstruction. In our reconstruction, these holes are all regarded as taphonomic artifacts related to physical breakage. Finally, Han et al.^8^ illustrated numerous “circular pores” on the body wall. In our opinion, these circular pores are also taphonomic artifacts: some represent broken posterior spines (e.g., their fig. 1f), others may represent broken dorso-lateral spinose sclerites (the largest hole in their extended data fig. 2a), and still others may simply be diagenetic artifacts (e.g., circular pore within Rbc3 in their fig. 1a). Better preserved specimens in our collection clearly demonstrate that posterior spines are present on much of the posterior and part of the dorsal and lateral sides of the animal. Once again, the posterior spines are not damaged by abrasion only if they are adpressed on the body wall.

**Taxonomic consideration**: Specimens of *Saccorhytus coronarius* in our collection do show some variations in the number of anterior protrusions and dorso-lateral spinose sclerites. For example, the number of the anterior protrusions varies from one to five, and the spinose sclerites 1–2l/rls and 4–5l/rls may not be present in all specimens. Considering the overall similarity of the specimens in our collection, we tentatively regard these minor variations in the number of anterior protrusions and dorso-lateral spinose sclerites as intraspecific variations (e.g., ontogenetic, ecophenotypic, or sexual variations). We note that intraspecific variations in the numbers of sclerites are common among the Cambrian ecdysozoans. For example, the number of sclerites (plates, microplates, and platelets) can vary greatly between individuals of the same palaeoscolecid species^35^. However, we do acknowledge that future investigation may recognize multiple species of *Saccorhytus* given that we did not observe any consistent relationship between body size and the number of anterior protrusions.

**Comments on possible similarity to scalidophorans**: Scalidophorans (i.e., priapulids, kinorhynchs, and loriciferans) are united by the presence of internally hollow scalids, which contain sensory cells and other living tissues^14, 23^. Similarly, scalidophoran fossils from the Fortunian Stage^5–7^ also have internally hollow scalids, and they are inferred to have contained sensory cells and other living tissues when alive. Other cycloneuralians such as nematoids may have superficially similar structures known as hooks, but these are simple thickenings of the cuticle and are internally solid^14, 23^. We note that the circumoral protrusions and anterior protrusions of *Saccorhytus* are also internally hollow, and by analogy they may have contained sensory cells and other living tissues as well. More importantly, the circumoral protrusions are radially arranged around the mouth opening, although the scalidophoran scalids exhibit pentaradial arrangement whereas *Saccorhytus*’ circumoral protrusions exhibit biradial arrangement. Thus, although the arrangement and symmetrical patterns are different between the scalids of scalidophorans and the circumoral/anterior protrusions of *Saccorhytus*, it is possible that these structures are homologous. The possibility is further support by the remarkable similar spinose sclerites between *Saccorhytus* and the co-occurring stem-group scalidophorans *Qinscolex* and *Shanscolex*^7^ and the coeval stem-group kinorhynch *Eokinorhynchus*^6^. These spinose sclerites are internally hollow structures with an expanded conical base and an elongate apical spine. Thus, a phylogenetic relationship with the scalidophorans is an intriguing possibility for *Saccorhytus*, and this should be further tested with additional evidence.

#### Taxa, characters, and codings for phylogenetic analyses

**Data matrix 1** was modified from ref^8^, and *Saccorhytus* was coded as a total-group bilaterian with a terminal mouth and without an anus. The selected taxa and characters follow those of ref^8^, whereas the codings of *Saccorhytus* were revised according to the new observations made in this study. Specifically, codings of characters 9, 11, 22, 24, 26, 27, 31, 32, 34, 37–44, 46–48, 51, 54, and 55 are revised (Supplementary Table 2). An analysis of the revised data matrix of ref^8^ shows that *Saccorhytus* is not a deuterostome, but instead forms a sister-group relationship with the Acoelomorpha (Extended Data Fig. 9a). The radially arranged circumoral structures do not unite *Saccorhytus* with vetulicolians, because *Saccorhytus* lacks key deuterostome apomorphies such as pharyngeal gill slits. Instead, *Saccorhytus* shares with the acoelomorphs characters such as radially arranged circumoral structures and absence of anus. Here, radially arranged circumoral structures are homoplastic structures between *Saccorhytus* and vetulicolians.

**Data matrices 2–4** were modified and expanded from ref^19^. In order to test the relationships between *Saccorhytus* and fossil ecdysozoans and deuterostomes, we included the Palaeoscolecida *sensu stricto*^35^, *Eopriapulites*^5, 36^, and *Eokinorhynchus*^6^ as representatives of Cambrian cycloneuralians, *Hallucigenia*^20^ as a representative of Cambrian stem-group arthropods, and Vetulicolia^16^ and Vetulocystida^17^ as representatives of Cambrian deuterostomes. We also split the Chordata into the Urochordata, Cephalochordata, and Vertebrata, so there are 51 selected taxa in the data matrix, with the Fungi as an outgroup. We also added 15 new characters to those in ref^19^, which are listed as below.

139. Endostyle^2, 14^: absent (0), present (1). This is an autapomorphy of the Deuterostomia. Endostyle is present in vetulicolians, but is uncertain in vetulocystids.

140. Endoskeleton mesodermally derived^8^: absent (0), present (1). This is an autapomorphy of the Deuterostomia, but the tunicates lack internal skeletons.

141. Cuticle^2^: absent (0), present (1). This character refers to the presence/absence of cuticle in the integument, irrespective of chemical composition of the cuticle (i.e., chitinous vs non-chitinous cuticle). Thus, this character is different from character 78 (cuticle with chitin). The presence of cuticle (and by extension, ecdysis) can be inferred for certain fossils, but it is difficult to determine the chemical composition of cuticles in the fossil record. Thus, character 78 is coded as “uncertain” for all fossil taxa.

142. Radially arranged circumoral structures^20^: absent (0), present (1). This is an autapomorphy of the total-group Ecdysozoa^20^, but it evolved convergently in acoelomorphs, and vetulicolians^16^ and vetulocystids^17^. The circumoral folds of *Saccorhytus* are regarded as circumoral structures and are coded as “present”.

143. Sclerotized pharyngeal teeth^20^: absent (0), present (1).

144. Trunk with macroannuli^6, 37^: absent (0), present (1).

145. Trunk with segmentation^2, 37^: absent (0), present (1).

146. Trunk with annular rings of sclerites^6^: absent (0), present (1).

147. Spinoscalids^2, 14^: absent (0), present (1). Character 112 refers to the presence of both spinoscalids and clavoscalids (coded 1, e.g., for loriciferans) or the lack of either spinoscalids or clavoscalids (coded 0, e.g., for all scalidophorans other than loriciferans). However, spinoscalids occur in crown-group scalidophorans as well as in all the Cambrian cycloneuralians (i.e., *Eopriapulites*, *Eokinorhynchus*, and the Palaeoscolecida), whereas clavoscalids occur only in crown-group loriciferans^2^. Thus, character 147 (spinoscalids) is added to refer specifically to the presence/absence of spinoscalids (coded 1 for all scalidophorans). The circumoral and anterior protrusions of *Saccorhytus* may represent scalids/spinoscalids, in which case character 111 (scalids) and character 147 (spinoscalids) are both coded as presence for *Saccorhytus*.

148. Filter feeding accomplished within pharynx^8^: absent (0), present (1).

149. Pharyngeal food transport groove^8^: absent (0), present (1).

150. Peripharyngeal atrium^8^: absent (0), present (1).

151. V- or W-shaped myomeres^8^: absent (0), present (1).

152. Neural crest cells^8^: absent (0), present (1).

153. Neurogenic placodes^8^: absent (0), present (1).

The original data matrix was revised according to Peterson and Eernisse’s comments in ref^19^. Specifically, the original character 117 was deleted and not included in Supplementary Table 3 or in phylogenetic analyses, so there are 152 characters in total. The codings of character 4 for Onychophora, character 25 for Acoela, Nematodermatida, and Catenulida, character 44 for Loricifera, character 78 for Entoprocta, character 87 for Onychophora and Arthropoda, original character 121 for Acoela, Nematodermatida, Catenulida, Onychophora, Sipunculan, Echiuran, and Ectoprocta, original character 125 for Placozoa, original character 127 for Acoela and Nematodermatida, and original characters 127–135 for sponges, were revised. Additionally, character 78 for Nematoda and Nematomorpha is revised to be “absent”, because the adult nematoids lack chitin in their cuticle^14^ (Supplementary Table 3).

For *Saccorhytus*, we employed several coding strategies to accommodate the uncertainty about several characters and to test the sensitivity of our phylogenetic results to such uncertainties. Key examples of these characters are listed as follows. **Characters 13 (ciliated epidermis), 14 (densely multiciliated epidermis), and 15 (distinct “step” in cilia)**: our preferred interpretation is that the integument of *Saccorhytus* is non-ciliated and the three characters were coded as (0, 0, 0), but we also tested alternative codings (?, ?, ?) and (1, ?, ?). **Character 87 (terminal mouth)**: we strongly prefer a terminal mouth for *Saccorhytus* (see discussion above on orientation), but we also tested an alternative coding of “uncertain” for this character. **Character 83 (ecdysis) and character 141 (cuticle)**: our preferred coding is (1, 1) but we also tested the alternative coding (?, ?). **Character 111 (scalids) and character 147 (spinoscalids)**: we tested two alternative codings (?, 0) and (1, 1).

Three data matrices (**Data matrices 2–4**) with alternative codings for characters 83, 111, 141, and 147 of *Saccorhytus* were analyzed using PAUP4^27^ and MrBayes3.2.7a^28^ software. In the analysis of each data matrix, we also carried out two **sensitivity tests** using alternative codings for characters 13–15 and 87 to evaluate the influence of presence/absence of cilia and a terminal mouth on the position of *Saccorhytus*. For example, although **character 13–15** were coded as (0, 0, 0) in data matrices 2–4 and corresponding cladograms in Extended Data Figs. 9B–C and 10, we also generated sensitivity-test cladograms using alternative codings of (?, ?, ?) and (1, ?, ?) for these three characters in each data matrix, and the results are shown below. Details of the data matrices and results of cladistic analyses are described below.

First, we coded *Saccorhytus* as a total-group bilaterian animal, using our reconstruction of *Saccorhytus* but not assuming the presence of cuticle, ecdysis, or scalids [i.e., these three characters were coded (?, ?, ?); **Data matrix 2**]. The results show that *Saccorhytus* is a total-group ecdysozoan (Extended Data Fig. 9b; PAUP4) or a stem-group nematoid (Extended Data Fig. 9c; MrBayes3.2.7a). The key features uniting *Saccorhytus* with ecdysozoans are characters 13 (cilia absent), 87 (a terminal mouth), and 142 (radially arranged circumoral structures, herein the circumoral folds). Although radially arranged circumoral structures also occur in acoelomorphs, vetulicolians, and vetulocystids, the results imply convergent evolution among these groups. The software MrBayes3.2.7a may have interpreted character 78 (cuticle with chitin) as absent, and as such *Saccorhytus* is united with the Nematoida (Extended Data Fig. 9c; MrBayes3.2.7a).

As a sensitivity test (**Sensitivity Test 1**), we recoded characters 13, 14, 15, and 87 as uncertain (“?”) and re-ran the analysis using PAUP4. We obtained 9,792 most parsimonious trees (tree length = 284, consistency index = 0.570, retention index = 0.828). The 50% majority rule tree indicates that *Saccorhytus* is resolved to be united with acoelomorphs (**Supplementary Fig. 1**). In this case, the software PAUP4 parsimoniously interpreted *Saccorhytus* as a ciliated animal without a terminal mouth, i.e., an acoelomorph-like animal.

As another sensitivity test (**Sensitivity Test 2**), characters 13–15 were coded as uncertain (“?”) but character 87 (terminal mouth) as present as we strongly prefer. A parsimonious analysis using PAUP4 recovered 32,640 most parsimonious trees (tree length = 285, consistency index = 0.568, retention index = 0.827). The 50% majority rule tree indicates that *Saccorhytus* falls within the total-group Ecdysozoa (**Supplementary Fig. 2**). In this case, the software parsimoniously interpreted *Saccorhytus* as an animal with a terminal mouth but with no cilia. This test shows that a terminal mouth is a key character that unites *Saccorhytus* with the Ecdysozoa.

**Supplementary Fig. 1.**
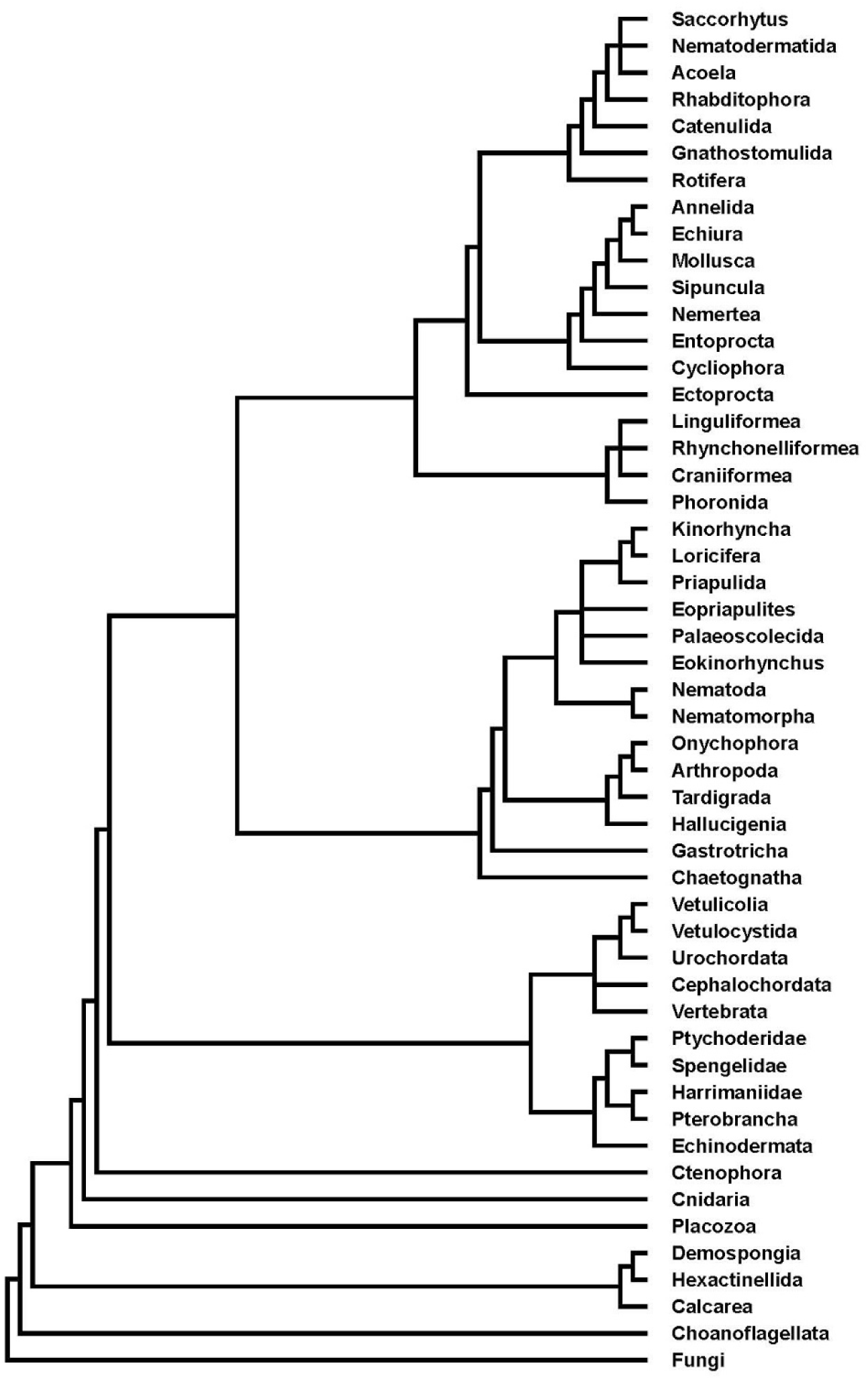
Phylogenetic result of Sensitivity Test 1.

**Supplementary Fig. 2.**
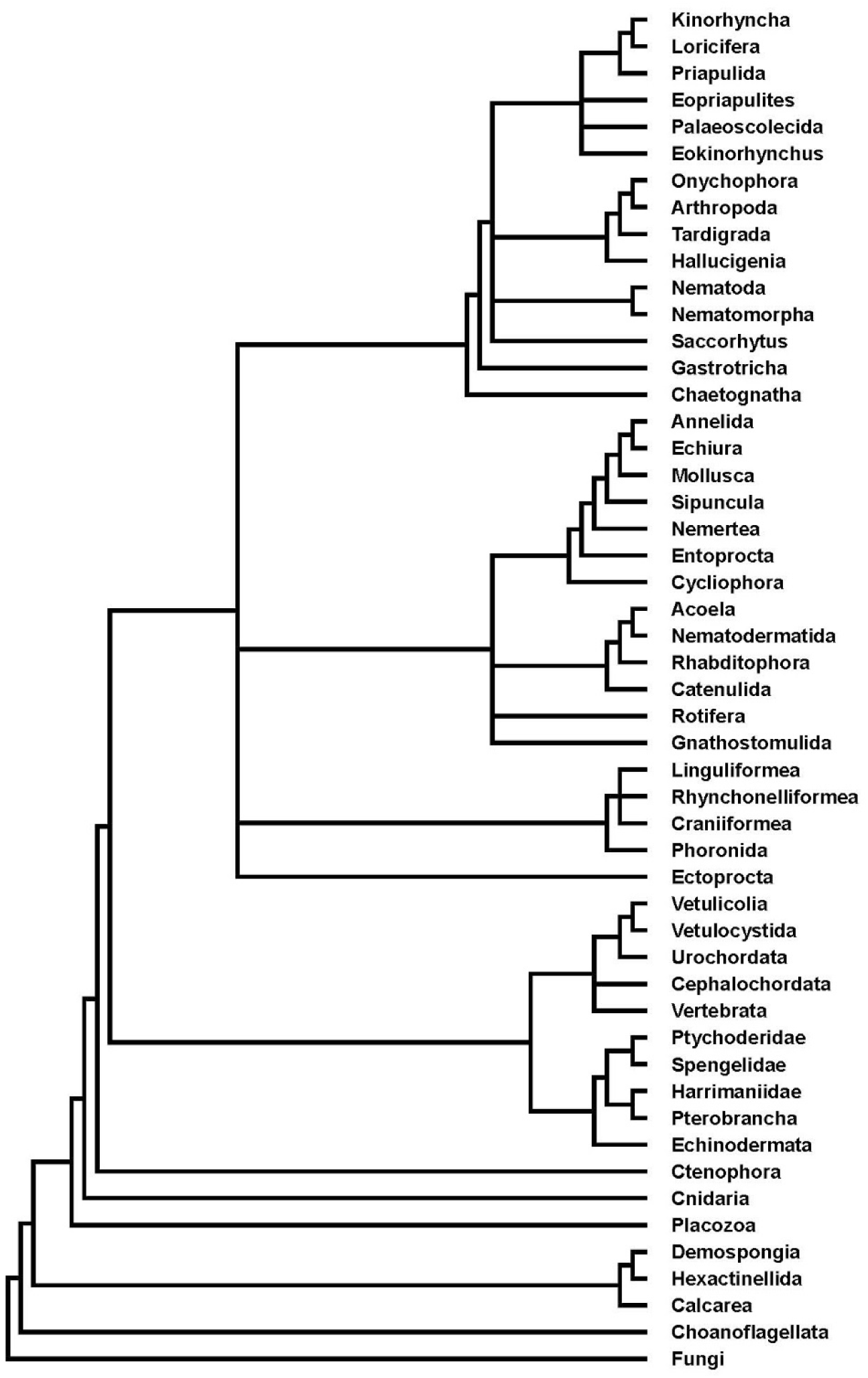
Phylogenetic result of Sensitivity Test 2.

In **Data matrix 3**, characters were coded as Data matrix 2 except for characters 83 (ecdysis) and 141 (cuticle). Because we interpret the two-layered integument of *Saccorhytus* as evidence for epicuticle and procuticle (with the innermost epidermis likely lost during degradation), the fossils represent molts. Accordingly, codings of characters 83 (ecdysis) and 141 (cuticle) were revised from (?, ?) to (1, 1). Similar to Data matrix 2, the results show that *Saccorhytus* falls within the total-group Ecdysozoa (Extended Data Fig. 10a; PAUP4) or stem-group Nematoida (Extended Data Fig. 10b; MrBayes3.2.7a). The key features uniting *Saccorhytus* with Ecdysozoa are the presence of cuticle, ecdysis, and radially arranged circumoral structures, whereas the stem-group Nematoida affinity may also result from the uncertainty about the presence of chitin.

Because the presence of cuticle does not necessarily exclude the presence of cilia, as in gastrotrichs^2^, as a sensitivity test (**Sensitivity Test 3**) we recoded characters 13–15 and 87 as uncertain (“?”) and re-ran the analysis using PAUP4. We recovered 22,848 most parsimonious trees (tree length = 285, consistency index = 0.568, retention index = 0.827). The 50% majority rule tree indicates that *Saccorhytus* falls within the total-group Ecdysozoa (**Supplementary Fig. 3**). Here the software PAUP4 parsimoniously interpreted *Saccorhytus* as an animal with a terminal mouth and with no cilia.

As another sensitivity test (**Sensitivity Test 4**), characters 13 was coded present, whereas characters 14, 15, and 87 were coded uncertain. A parsimonious analysis using PAUP4 recovered 22,848 most parsimonious trees (tree length = 286, consistency index = 0.566, retention index = 0.826). The 50% majority rule tree indicates that *Saccorhytus* still falls within the total-group Ecdysozoa (**Supplementary Fig. 4**). This phylogenetic result implies that Saccorhytus may have inherited cilia as a plesiomorphy if it is a stem-group ecdysozoan, or independently acquired cilia if it is a crown-group ecdysozoan.

**Supplementary Fig. 3.**
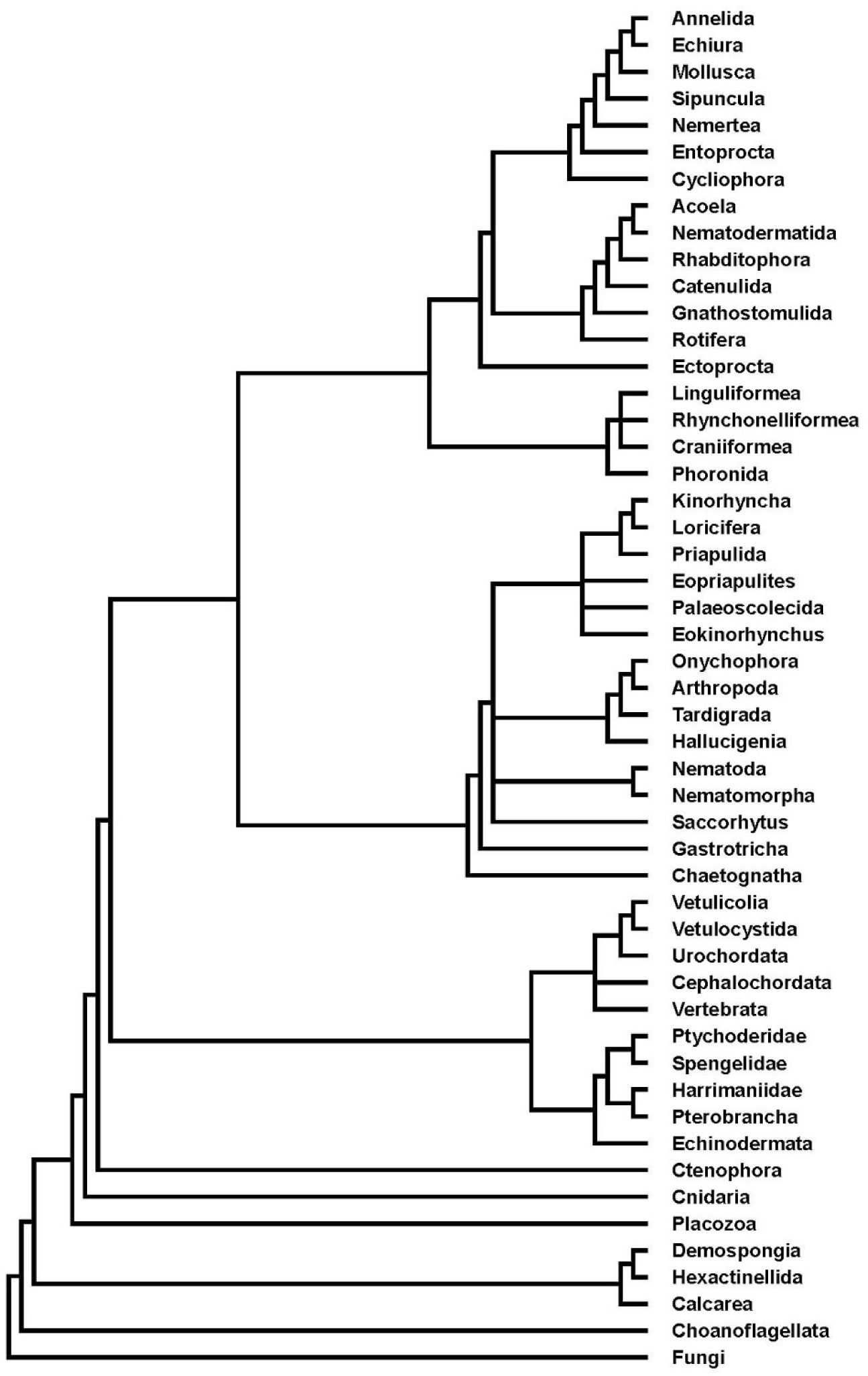
Phylogenetic result of Sensitivity Test 3.

**Supplementary Fig. 4.**
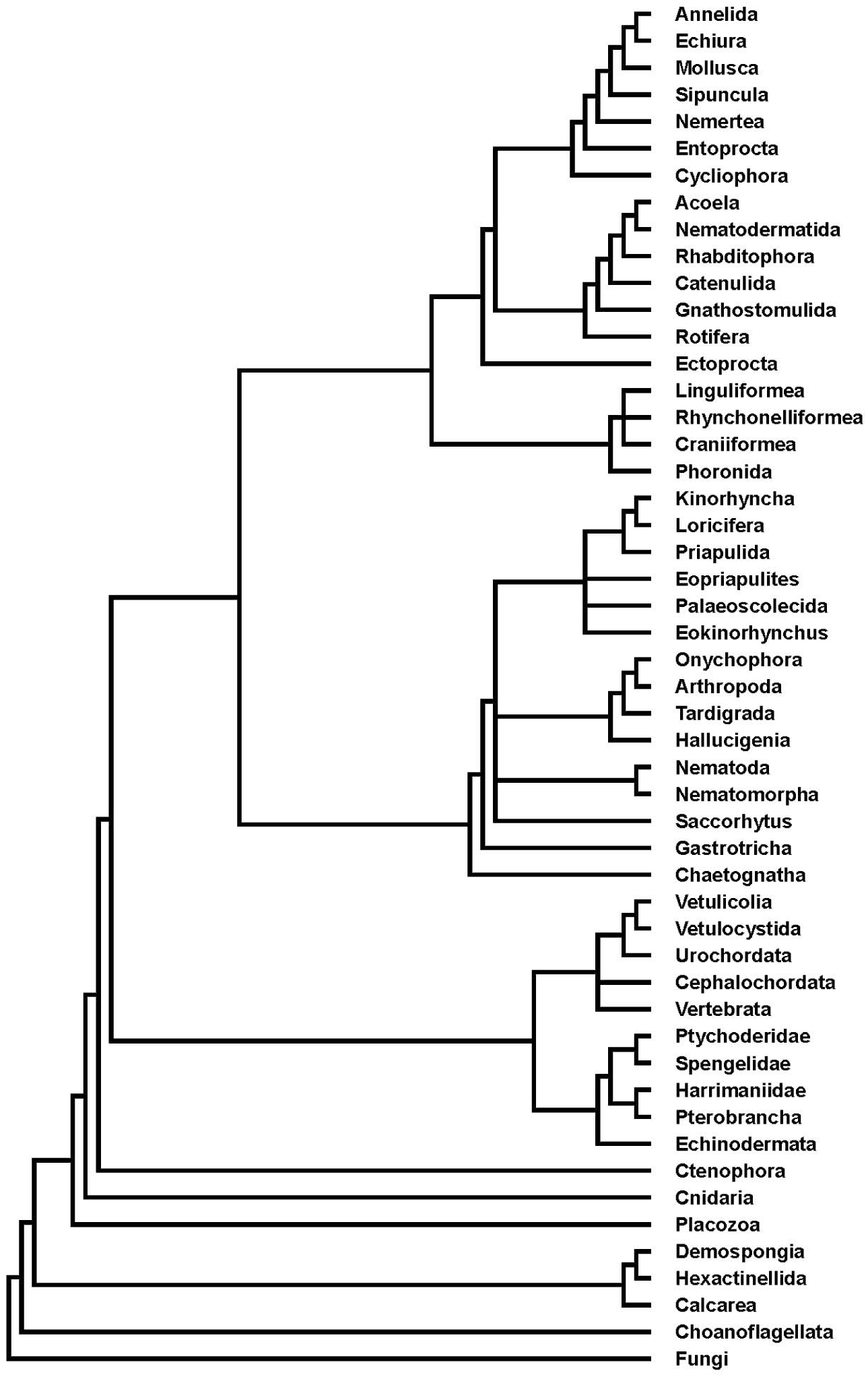
Phylogenetic result of Sensitivity Test 4.

In **Data matrix 4**, we retained the codings of Data matrix 3 except for characters 111 (scalids) and 147 (spinoscalids). Because of the possibility that the circumoral protrusions and anterior protrusions of *Saccorhytus* may be homologous with scalids (spinoscalids), characters 111 (scalids) and 147 (spinoscalids) were both coded as present in this data matrix. The results show that *Saccorhytus*, together with *Eopriapulites*, *Eokinorhynchus*, and the Palaeoscolecida *sensu stricto*, are stem-group scalidophorans (Extended Data Fig. 10c, d), and the key feature uniting these taxa is the presence of scalids/spinoscalids.

As a sensitivity test (**Sensitivity Test 5**), we recoded characters 13–15 and 87 as uncertain (“?”) and re-ran the analysis using PAUP4. We obtained 3,264 most parsimonious trees (tree length = 285, consistency index = 0.568, retention index = 0.827). The 50% majority rule tree indicates that *Saccorhytus* is a stem-group scalidophoran (**Supplementary Fig. 5**). In this case, the software PAUP4 parsimoniously interpreted *Saccorhytus* as an animal with a terminal mouth but no cilia, and the result is the same as Data matrix 4 (Extended Data Fig. 10c).

As another sensitivity test (**Sensitivity Test 6**), we recoded characters 13 as present, and characters 14, 15, and 87 as uncertain. A parsimonious analysis recovered 3,264 most parsimonious trees (tree length = 286, consistency index = 0.566, retention index = 0.826). The 50% majority rule tree indicates that *Saccorhytus* is still a stem-group scalidophoran. This phylogenetic result implies that cilia originated secondarily within the total-group Scalidophora.

To sum up, a terminal mouth and radially arranged circumoral structures are key characters supporting *Saccorhytus* as a total-group ecdysozoan. With the inferred presence of cuticle and molting, *Saccorhytus* is firmly placed within the total-group Ecdysozoa. And with the inferred homology between circumoral/anterior protrusions and scalids/spinoscalids, *Saccorhytus* is resolved within the total-group Scalidophora. In contrast, the absence or presence of cilia on the integument has only minor influence on the phylogenetic interpretation of *Saccorhytus*.

**Supplementary Fig. 5.**
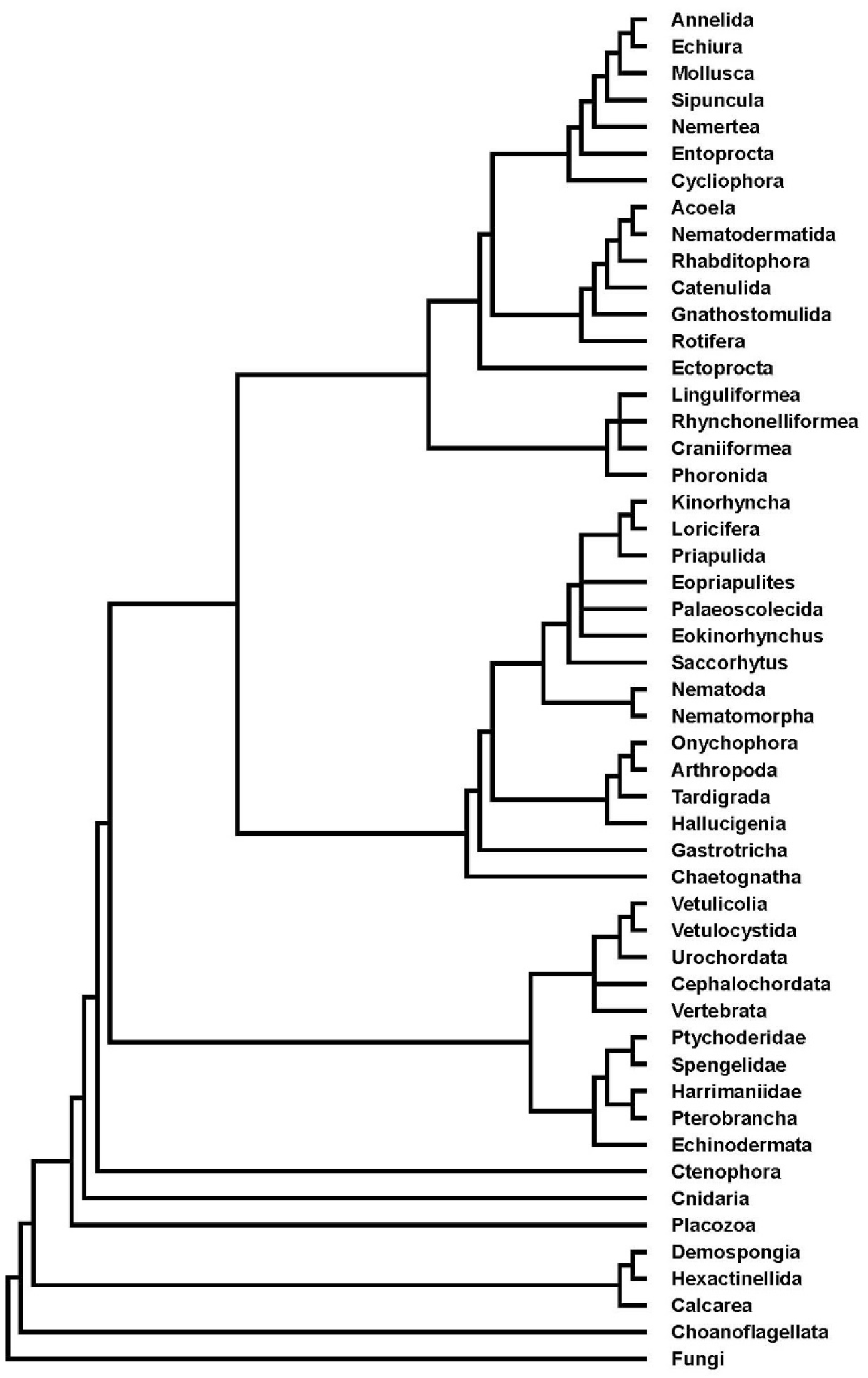
Phylogenetic result of Sensitivity Test 5.

**Supplementary Fig. 6.**
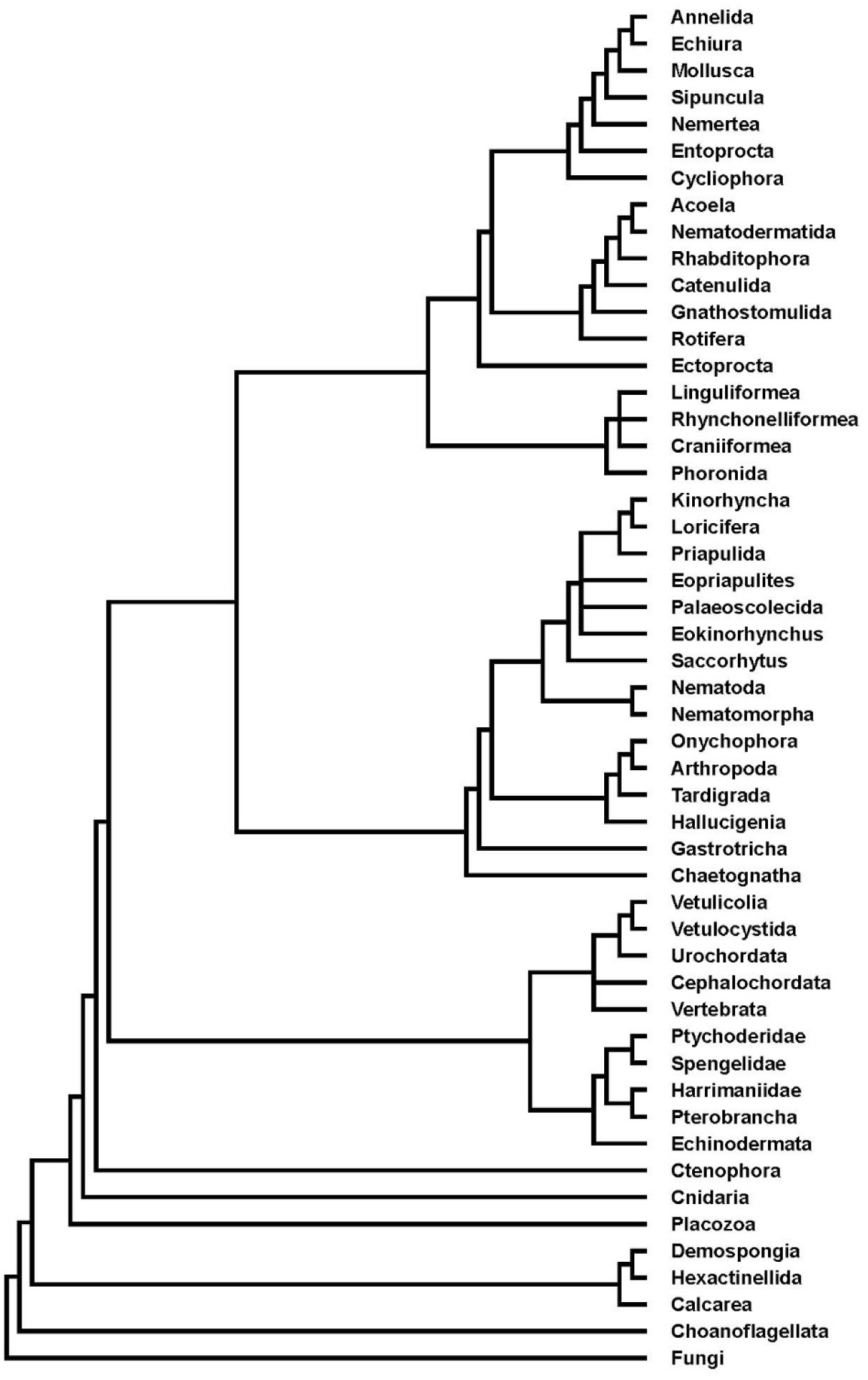
Phylogenetic result of Sensitivity Test 6.

#### Data matrices 1–4

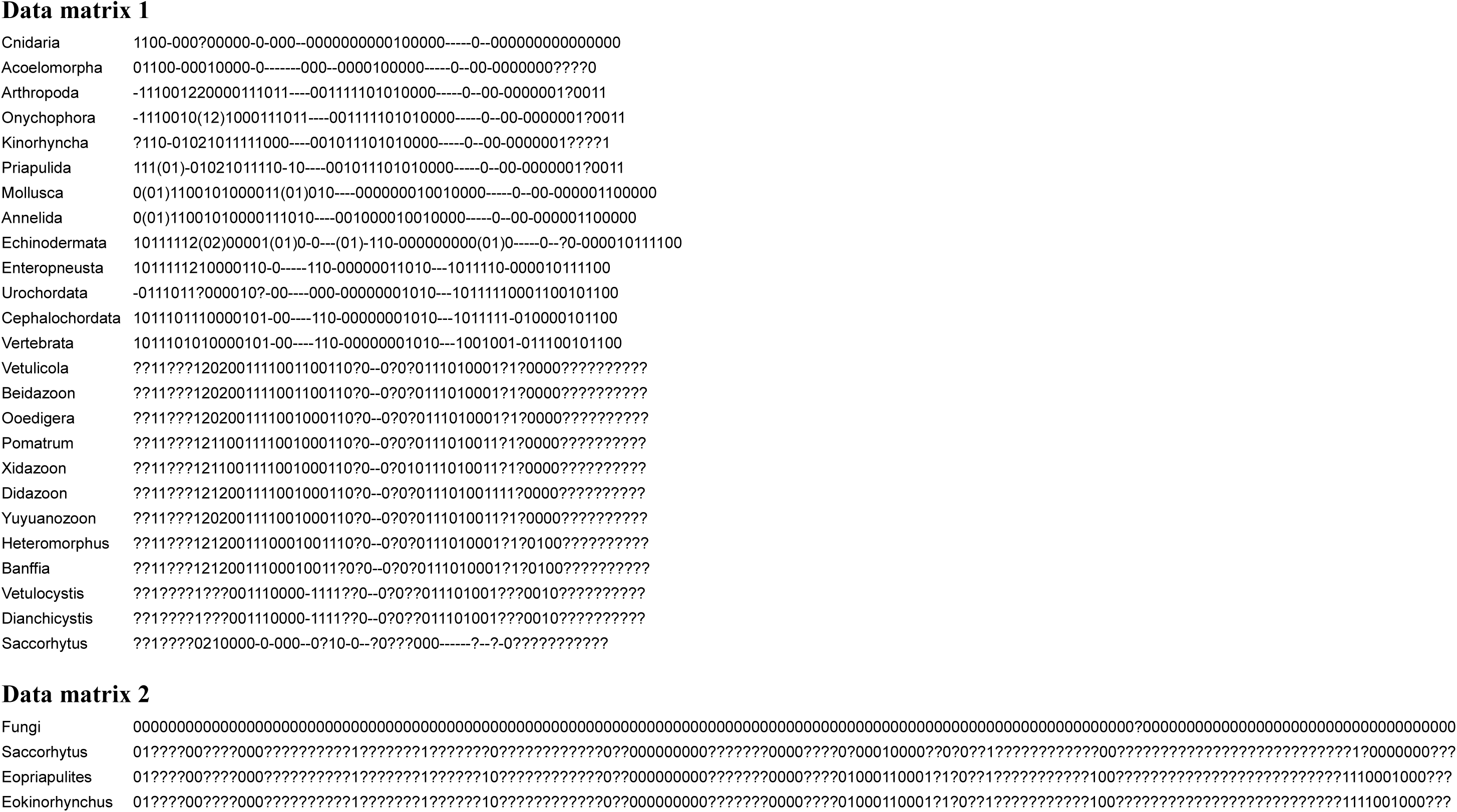
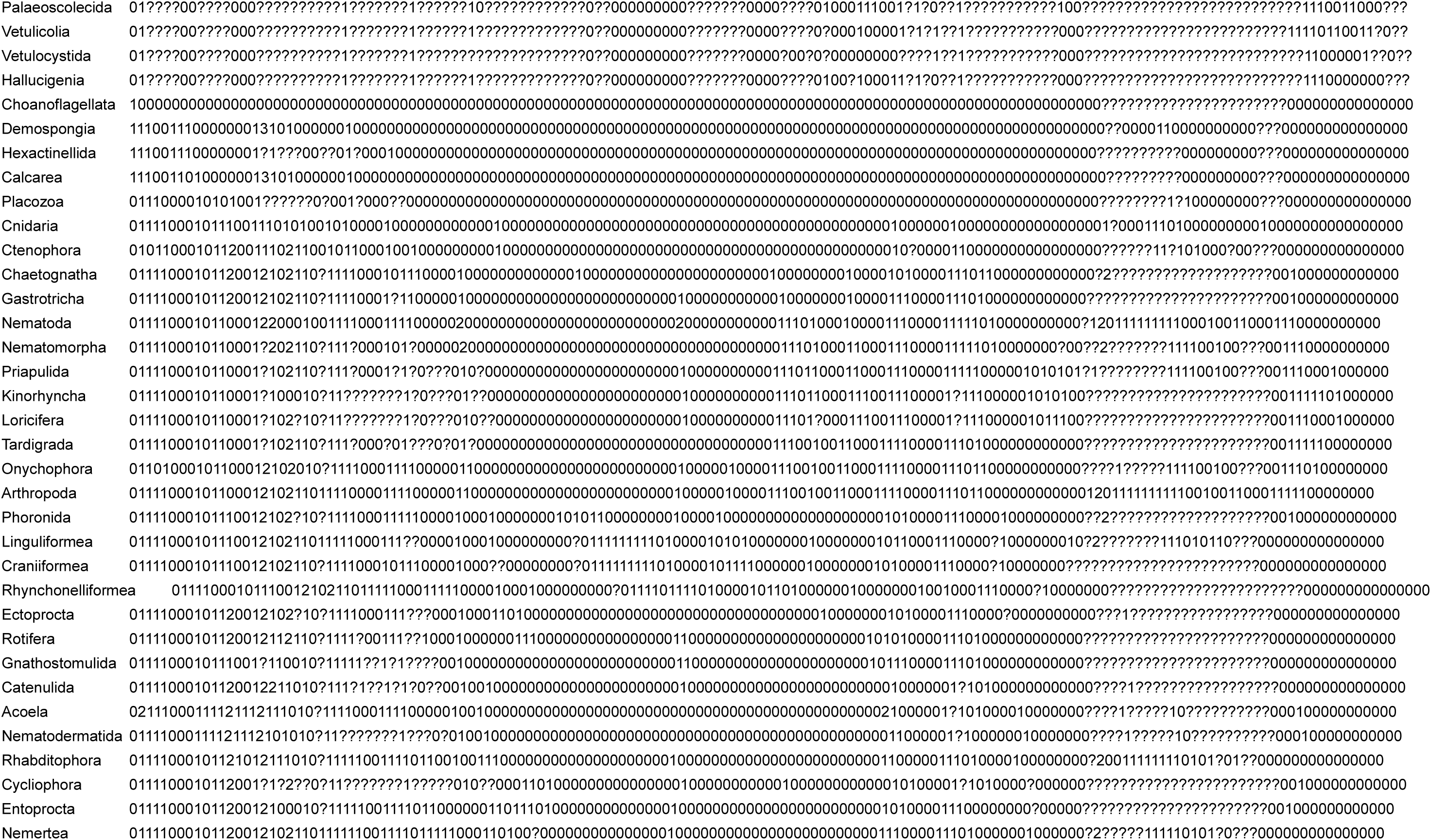
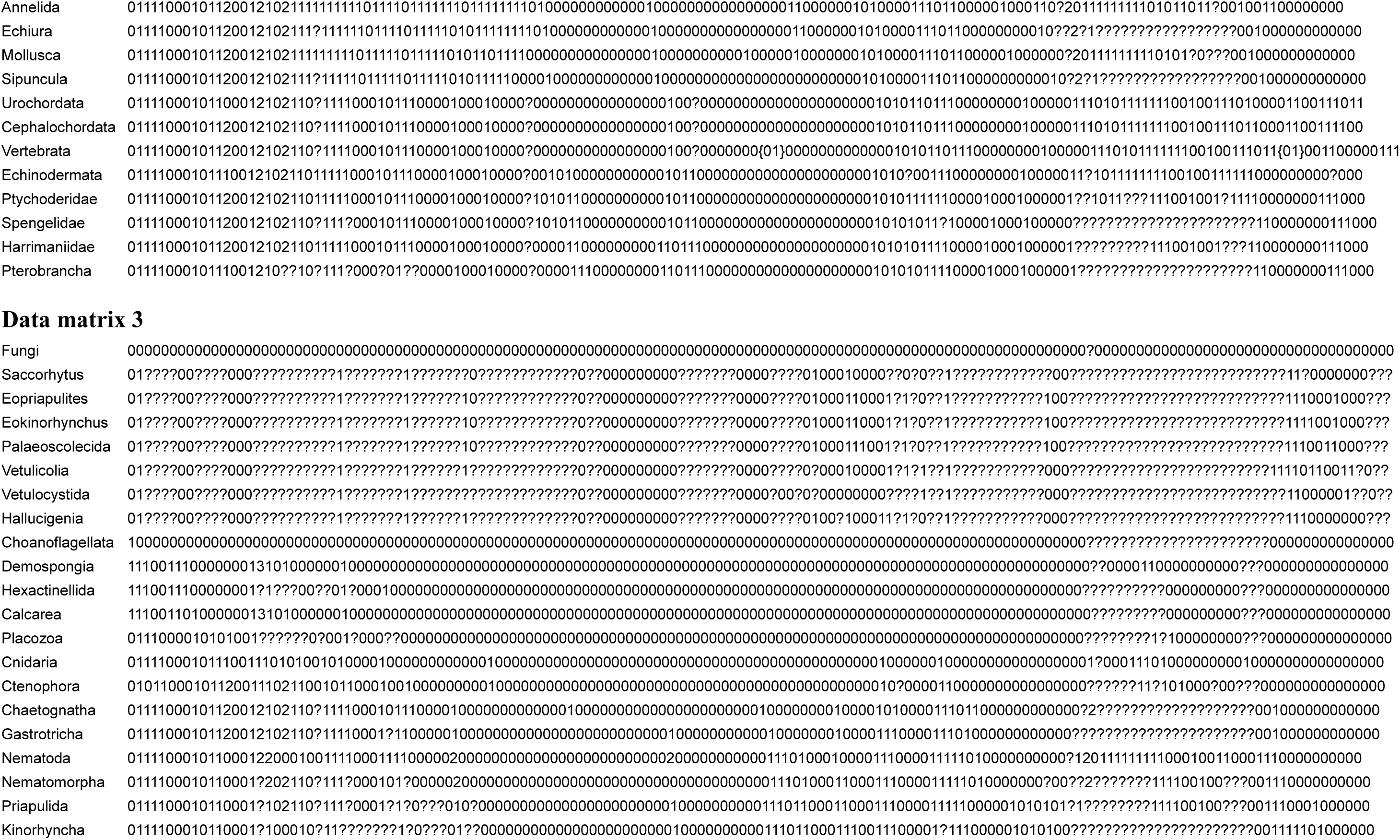
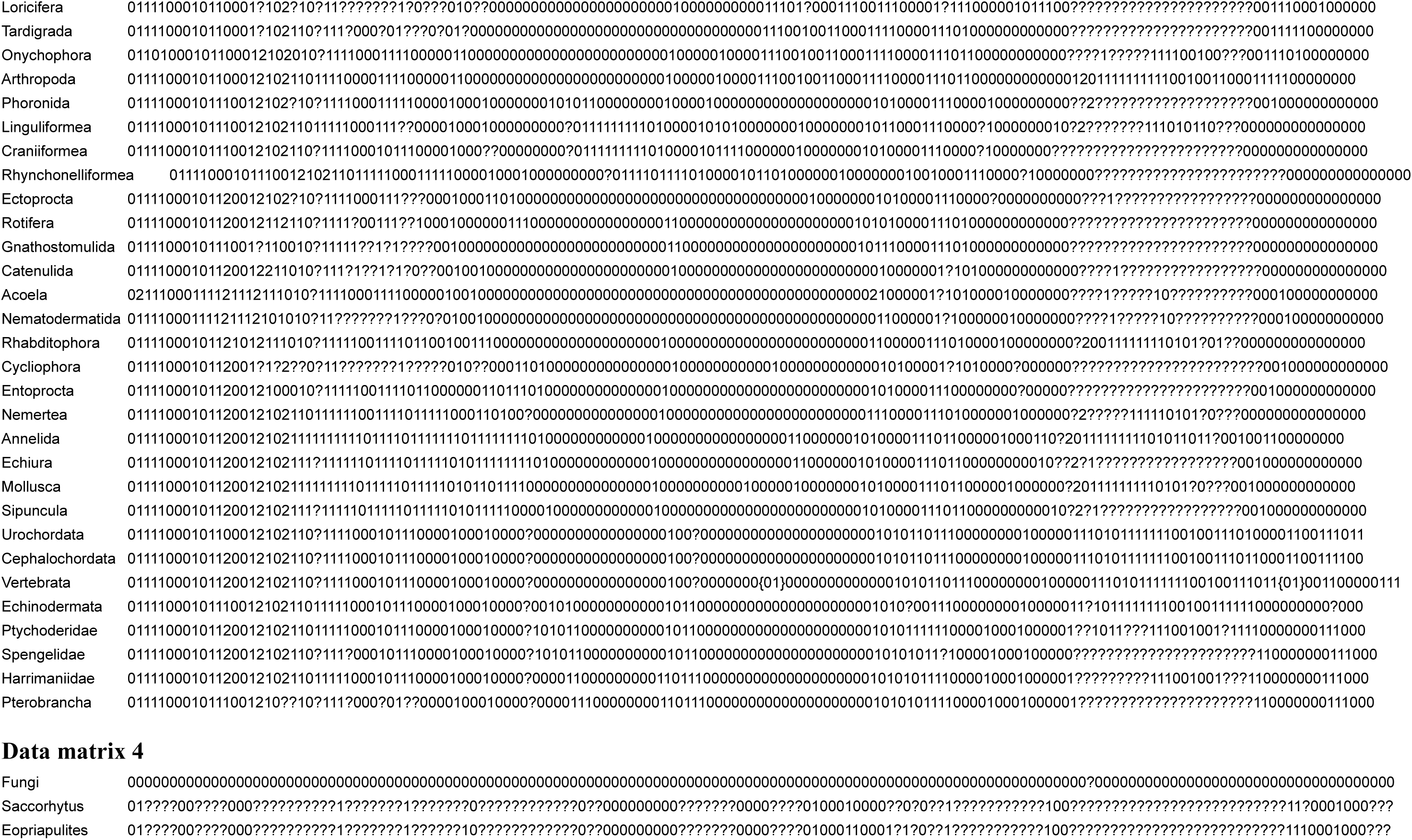
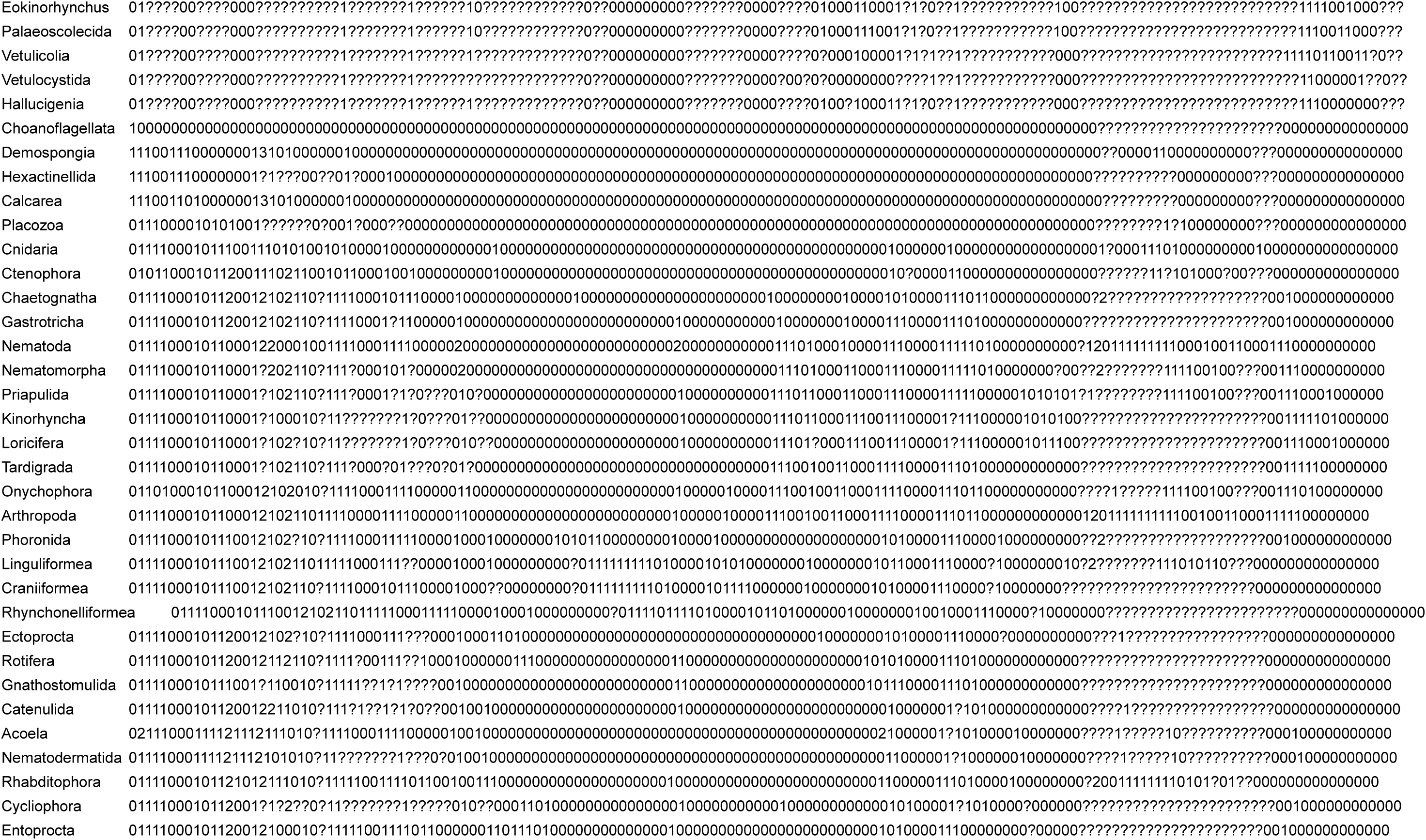
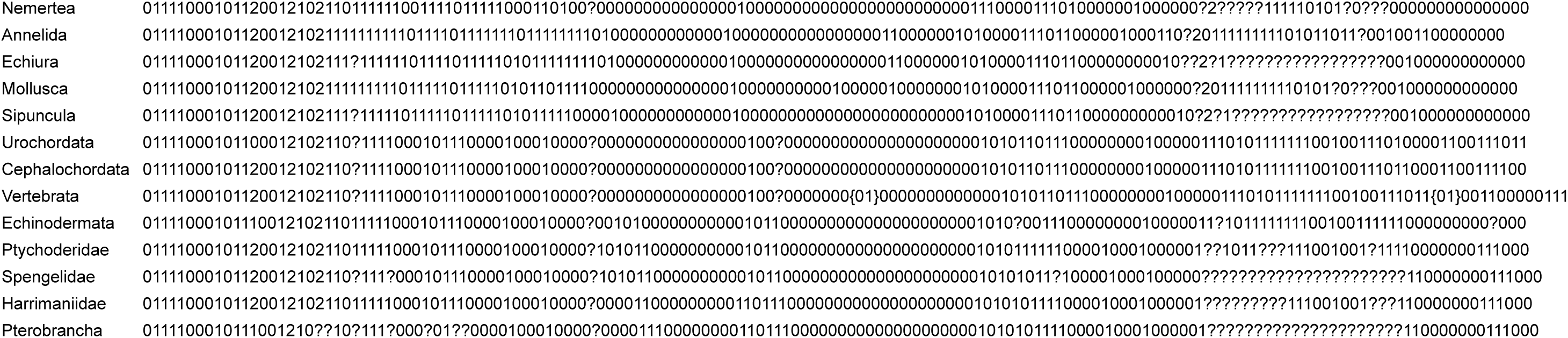

## Animations

**Supplementary Movie 1 |** SRXTM slice movie based on volume rendition of specimen UMCU2014005 (Extended Data Fig. 2a). Slices perpendicular to anterior-posterior body axis and start from anterior end.

**Supplementary Movie 2 |** SRXTM slice movie based on volume rendition of specimen UMCU2014005 (Extended Data Fig. 2a). Slices parallel to anterior-posterior body axis and start from dorsal side.

**Supplementary Movie 3 |** SRXTM movie based on volume rendition of specimen UMCU2014005 (Extended Data Fig. 2a).

**Supplementary Movie 4 |** SRXTM slice movie based on volume rendition of specimen UMCU2014001 (Extended Data Fig. 2d). Slices parallel to anterior-posterior body axis and start from antero-ventral side.

**Supplementary Movie 5 |** SRXTM slice movie based on volume rendition of specimen UMCU2014001 (Extended Data Fig. 2d). Slices parallel to anterior-posterior body axis and start from postero-dorsal side.

**Supplementary Movie 6 |** SRXTM movie based on volume rendition of specimen UMCU2014001 (Extended Data Fig. 2d).

**Supplementary Movie 7 |** SRXTM slice movie based on volume rendition of specimen UMCU2014002 (Extended Data Fig. 3d). Slices parallel to anterior-posterior body axis and start from left side.

**Supplementary Movie 8 |** SRXTM slice movie based on volume rendition of specimen UMCU2014002 (Extended Data Fig. 3d). Slices parallel to anterior-posterior body axis and start from dorsal side.

**Supplementary Movie 9 |** SRXTM movie based on volume rendition of specimen UMCU2014002 (Extended Data Fig. 3d).

**Supplementary Movie 10 |** 3D reconstruction animation showing general morphology of *Saccorhytus coronarius*.

---

**Table 1.**
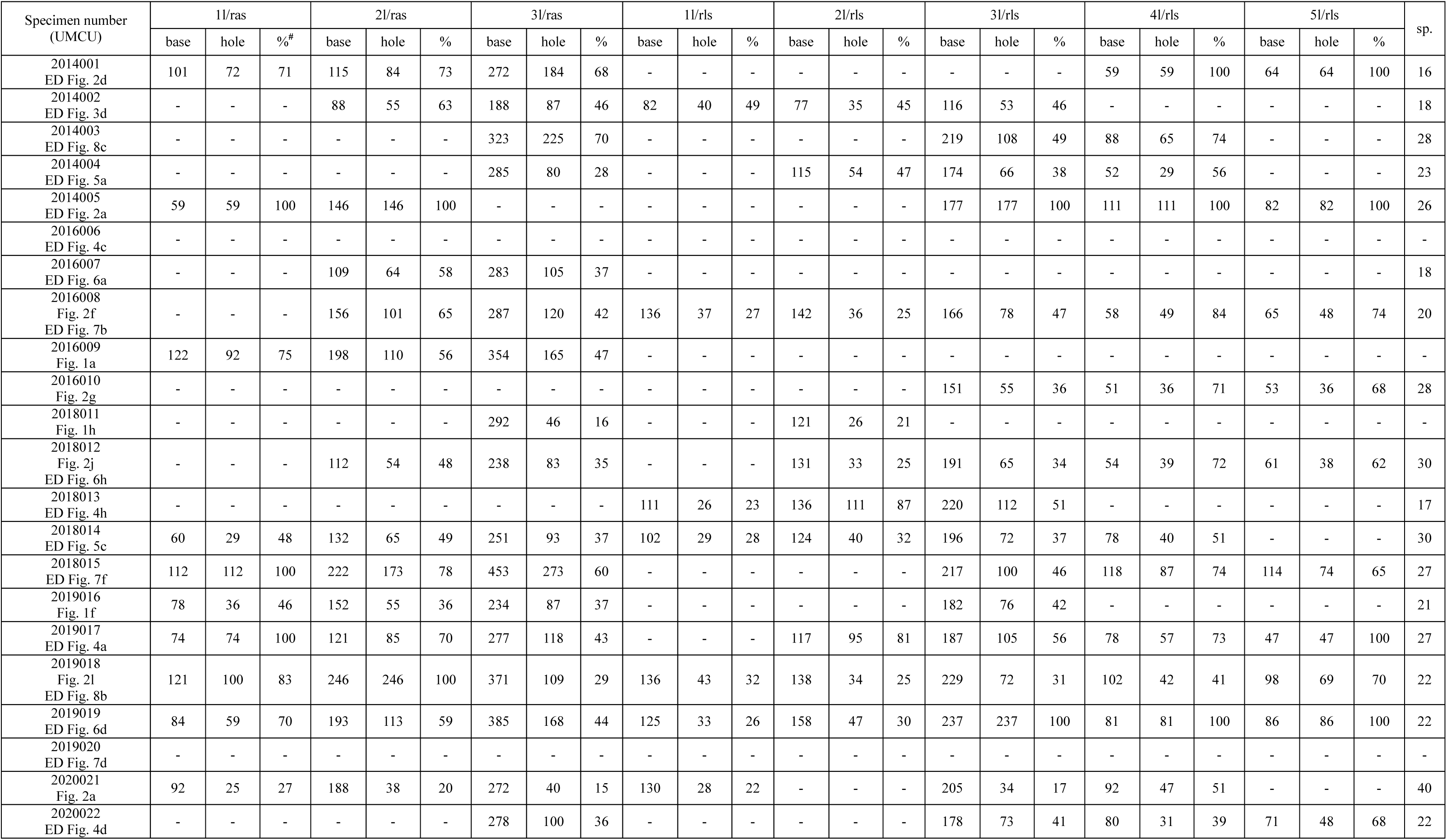
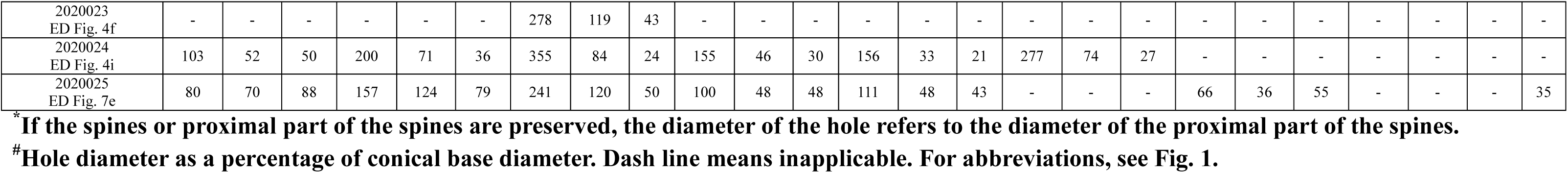
Measurements of spinose sclerites and posterior spines (μm)*

**Table 2.** Taxa and codings revised from ref^8^. Codings marked in red are revised codings. See ref^8^ for detailed description of characters.

| Character number | 1 | 2 | 3 | 4 | 5 | 6 | 7 | 8 | 9 | 10 | 11 | 12 | 13 | 14 | 15 | 16 | 17 | 18 | 19 | 20 | 21 |
| --- | --- | --- | --- | --- | --- | --- | --- | --- | --- | --- | --- | --- | --- | --- | --- | --- | --- | --- | --- | --- | --- |
| Character description | Cleavage | Fate of blastopore | Bilateral symmetry | Coelom | Coelom formation | Tripartite coelom | Protonephridia | Body division | Position of mouth | Mouth surrounded by oral folds or plates | Multi-circllets of oral folds | Introvert | Scalids | Anus | Position of anus | Segmentation or metamerism | Body divided into 13 segments | Terminal addition during ontogeny | Anterior segments fused and condensed into a head region with sensory or feeding organs | Anterior body with fused dorsal and ventral marginal zones | Shape of anterior body region |
| Cnidaria | 1 | 1 | 0 | 0 | - | 0 | 0 | 0 | ? | 0 | 0 | 0 | 0 | 0 | - | 0 | - | 0 | 0 | 0 | - |
| Acoelomorpha | 0 | 1 | 1 | 0 | 0 | - | 0 | 0 | 0 | 1 | 0 | 0 | 0 | 0 | - | 0 | - | - | - | - | - |
| Arthropoda | - | 1 | 1 | 1 | 0 | 0 | 1 | 2 | 2 | 0 | 0 | 0 | 0 | 1 | 1 | 1 | 0 | 1 | 1 | - | - |
| Onychophora | - | 1 | 1 | 1 | 0 | 0 | 1 | 0 | [12] | 1 | 0 | 0 | 0 | 1 | 1 | 1 | 0 | 1 | 1 | - | - |
| Kinorhyncha | ? | 1 | 1 | 0 | - | 0 | 1 | 0 | 2 | 1 | 0 | 1 | 1 | 1 | 1 | 1 | 0 | 0 | 0 | - | - |
| Priapulida | 1 | 1 | 1 | [01] | - | 0 | 1 | 0 | 2 | 1 | 0 | 1 | 1 | 1 | 1 | 0 | - | 1 | 0 | - | - |
| Mollusca | 0 | [01] | 1 | 1 | 0 | 0 | 1 | 0 | 1 | 0 | 0 | 0 | 0 | 1 | 1 | [01] | 0 | 1 | 0 | - | - |
| Annelida | 0 | [01] | 1 | 1 | 0 | 0 | 1 | 0 | 1 | 0 | 0 | 0 | 0 | 1 | 1 | 1 | 0 | 1 | 0 | - | - |
| Echinodermata | 1 | 0 | 1 | 1 | 1 | 1 | 1 | 2 | [02] | 0 | 0 | 0 | 0 | 1 | [01] | 0 | - | 0 | - | - | - |
| Enteropneusta | 1 | 0 | 1 | 1 | 1 | 1 | 1 | 2 | 1 | 0 | 0 | 0 | 0 | 1 | 1 | 0 | - | 0 | - | - | - |
| Urochordata | - | 0 | 1 | 1 | 1 | 0 | 1 | 1 | ? | 0 | 0 | 0 | 0 | 1 | 0 | ? | - | 0 | 0 | - | - |
| Cephalochordata | 1 | 0 | 1 | 1 | 1 | 0 | 1 | 1 | 1 | 0 | 0 | 0 | 0 | 1 | 0 | 1 | - | 0 | 0 | - | - |
| Vertebrata | 1 | 0 | 1 | 1 | 1 | 0 | 1 | 0 | 1 | 0 | 0 | 0 | 0 | 1 | 0 | 1 | - | 0 | 0 | - | - |
| *Vetulicola* | ? | ? | 1 | 1 | ? | ? | ? | 1 | 2 | 0 | 2 | 0 | 0 | 1 | 1 | 1 | 1 | 0 | 0 | 1 | 1 |
| *Beidazoon* | ? | ? | 1 | 1 | ? | ? | ? | 1 | 2 | 0 | 2 | 0 | 0 | 1 | 1 | 1 | 1 | 0 | 0 | 1 | 1 |
| *Ooedigera* | ? | ? | 1 | 1 | ? | ? | ? | 1 | 2 | 0 | 2 | 0 | 0 | 1 | 1 | 1 | 1 | 0 | 0 | 1 | 0 |
| *Pomatrum* | ? | ? | 1 | 1 | ? | ? | ? | 1 | 2 | 1 | 1 | 0 | 0 | 1 | 1 | 1 | 1 | 0 | 0 | 1 | 0 |
| *Xidazoon* | ? | ? | 1 | 1 | ? | ? | ? | 1 | 2 | 1 | 1 | 0 | 0 | 1 | 1 | 1 | 1 | 0 | 0 | 1 | 0 |
| *Didazoon* | ? | ? | 1 | 1 | ? | ? | ? | 1 | 2 | 1 | 2 | 0 | 0 | 1 | 1 | 1 | 1 | 0 | 0 | 1 | 0 |
| *Yuyuanozoon* | ? | ? | 1 | 1 | ? | ? | ? | 1 | 2 | 0 | 2 | 0 | 0 | 1 | 1 | 1 | 1 | 0 | 0 | 1 | 0 |
| *Heteromorphus* | ? | ? | 1 | 1 | ? | ? | ? | 1 | 2 | 1 | 2 | 0 | 0 | 1 | 1 | 1 | 0 | 0 | 0 | 1 | 0 |
| *Banffia* | ? | ? | 1 | 1 | ? | ? | ? | 1 | 2 | 1 | 2 | 0 | 0 | 1 | 1 | 1 | 0 | 0 | 0 | 1 | 0 |
| *Vetulocystis* | ? | ? | 1 | ? | ? | ? | ? | 1 | ? | ? | ? | 0 | 0 | 1 | 1 | 1 | 0 | 0 | 0 | 0 | - |
| *Dianchicystis* | ? | ? | 1 | ? | ? | ? | ? | 1 | ? | ? | ? | 0 | 0 | 1 | 1 | 1 | 0 | 0 | 0 | 0 | - |
| *Saccorhytus* (original of ref<sup>8</sup>) | ? | ? | 1 | ? | ? | ? | ? | 0 | 0 | 1 | 1 | 0 | 0 | 0 | - | 0 | - | 0 | 0 | 0 | - |
| *Saccorhytus* (revised) | ? | ? | 1 | ? | ? | ? | ? | 0 | 2 | 1 | 0 | 0 | 0 | 0 | - | 0 | - | 0 | 0 | 0 | - |

**Table 2 (continued). Taxa and codings revised from ref<sup>8</sup>. Codings marked in red are revised codings. See ref<sup>8</sup> for detailed description of characters.**
| Character number | 22 | 23 | 24 | 25 | 26 | 27 | 28 | 29 | 30 | 31 | 32 | 33 | 34 | 35 | 36 | 37 | 38 | 39 | 40 | 41 |
| --- | --- | --- | --- | --- | --- | --- | --- | --- | --- | --- | --- | --- | --- | --- | --- | --- | --- | --- | --- | --- |
| Character description | Shape of pharyngeal region | Posterior body with numerous narrow segments/annulations | Endoskeleton mesodermally derived | Surface membrane (epidermis) | Surface cuticle (or exoskeleton) underlain by epidermis | Cuticle mainly made of α-chitin | Growth by ecdysis (moulting) | Three-layered cuticle | Triannate epicuticle | Circumpharyngeal brain | Chaetae | Diffuse nerve net | Longitudinal ventral nerve cord(s) | Longitudinal dorsal nerve cord(s) | Longitudinal lateral grooves (median zones) | Pharyngeal gill slits | Body cones | Body cones with radial folds | Orientation of body cones | Number of body cones |
| Cnidaria | - | 0 | 0 | 0 | 0 | 0 | 0 | 0 | 0 | 0 | 0 | 1 | 0 | 0 | 0 | 0 | 0 | - | - | - |
| Acoelomorpha | - | - | 0 | 0 | 0 | - | - | 0 | 0 | 0 | 0 | 1 | 0 | 0 | 0 | 0 | 0 | - | - | - |
| Arthropoda | - | - | 0 | 0 | 1 | 1 | 1 | 1 | 1 | 0 | 1 | 0 | 1 | 0 | 0 | 0 | 0 | - | - | - |
| Onychophora | - | - | 0 | 0 | 1 | 1 | 1 | 1 | 1 | 0 | 1 | 0 | 1 | 0 | 0 | 0 | 0 | - | - | - |
| Kinorhyncha | - | - | 0 | 0 | 1 | 0 | 1 | 1 | 1 | 0 | 1 | 0 | 1 | 0 | 0 | 0 | 0 | - | - | - |
| Priapulida | - | - | 0 | 0 | 1 | 0 | 1 | 1 | 1 | 0 | 1 | 0 | 1 | 0 | 0 | 0 | 0 | - | - | - |
| Mollusca | - | - | 0 | 0 | 0 | 0 | 0 | 0 | 0 | 1 | 0 | 0 | 1 | 0 | 0 | 0 | 0 | - | - | - |
| Annelida | - | - | 0 | 0 | 1 | 0 | 0 | 0 | 0 | 1 | 0 | 0 | 1 | 0 | 0 | 0 | 0 | - | - | - |
| Echinodermata | [01] | - | 1 | 1 | 0 | - | 0 | 0 | 0 | 0 | 0 | 0 | 0 | 0 | 0 | [01] | 0 | - | - | - |
| Enteropneusta | - | - | 1 | 1 | 0 | - | 0 | 0 | 0 | 0 | 0 | 0 | 1 | 1 | 0 | 1 | 0 | - | - | - |
| Urochordata | - | - | 0 | 0 | 0 | - | 0 | 0 | 0 | 0 | 0 | 0 | 0 | 1 | 0 | 1 | 0 | - | - | - |
| Cephalochordata | - | - | 1 | 1 | 0 | - | 0 | 0 | 0 | 0 | 0 | 0 | 0 | 1 | 0 | 1 | 0 | - | - | - |
| Vertebrata | - | - | 1 | 1 | 0 | - | 0 | 0 | 0 | 0 | 0 | 0 | 0 | 1 | 0 | 1 | 0 | - | - | - |
| *Vetulicola* | 0 | 0 | 1 | 1 | 0 | ? | 0 | - | - | 0 | ? | 0 | ? | 0 | 1 | 1 | 1 | 0 | 1 | 0 |
| *Beidazoon* | 0 | 0 | 1 | 1 | 0 | ? | 0 | - | - | 0 | ? | 0 | ? | 0 | 1 | 1 | 1 | 0 | 1 | 0 |
| *Ooedigera* | 0 | 0 | 1 | 1 | 0 | ? | 0 | - | - | 0 | ? | 0 | ? | 0 | 1 | 1 | 1 | 0 | 1 | 0 |
| *Pomatrum* | 0 | 0 | 1 | 1 | 0 | ? | 0 | - | - | 0 | ? | 0 | ? | 0 | 1 | 1 | 1 | 0 | 1 | 0 |
| *Xidazoon* | 0 | 0 | 1 | 1 | 0 | ? | 0 | - | - | 0 | ? | 0 | 1 | 0 | 1 | 1 | 1 | 0 | 1 | 0 |
| *Didazoon* | 0 | 0 | 1 | 1 | 0 | ? | 0 | - | - | 0 | ? | 0 | ? | 0 | 1 | 1 | 1 | 0 | 1 | 0 |
| *Yuyuanozoon* | 0 | 0 | 1 | 1 | 0 | ? | 0 | - | - | 0 | ? | 0 | ? | 0 | 1 | 1 | 1 | 0 | 1 | 0 |
| *Heteromorphus* | 0 | 1 | 1 | 1 | 0 | ? | 0 | - | - | 0 | ? | 0 | ? | 0 | 1 | 1 | 1 | 0 | 1 | 0 |
| *Banffia* | 0 | 1 | 1 | ? | 0 | ? | 0 | - | - | 0 | ? | 0 | ? | 0 | 1 | 1 | 1 | 0 | 1 | 0 |
| *Vetulocystis* | 1 | 1 | 1 | 1 | ? | ? | 0 | - | - | 0 | ? | 0 | ? | ? | 0 | 1 | 1 | 1 | 0 | 1 |
| *Dianchicystis* | 1 | 1 | 1 | 1 | ? | ? | 0 | - | - | 0 | ? | 0 | ? | ? | 0 | 1 | 1 | 1 | 0 | 1 |
| *Saccorhytus* (original of ref<sup>8</sup>) | 1 | 0 | 1 | 1 | ? | ? | 0 | - | - | 0 | ? | ? | 0 | ? | 0 | 1 | 1 | 2 | 0 | 2 |
| *Saccorhytus* (revised) | - | 0 | ? | 1 | 0 | - | 0 | - | - | ? | 0 | ? | ? | ? | 0 | 0 | 0 | - | - | - |

**Table 2 (continued). Taxa and codings revised from ref<sup>8</sup>. Codings marked in red are revised codings. See ref<sup>8</sup> for detailed description of characters.**
| Character number | 42 | 43 | 44 | 45 | 46 | 47 | 48 | 49 | 50 | 51 | 52 | 53 | 54 | 55 | 56 | 57 | 58 | 59 | 60 | 61 |
| --- | --- | --- | --- | --- | --- | --- | --- | --- | --- | --- | --- | --- | --- | --- | --- | --- | --- | --- | --- | --- |
| Character description | Gill bars/arches | Gill cowls (hoods) | Filter feeding accomplished within pharynx | Endostyle, epibranchial ridge, or thyroid that accumulates iodine and secretes mucous | Pharyngeal food-transport groove | Peripharyngeal atrium | Notochord | X-shaped mark around the antero-posterior junction | ?Lenticulate respiratory organ | V- or W-shaped myomeres | Neural crest cells | Neurogenic placodes | Tomaria-like larva | Trochophore | Oxygen transport proteins | tRNA Lys | Lox2/4 | Abd-B duplication | Antp | HRP |
| Cnidaria | - | - | 0 | - | - | 0 | 0 | 0 | 0 | 0 | 0 | 0 | 0 | 0 | 0 | 0 | 0 | 0 | 0 | 0 |
| Acoelomorpha | - | - | 0 | - | - | 0 | 0 | - | 0 | 0 | 0 | 0 | 0 | 0 | 0 | ? | ? | ? | ? | 0 |
| Arthropoda | - | - | 0 | - | - | 0 | 0 | - | 0 | 0 | 0 | 0 | 0 | 0 | 1 | ? | 0 | 0 | 1 | 1 |
| Onychophora | - | - | 0 | - | - | 0 | 0 | - | 0 | 0 | 0 | 0 | 0 | 0 | 1 | ? | 0 | 0 | 1 | 1 |
| Kinorhyncha | - | - | 0 | - | - | 0 | 0 | - | 0 | 0 | 0 | 0 | 0 | 0 | 1 | ? | ? | ? | ? | 1 |
| Priapulida | - | - | 0 | - | - | 0 | 0 | - | 0 | 0 | 0 | 0 | 0 | 0 | 1 | ? | 0 | 0 | 1 | 1 |
| Mollusca | - | - | 0 | - | - | 0 | 0 | - | 0 | 0 | 0 | 0 | 0 | 1 | 1 | 0 | 0 | 0 | 0 | 0 |
| Annelida | - | - | 0 | - | - | 0 | 0 | - | 0 | 0 | 0 | 0 | 0 | 1 | 1 | 0 | 0 | 0 | 0 | 0 |
| Echinodermata | - | - | 0 | - | - | ? | 0 | - | 0 | 0 | 0 | 0 | 1 | 0 | 1 | 1 | 1 | 1 | 0 | 0 |
| Enteropneusta | 1 | 0 | 1 | 1 | 1 | 1 | 0 | - | 0 | 0 | 0 | 0 | 1 | 0 | 1 | 1 | 1 | 1 | 0 | 0 |
| Urochordata | 1 | 0 | 1 | 1 | 1 | 1 | 1 | 0 | 0 | 0 | 1 | 1 | 0 | 0 | 1 | 0 | 1 | 1 | 0 | 0 |
| Cephalochordata | 1 | 0 | 1 | 1 | 1 | 1 | 1 | - | 0 | 1 | 0 | 0 | 0 | 0 | 1 | 0 | 1 | 1 | 0 | 0 |
| Vertebrata | 1 | 0 | 0 | 1 | 0 | 0 | 1 | - | 0 | 1 | 1 | 1 | 0 | 0 | 1 | 0 | 1 | 1 | 0 | 0 |
| *Vetulicola* | 0 | 0 | 1 | ? | 1 | ? | 0 | 0 | 0 | 0 | ? | ? | ? | ? | ? | ? | ? | ? | ? | ? |
| *Beidazon* | 0 | 0 | 1 | ? | 1 | ? | 0 | 0 | 0 | 0 | ? | ? | ? | ? | ? | ? | ? | ? | ? | ? |
| *Ooedigera* | 0 | 0 | 1 | ? | 1 | ? | 0 | 0 | 0 | 0 | ? | ? | ? | ? | ? | ? | ? | ? | ? | ? |
| *Pomatrum* | 0 | 1 | 1 | ? | 1 | ? | 0 | 0 | 0 | 0 | ? | ? | ? | ? | ? | ? | ? | ? | ? | ? |
| *Xidazon* | 0 | 1 | 1 | ? | 1 | ? | 0 | 0 | 0 | 0 | ? | ? | ? | ? | ? | ? | ? | ? | ? | ? |
| *Didazon* | 0 | 1 | 1 | 1 | 1 | ? | 0 | 0 | 0 | 0 | ? | ? | ? | ? | ? | ? | ? | ? | ? | ? |
| *Yuyuanozoon* | 0 | 1 | 1 | ? | 1 | ? | 0 | 0 | 0 | 0 | ? | ? | ? | ? | ? | ? | ? | ? | ? | ? |
| *Heteromorphus* | 0 | 0 | 1 | ? | 1 | ? | 0 | 1 | 0 | 0 | ? | ? | ? | ? | ? | ? | ? | ? | ? | ? |
| *Banffia* | 0 | 0 | 1 | ? | 1 | ? | 0 | 1 | 0 | 0 | ? | ? | ? | ? | ? | ? | ? | ? | ? | ? |
| *Vetulocystis* | 0 | 0 | 1 | ? | ? | ? | 0 | 0 | 1 | 0 | ? | ? | ? | ? | ? | ? | ? | ? | ? | ? |
| *Dianchicystis* | 0 | 0 | 1 | ? | ? | ? | 0 | 0 | 1 | 0 | ? | ? | ? | ? | ? | ? | ? | ? | ? | ? |
| *Saccorhytus* (original of ref<sup>8</sup>) | 0 | 0 | 1 | ? | 0 | ? | 0 | - | 0 | 0 | ? | ? | 0 | 0 | ? | ? | ? | ? | ? | ? |
| *Saccorhytus* (revised) | - | - | - | ? | - | - | ? | - | 0 | ? | ? | ? | ? | ? | ? | ? | ? | ? | ? | ? |

**Table 3.**
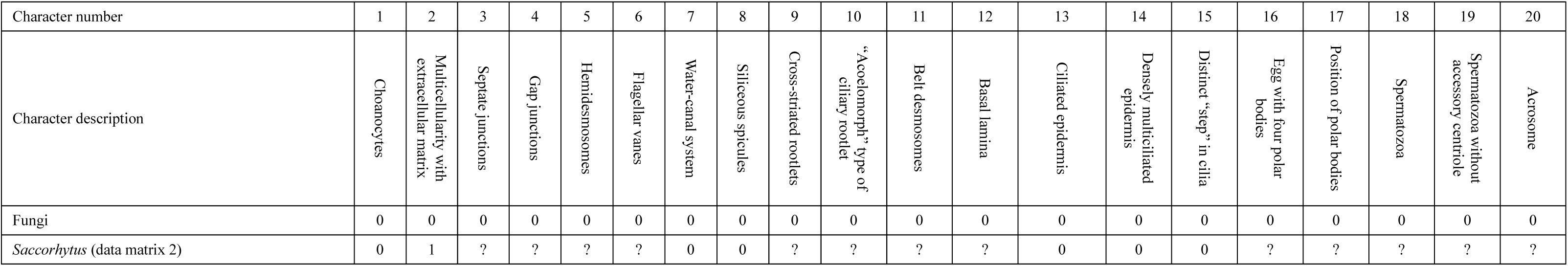
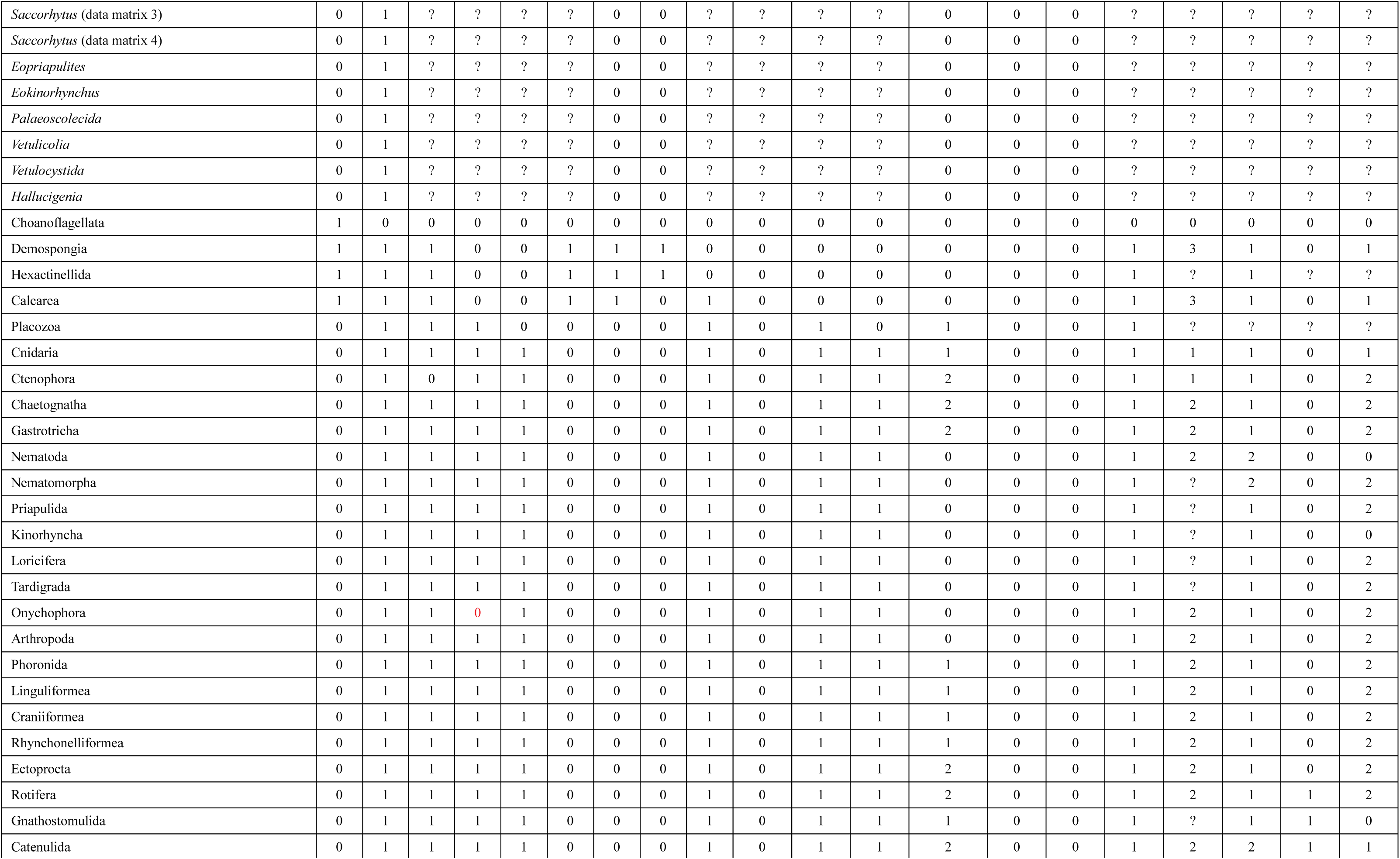
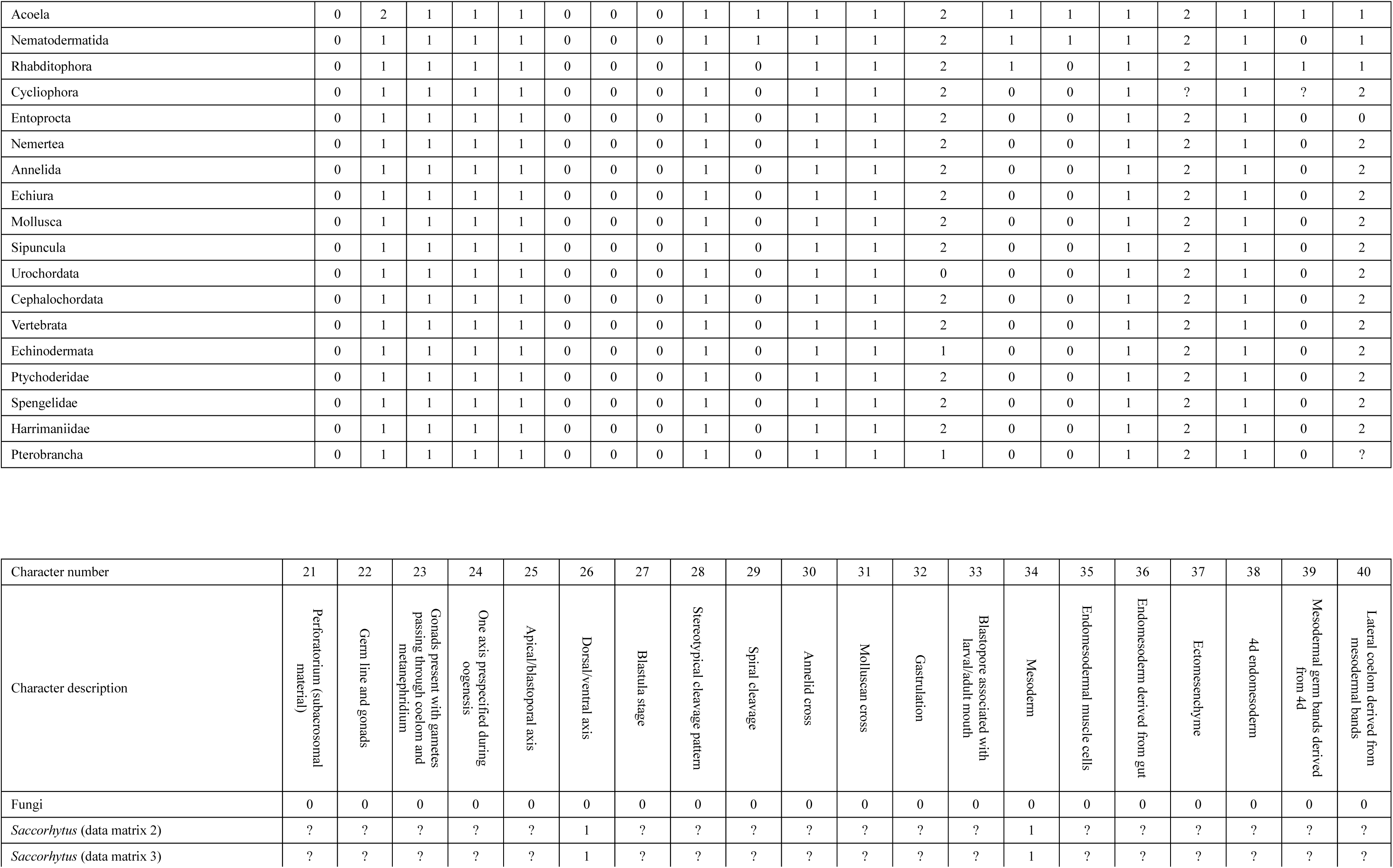
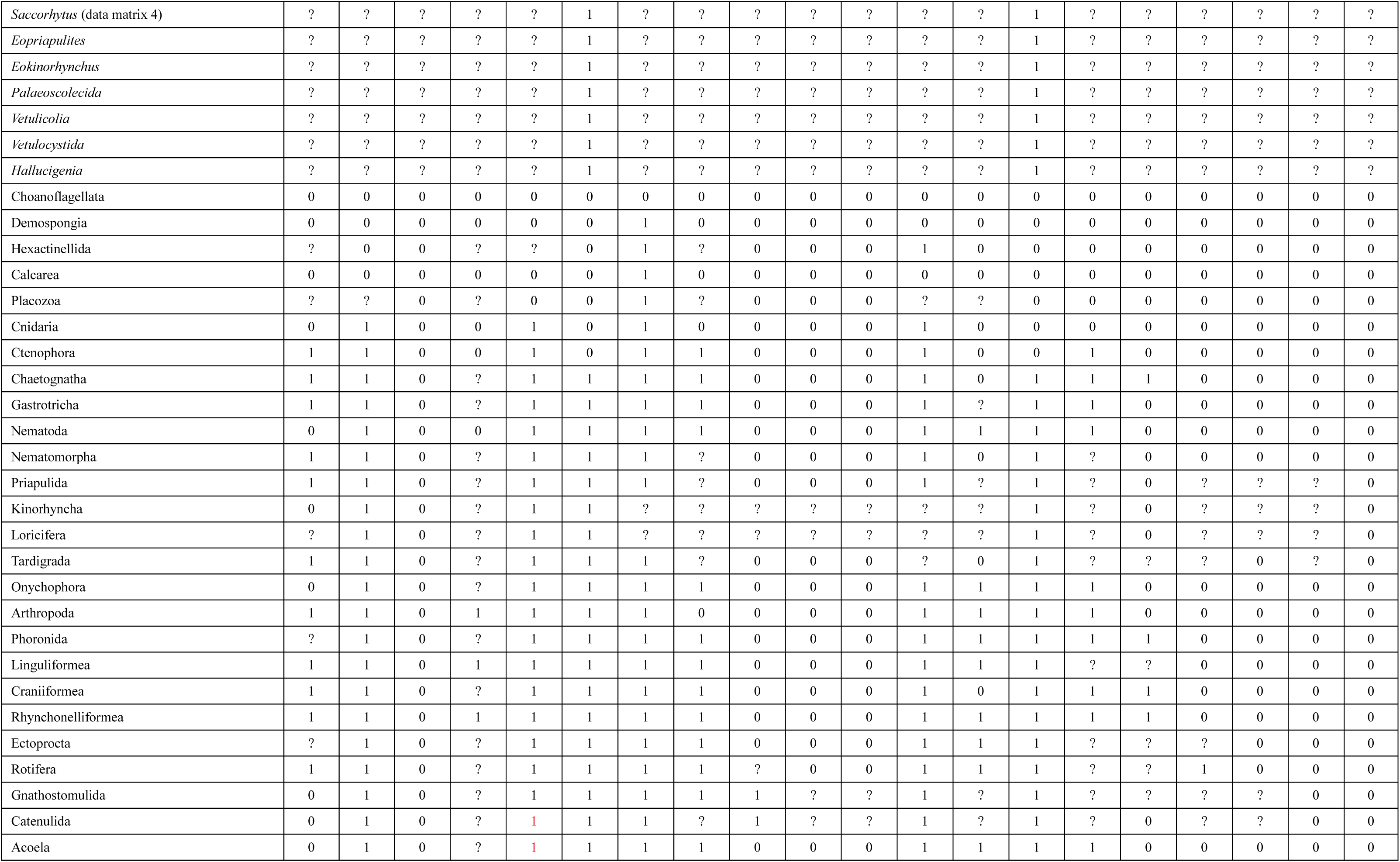
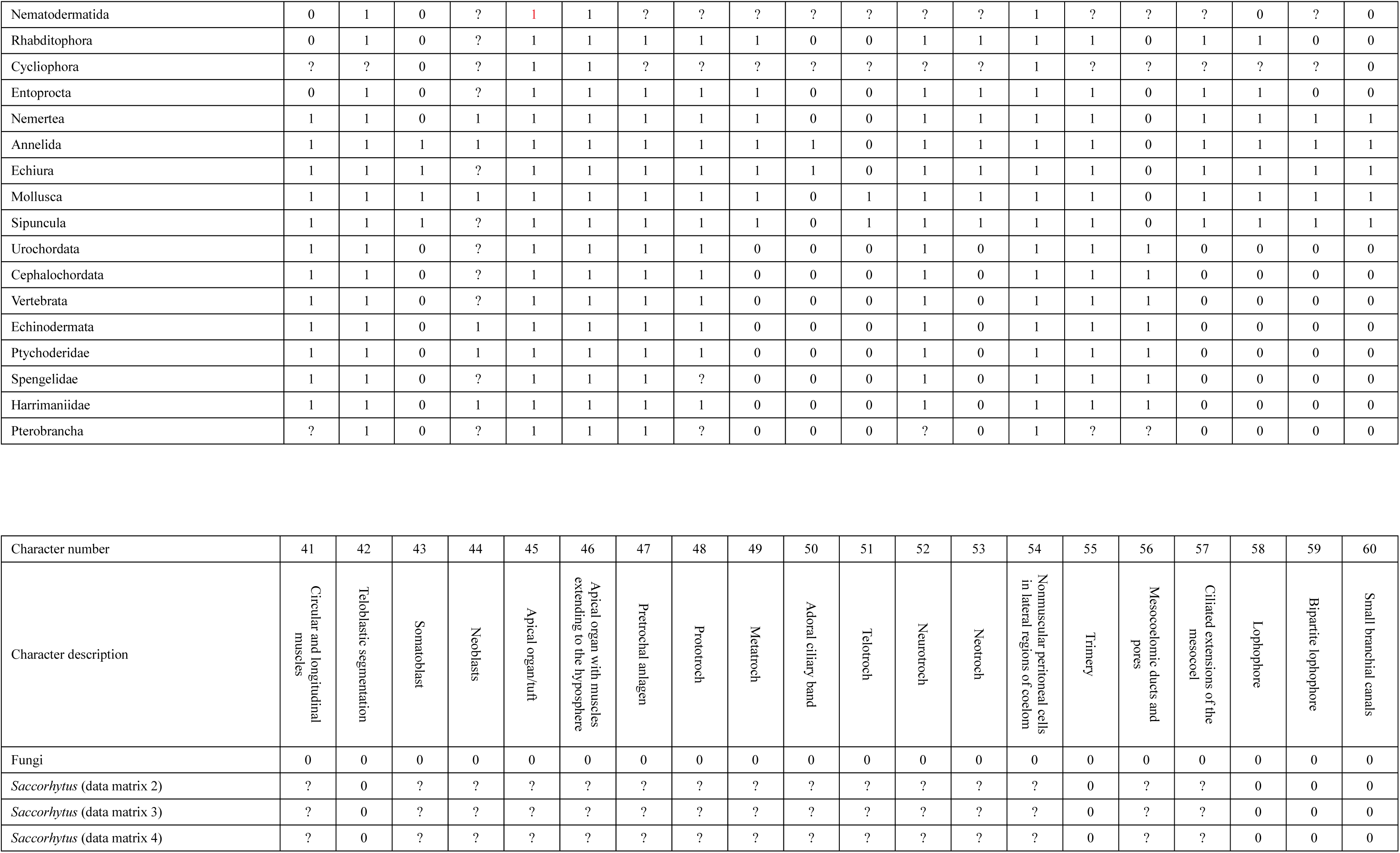
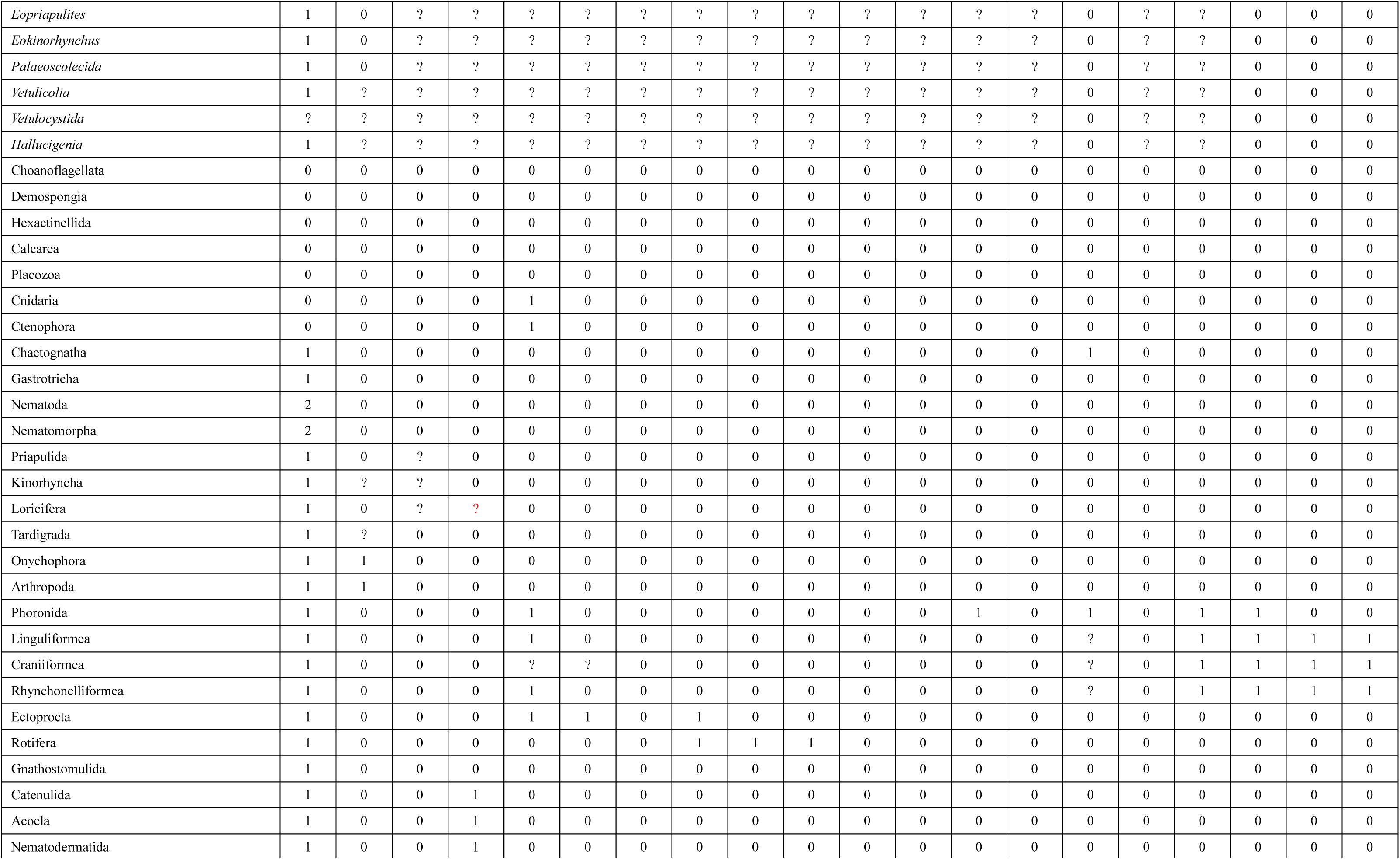
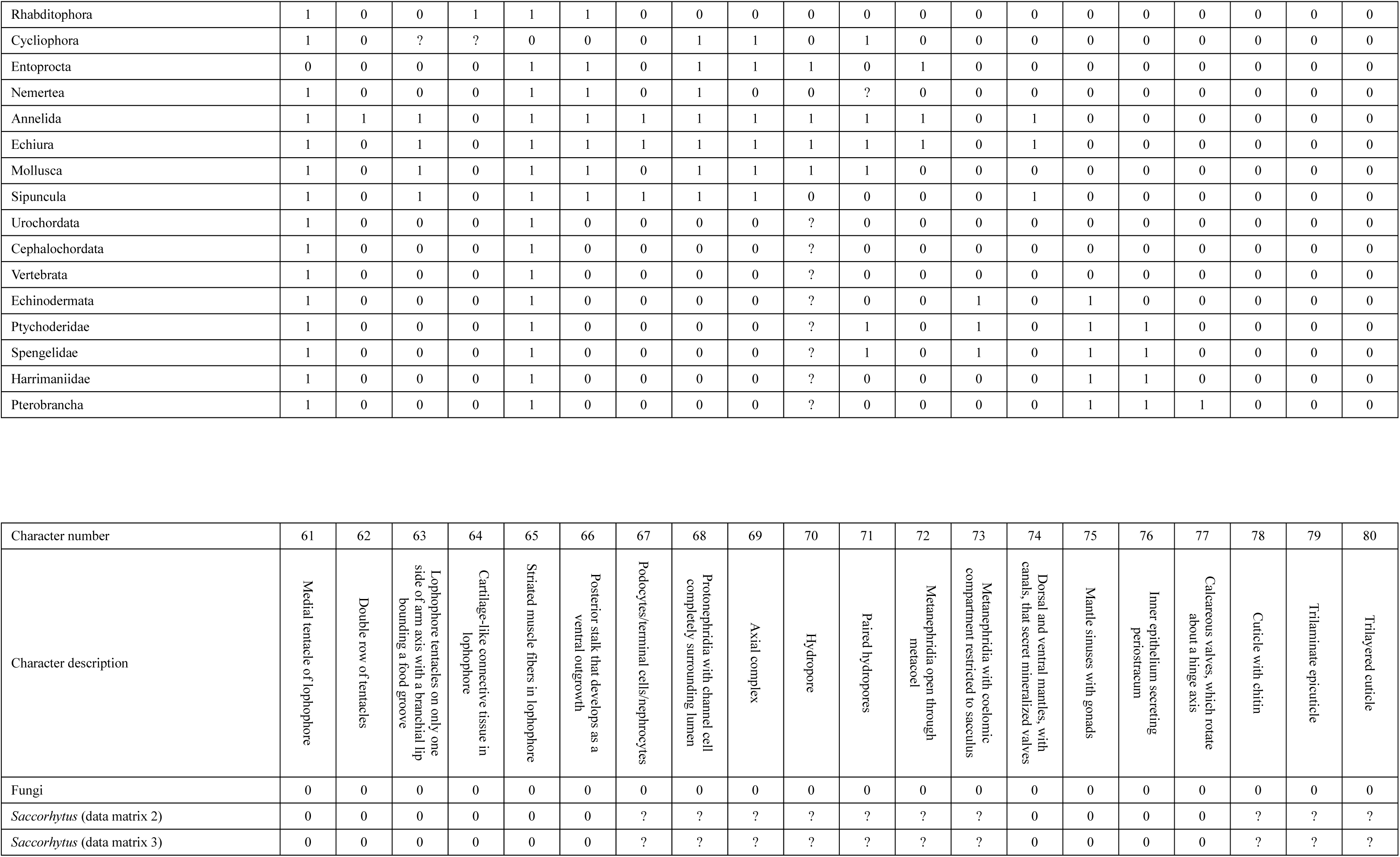
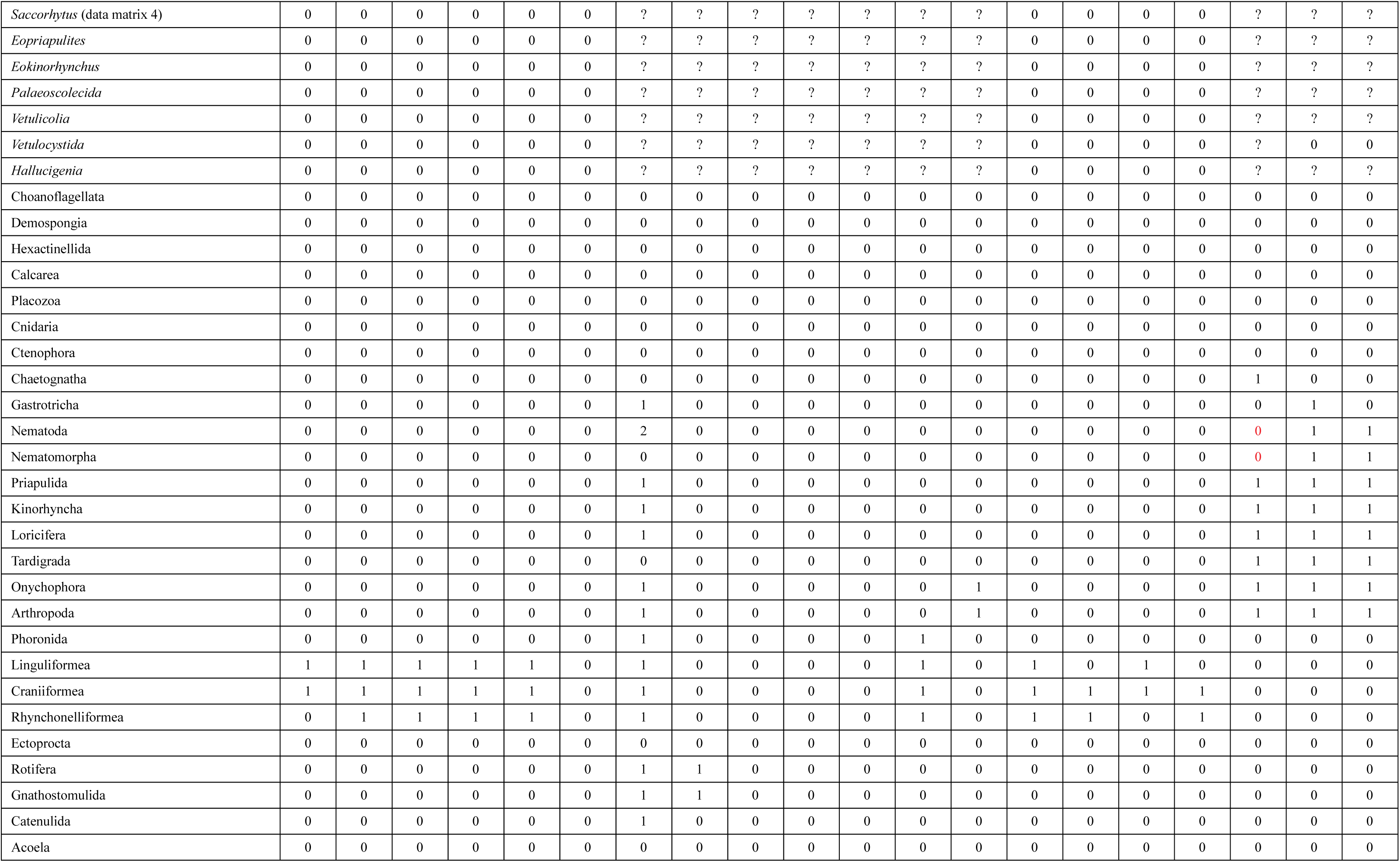
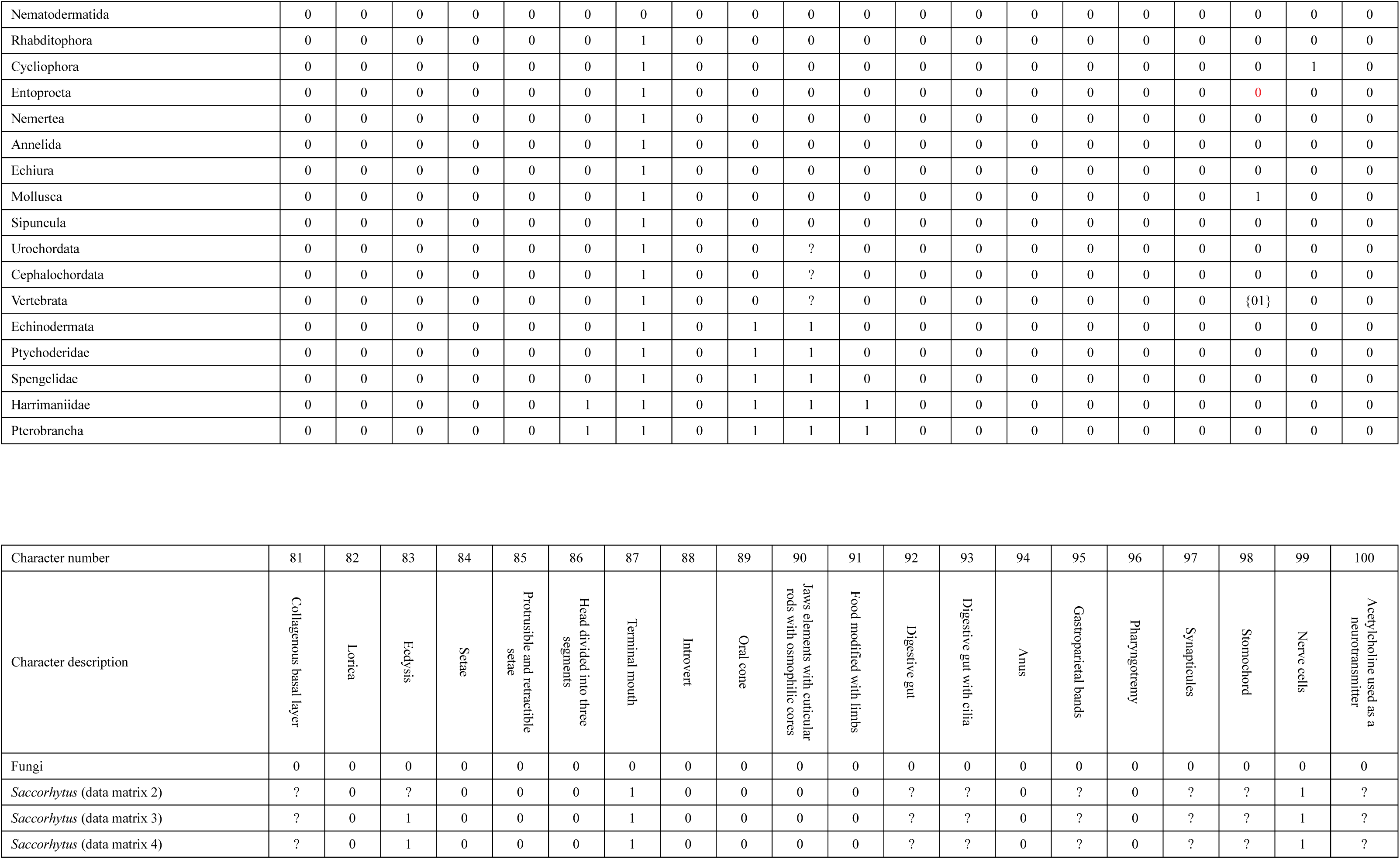
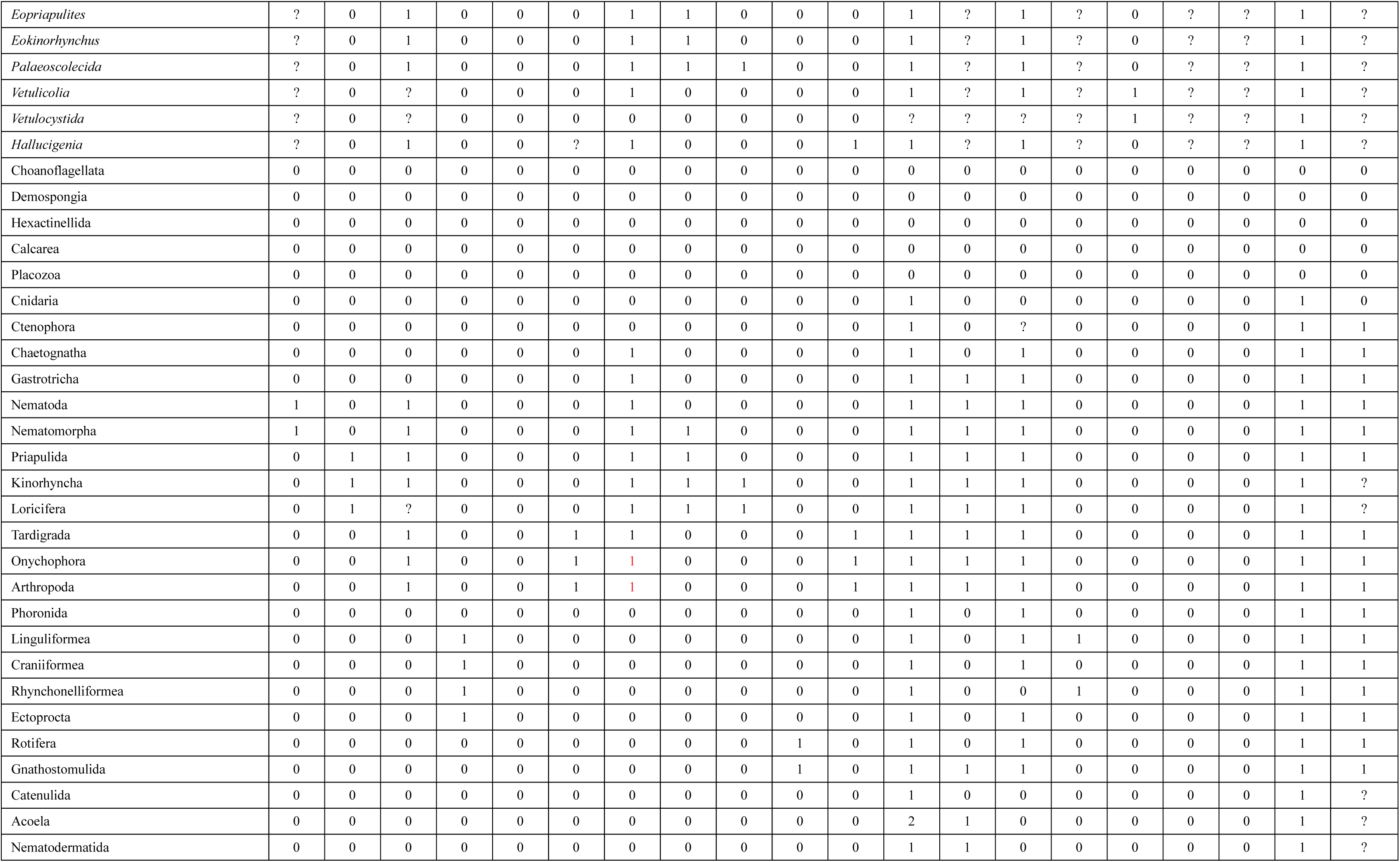
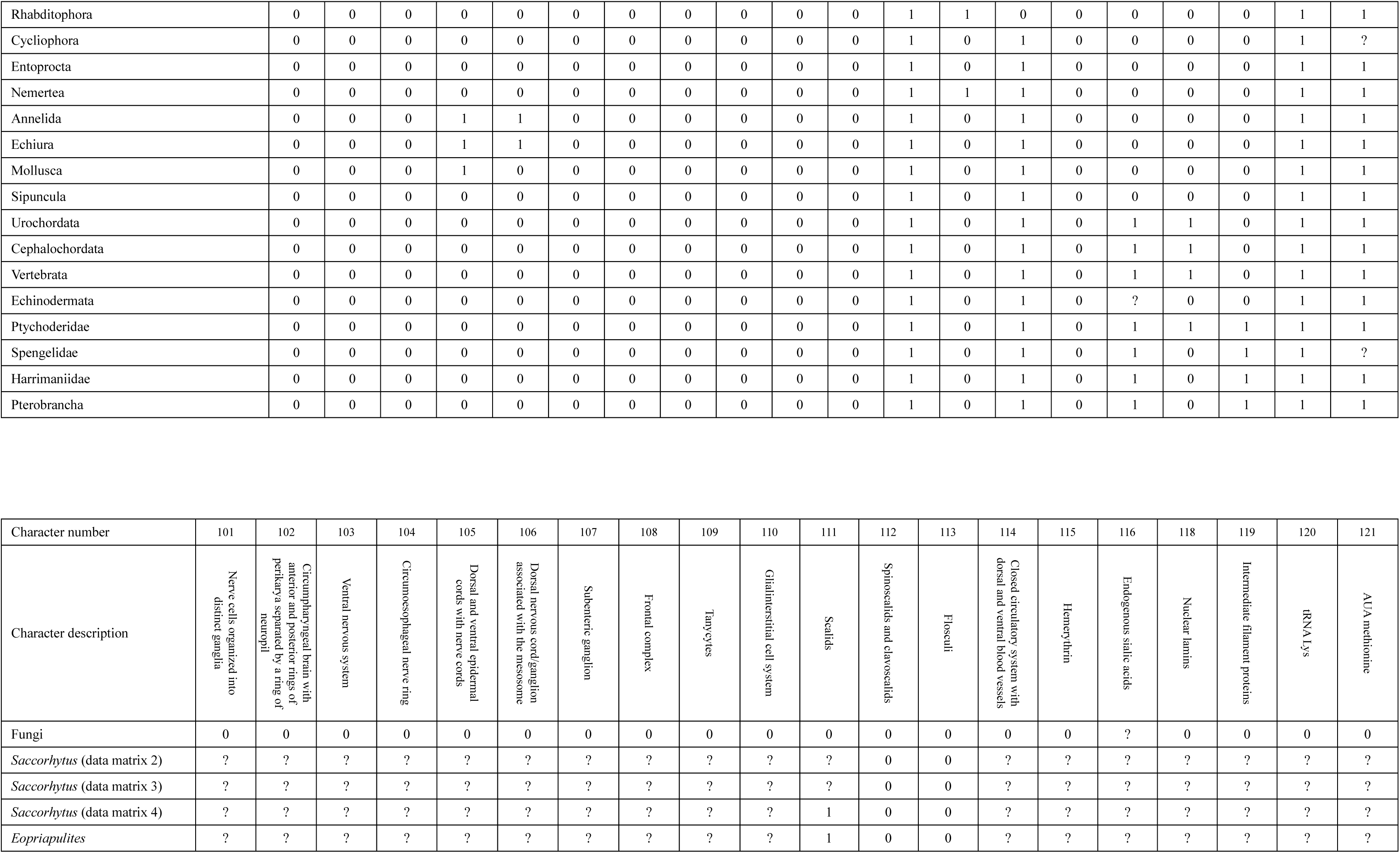
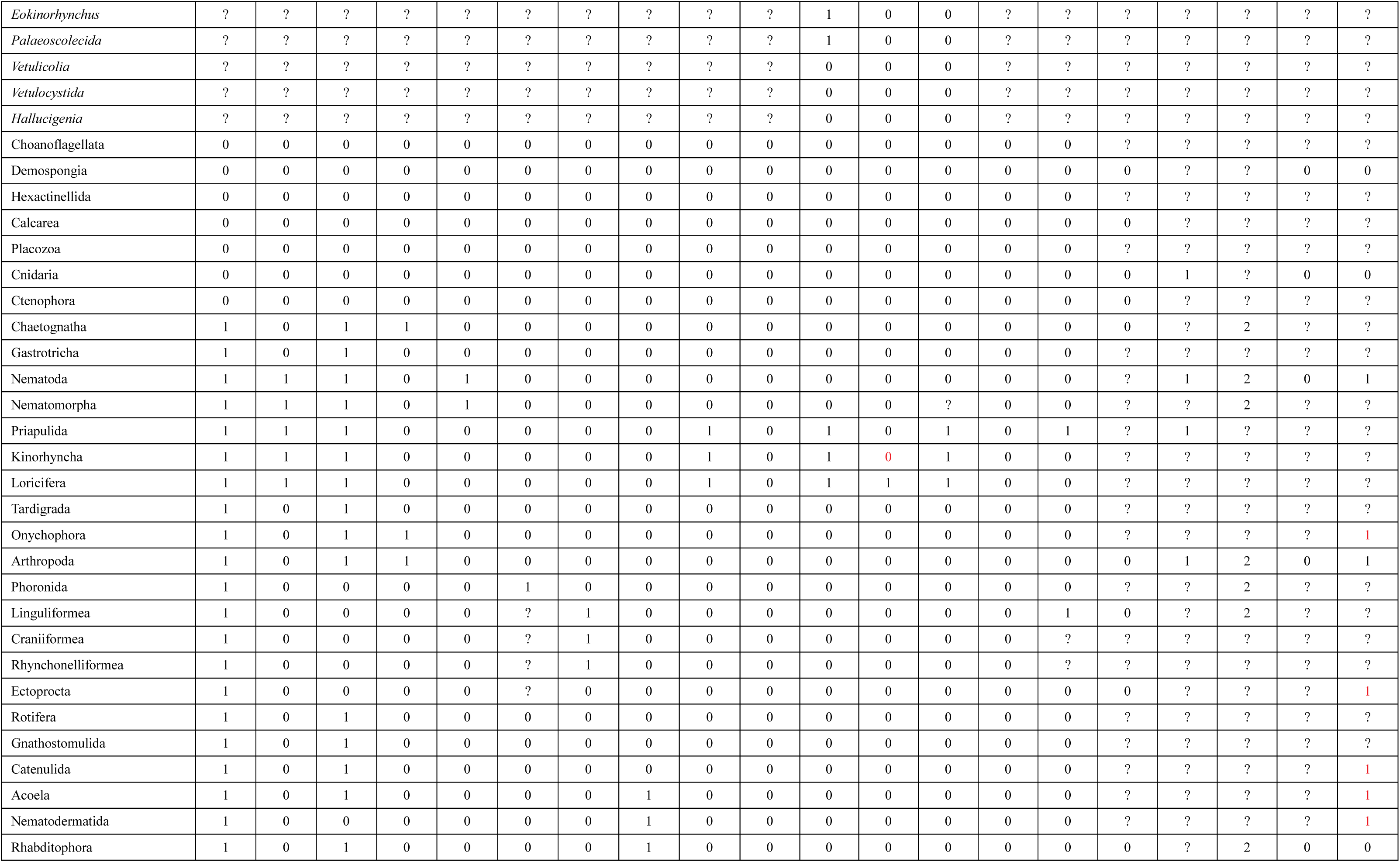
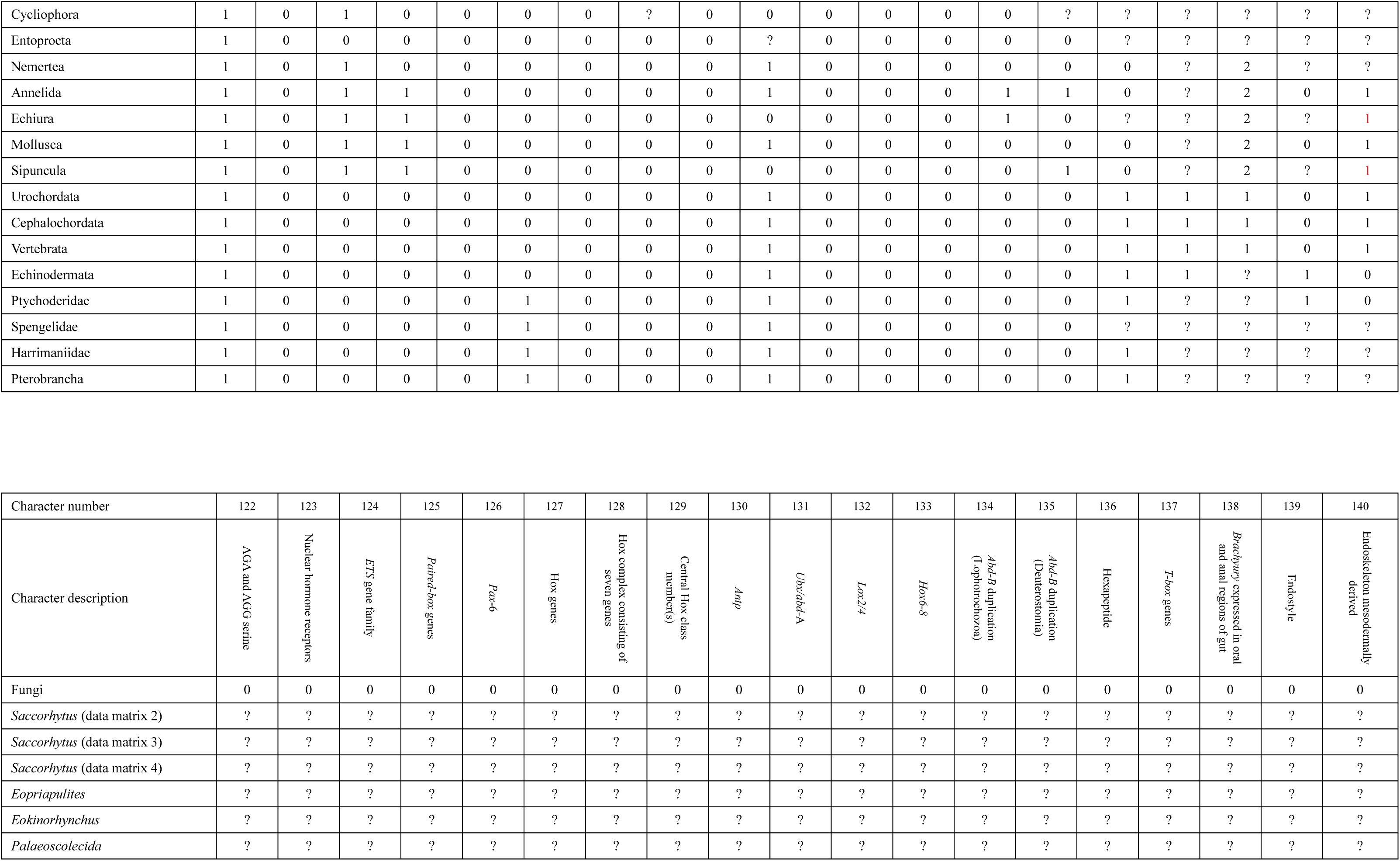
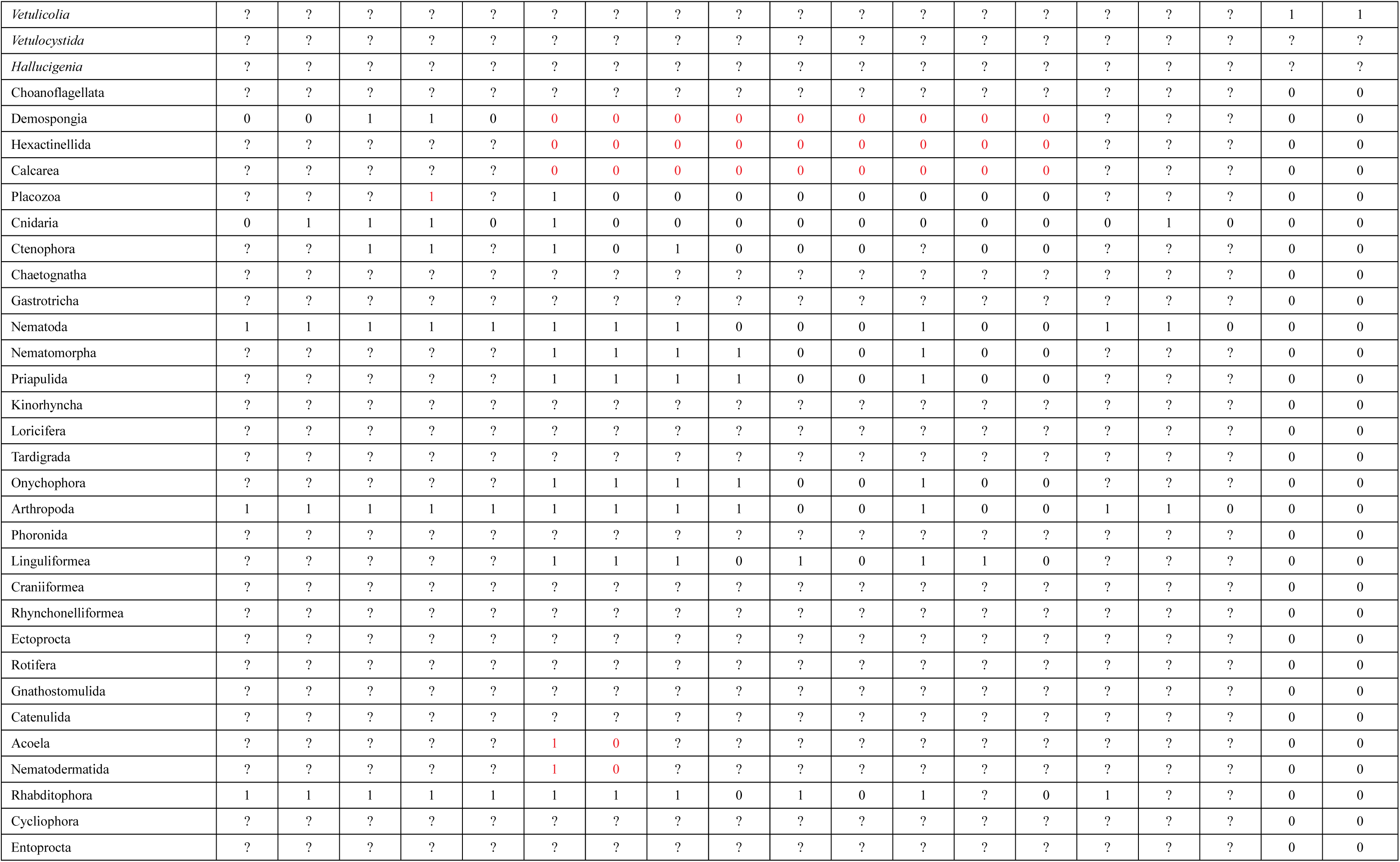
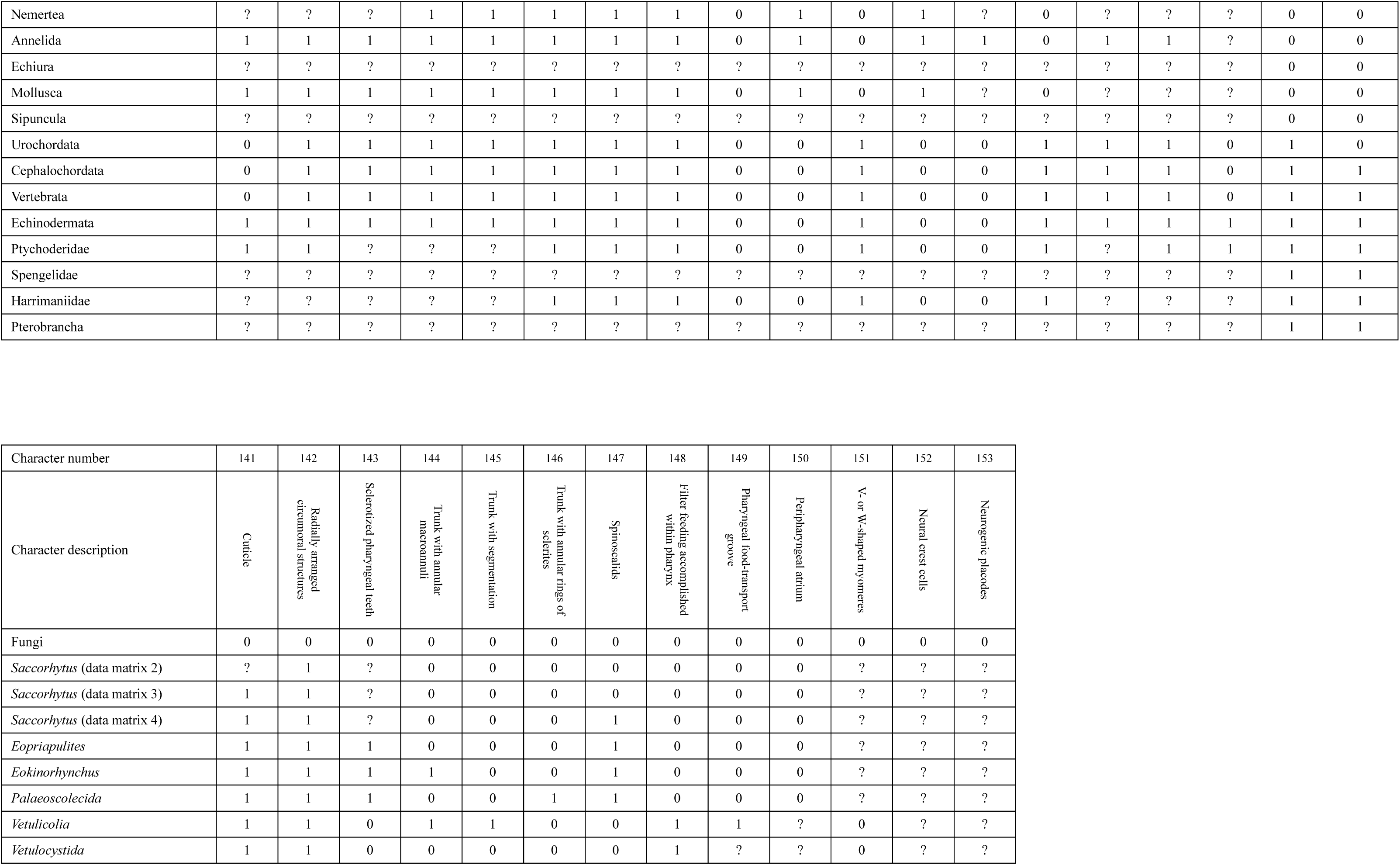
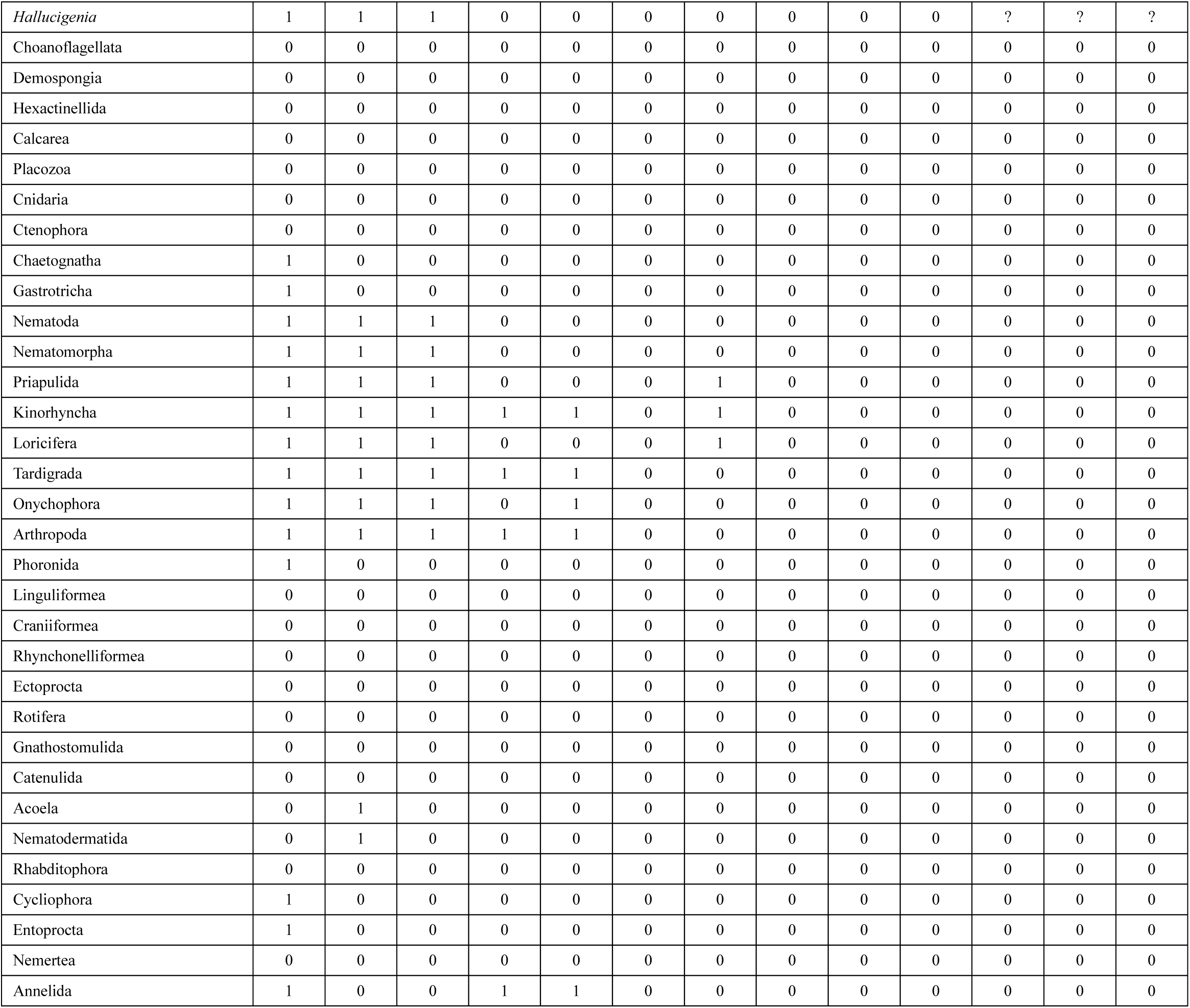
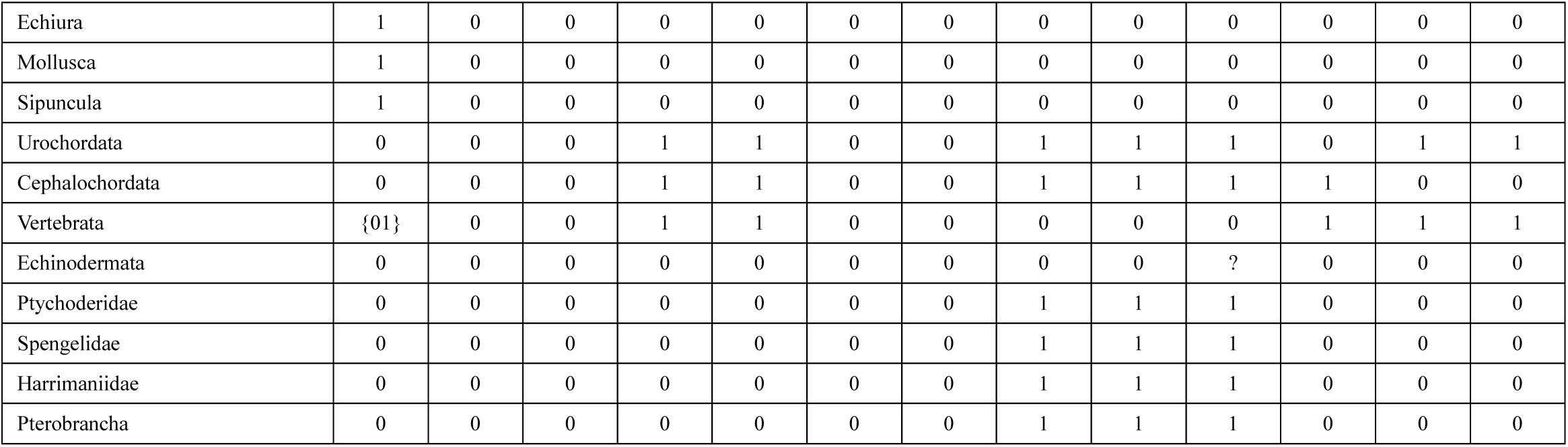
Taxa and codings modified from ref^19^. Character numbers retained those of ref^19^. Character 117 were not included in phylogenetic analysis. Revised codings are marked in red. See ref^8^ and ref^19^ for detailed description of characters.

## Notes

### Competing Interest Statement

The authors have declared no competing interest.

